# The history of chromosomal instability in genome doubled tumors

**DOI:** 10.1101/2023.10.22.563273

**Authors:** Toby M. Baker, Siqi Lai, Tom Lesluyes, Haixi Yan, Annelien Verfaillie, Stefan Dentro, Andrew R. Lynch, Amy L. Bowes, Nischalan Pillay, Adrienne M. Flanagan, Charles Swanton, Maxime Tarabichi, Peter Van Loo

## Abstract

Tumors frequently display high chromosomal instability (CIN) and contain multiple copies of genomic regions. Here, we describe GRITIC, a generic method for timing genomic gains leading to complex copy number states, using single-sample bulk whole-genome sequencing data. By applying GRITIC to 5,656 tumors, we found that non-parsimonious evolution is frequent in the formation of complex copy number states in genome-duplicated tumors. We measured CIN before and after genome duplication in human tumors and found that late genome doubling was followed by an increase in the rate of copy number gain. Copy number gains often accumulate as punctuated bursts, commonly after genome duplication. We infer that genome duplications typically affect the selection landscape of copy number losses, while only minimally impacting copy number gains. In summary, GRITIC is a novel copy number gain timing framework that permits the analysis of copy number evolution in chromosomally unstable tumors.

**Statement of significance:** Complex genomic gains are associated with whole-genome duplications, which are frequent across tumors, span a large fraction of their genomes, and are linked to poorer outcomes. GRITIC infers when these gains occur during tumor development, which will help to identify the genetic events that drive tumor evolution.

## Introduction

Copy number gains and losses are common somatic alterations in cancer (1,2). While somatic single nucleotide variants (SNVs) and indels linked to cancer drivers are found in ostensibly healthy tissues, copy number events rarely occur in normal cells (3–6). Identifying when copy number events occur is important for screening purposes and for gaining an understanding of the key molecular mechanisms underlying cancer development.

Of particular interest is the evolution of copy number events in tumors with the most aberrant genomes, as CIN is linked to poorer outcomes (7). Tumors that have undergone whole genome duplication (WGD) often show elevated aneuploidy (8–10) which arises from multiple factors, including chromosomal missegregation from centrosomal amplification (11) and a shortage of replication machinery proteins immediately following WGD (12). While increased aneuploidy supports CIN, the temporal relationship between WGDs and CIN is difficult to assess from single-timepoint biopsies in human tumors. The extent of genomic aberration, often used as an indirect proxy for CIN, does not convey the temporal dynamics that define CIN.

To observe the evolution of genomic gains in genome-doubled tumors, the timing of copy number gains and WGDs relative to the accumulation of SNVs can be inferred from whole-genome sequencing data (13–15). Clonal copy number gains, which are present in every tumor cell, can be placed on a timeline from 0 to 1, where 0 represents conception and 1 represents the end of the tumor’s clonal evolutionary period. Previous approaches that have used this principle to time copy number gains (16–18) were unable to fully time gains leading to complex copy number states (those with three or more copies of one parental allele). This is because of the higher ambiguity in the route history of these complex states relative to simpler states. Either the most parsimonious route history was assumed (17,18) or these states were not timed at all (16). Recently, a method was developed that can provide bounds on the timing of the first and last gains for complex copy number states (bioRxiv 2022.06.14.495959); however, it does not resolve the timing of the intermediate gains.

Here, we present GRITIC (Gain Route Identification and Timing In Cancer), a method that can time sequential gains leading to complex clonal copy number states, thereby elucidating the genome-wide evolution of gains in tumors with high CIN. As GRITIC is designed to time clonal copy number gains, it is well suited to unravel the evolution of the earliest genomic events in tumors, those that arise before the emergence of the tumor’s most recent common ancestor. We applied GRITIC to a cohort of 1,897 tumors from the Pan-Cancer Analysis of Whole Genomes (PCAWG) dataset (19) and 3,759 metastases from the Hartwig Medical Foundation dataset (20,21). Surprisingly, we observed that the commonly held principle of maximum parsimony (i.e., that copy number states are formed through the simplest possible route) is frequently violated for complex copy number gains in WGD tumors. We found that synchronous bursts of gains, independent of WGD, were common across cancer types. We infer the rate of gains pre- and post-WGD across our cohort and observe that late WGD causes an immediate increase in the rate of gains, a proxy for chromosomal instability. By considering the landscape of copy number events before and after a WGD, we found that WGD appears to have a low impact on the selection of copy number gains, but a greater impact on the selection of losses.

## Results

### GRITIC leverages SNVs to time complex copy number gains

Tumors frequently gain additional copies of their genomic regions during development. In the Hartwig and PCAWG datasets, copy number gains affected an average of 44.0% of the tumor genomes (51.6% and 31.6%, respectively). Complex gains were common, with 26.9% of the gained genome having three or more copies of one parental allele on average, predominantly major copy number three or four. This was 28.9% and 21.2% for Hartwig metastases and PCAWG primary tumors, respectively (Fig. 1A). The frequency of a given complex copy number state was inversely correlated with the largest number of copies of the parental allele for the state, known as the major copy number (Fig. 1A). Metastases also had a higher rate of WGD in our cohorts: 51.9% for Hartwig metastases compared to 28.7% for PCAWG primary tumors (Fig. 1B, S1A), although this appears to be a phenomenon specific to certain cancer types (Fig. S1B) (8,20). Although the difference in complex copy number fraction is largely explained by the higher proportion of WGD tumors, it was still higher in metastases when controlling for WGD frequency (Fig. 1C), likely reflecting increased CIN in metastatic cancers.

**Fig. 1.**
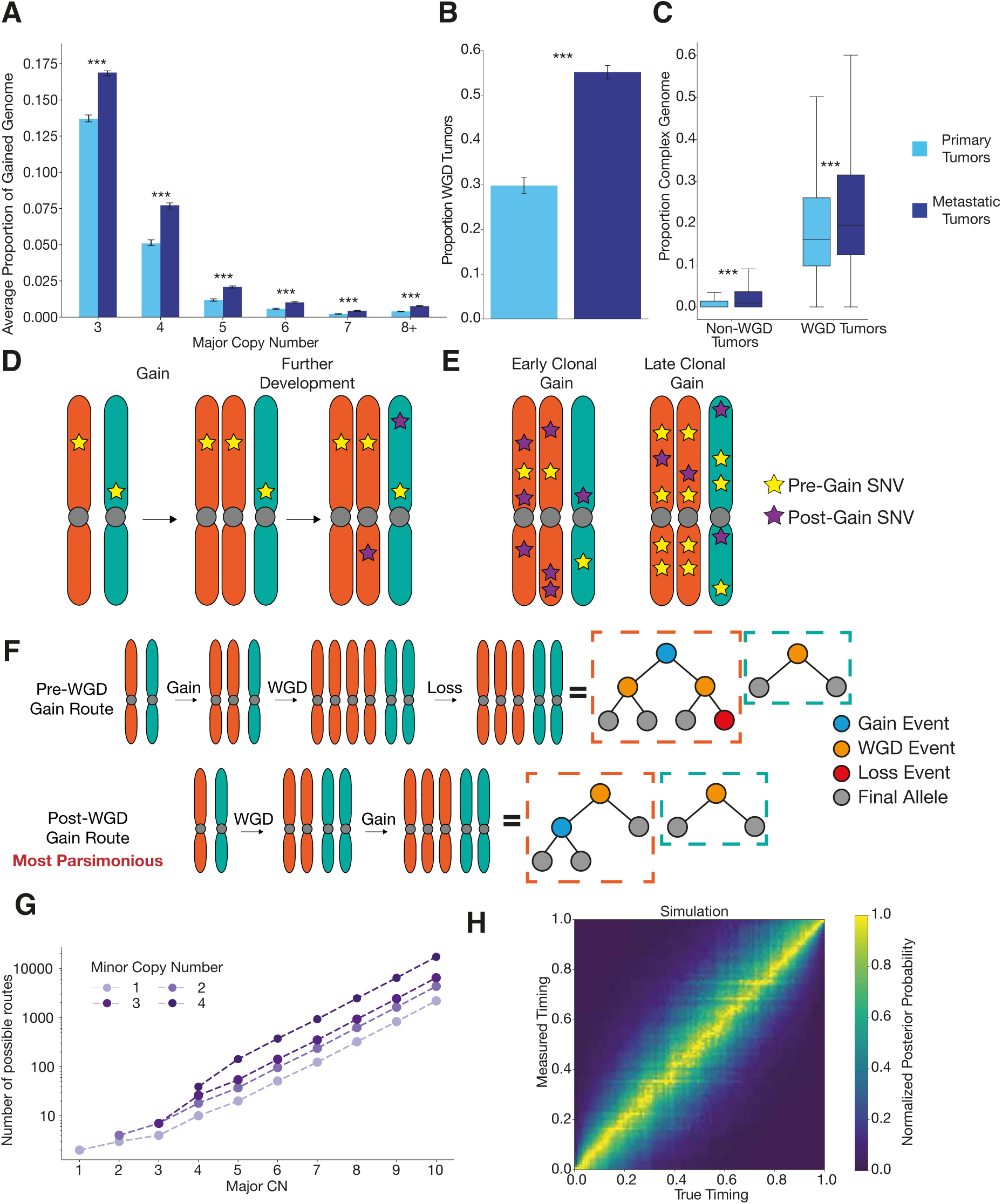
Principles of timing complex copy number gains. **A**, Average proportion of the genome with different major copy number states split by primary and metastatic cohorts. Statistical significance was calculated by permutation test and 95% confidence intervals by bootstrapping over samples. **B**, Proportion of tumor samples identified as WGD in primary and metastatic cohorts. Statistical significance is calculated by proportion test and 95% confidence intervals by normal approximation to a binomial proportion. **C**, Proportion of the genome with a major copy number of at least three in the primary and metastatic cohorts, split by WGD status. Statistical significance was calculated by permutation test and 95% confidence intervals by bootstrapping over samples **D**, Schematic showing SNVs on a gained allele are duplicated by the gain, the principle underlying copy number gain timing. **E**, Schematic showing the difference in SNVs on multiple copies between a gain that occurs early in the clonal evolutionary period and a gain that occurs later. **F**, Binary tree representation of two possible routes that result in a 3+2 copy number state in a WGD tumor. The post-WGD route is the most parsimonious as it involves the fewest events. **G**, The number of theoretically distinguishable unique routes that can result in different allele-specific copy number states given a single WGD. **H**, Distribution of measured posterior probability on gain timing against true gain timing across a representative simulated cohort of complex gains.

Clonal copy number gains can be quantitatively timed by considering SNVs in the gain region. When a gain occurs, all SNVs present on the gained allele are duplicated over to the new allele (Fig. 1D). With the reasonable assumption that each base pair in the genome is mutated at most once (22), any SNV on multiple copies in a gained region must have occurred before copy number gain. This principle can be used to infer the timing of the gain (14) (Fig. 1E, Supplementary Methods). However, for more complex gains, further consideration of the possible routes that lead to these complex states is required. To accomplish this, we developed GRITIC, a new method that can identify, distinguish, and time the gains in these routes.

GRITIC uses a binary tree representation conceptually similar to an earlier approach (23) to represent the gain history of a given segment. These representations can be used to calculate all possible routes (assuming, at most, a single WGD), resulting in a particular copy number state (Fig. 1F, Supplementary Methods). We found that the number of possible routes increases exponentially with the complexity of the copy-number state (Fig. 1G). Therefore, we limit GRITIC to the timing of copy number gains of segments with a total copy number of no more than 9. GRITIC uses a Bayesian Markov Chain Monte Carlo (MCMC) approach to infer the posterior probability of all possible route histories and the corresponding set of gain timings from SNV read counts for each gained segment. GRITIC is particularly suited to timing tumors with a WGD, as it uses the simultaneous occurrence of a WGD across all genomic regions as a constraint during inference to improve timing accuracy (Fig. S2, Methods).

We applied GRITIC to a realistic simulated cohort of WGD tumors (Methods) that contained equal proportions of all routes, leading to the 20 most common complex copy number states. We found that with simulated tumor purity and sequencing coverage representative of the tumors in PCAWG and Hartwig, GRITIC can accurately measure the timing of all gains, leading to complex states under different assumptions of the prevalence of parsimonious routes and penalties applied to non-parsimony during inference (Fig. 1H, S3-S7, Methods). Although more sensitive to simulation and inference conditions compared to measuring the gain timing itself, GRITIC can also accurately estimate the probabilities of different gain routes (Fig. S8-S11).

### Non-parsimony is common in WGD tumor gain evolution

We then applied GRITIC to time the gains that led to 131,296 clonally gained autosomal regions across 5,656 tumors in the PCAWG and Hartwig datasets. GRITIC reconstructs the timing of both the independent gains and the WGD (if present) in each sample and can time multiple sequential independent gains in the same genomic region (Fig. 2A).

**Fig. 2.**
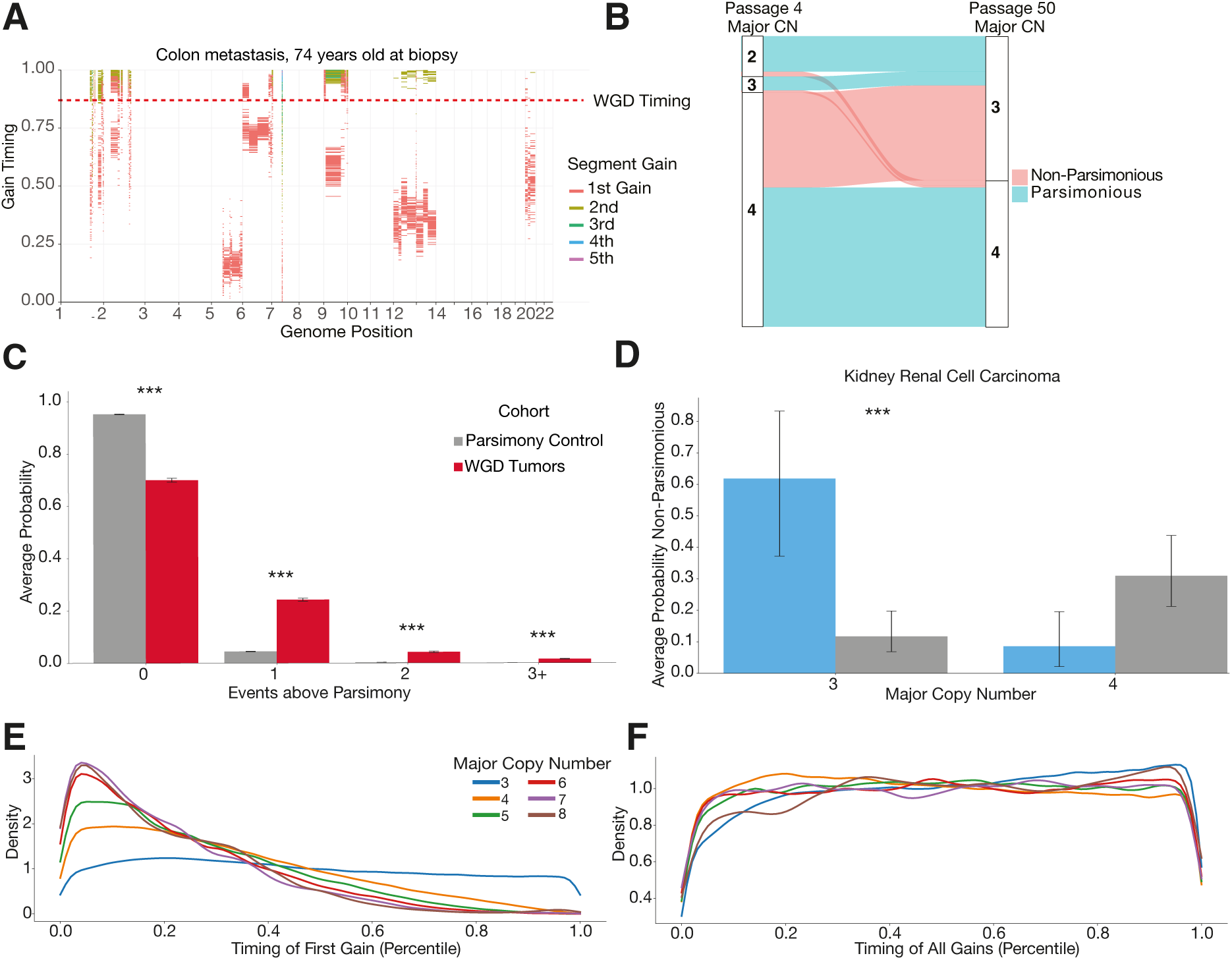
Non-parsimonious copy number evolution in cancer. **A**, Example posterior distribution of copy number gain timing in a whole-genome duplicated sample from Hartwig with GRITIC. 100 independent draws from the posterior sample for each gained segment are shown. **B**, The proportion of different copy number states at passage 4 that result in copy number states with a major copy number of 3 or 4 at passage 50 in four tetraploid clones in a colorectal cancer cell line. The flows are colored by whether the transition would be considered a parsimonious route by the copy number at passage 50. **C**, The average posterior probability on the number of additional events required to reach the final state over the most parsimonious route for complex gained states in the PCAWG and Hartwig cohort, compared to a simulated control where only parsimonious routes were included. A penalty on non-parsimony was applied during inference. Statistical significance is calculated with a permutation test and 95% confidence intervals by bootstrapping over samples. **D**, The average probability on non-parsimonious routes for gained segments in clear cell renal cell carcinoma, split by major copy number and gain location. A penalty on non-parsimony was applied during inference. Statistical significance was calculated with a permutation test and 95% confidence intervals were calculated by bootstrapping over samples. **E**, The distribution of the percentile timing of the first gains in complex segments relative to other gains in the same sample, split by major copy number. **F**, The distribution of the percentile timing of all gains in complex segments relative to other gains in the same sample, split by major copy number.

Parsimonious route histories have often been assumed for the development of copy number states in WGD tumors (17,18). This is because the total number of allelic copies gained through WGD versus individual independent gains will vary between different routes, as will the number of losses required to make the route self-consistent (Supplementary Methods). Therefore, we sought to evaluate the assumption of parsimonious evolution in WGD tumors.

We first tested this assumption by reanalyzing copy number data from isogenic tetraploid colorectal cancer HCT-116 cell lines obtained from two different passages derived from a diploid progenitor (24,25). By considering the change in copy number states between the two passages, we found that most events with a major copy number of three in the later passage arose through complex routes that would violate the assumption of parsimony if applied to the later passage in isolation (Fig. 2B). This result suggests that the assumption of parsimonious copy number evolution is often invalid.

We used GRITIC to test this parsimonious evolution assumption. Owing to the inherent uncertainties (Supplementary Methods) in estimating copy number gain routes from SNVs, GRITIC assigns an average posterior probability of 51.1% to non-parsimonious routes from a simulated set of tumors with completely parsimonious evolutionary histories (Fig. S12A). Although the model evidence used in GRITIC provides a natural penalty against additional independent gain timing parameters, it does not penalize the number of losses implied by a given route. Thus, to ensure a conservative estimate of non-parsimony, we applied a penalty term to the number of events required for each route in WGD tumors (Methods). This penalty was fitted such that the average posterior probability of non-parsimonious routes was less than 5% on a representative cohort of simulated WGD tumors with only parsimonious routes (Fig. S13-14).

With this penalty term, we evaluated non-parsimonious evolution across genome-duplicated PCAWG and Hartwig tumors. Surprisingly, we found that non-parsimony was common across cancer copy number evolution: 30.0% of the total posterior probability on gained segments in WGD tumors was on non-parsimonious route histories, with 5.8% on routes with two or more events in addition to the simplest route (Fig. 2C, S12B, p<0.001, permutation test). Owing to our conservative penalty term, these are likely underestimates of non-parsimony in copy number evolution.

Non-parsimony occurs in agreement with known phenomena. Gains on chromosome 5q are known to be and initiating event in clear cell renal cell carcinoma, combined with the loss of 3p (26). In line with this, GRITIC inferred that gains on chromosome 5 with a major copy number of 3 are significantly more non-parsimonious (*i.e.* earlier, Fig. S12C, Methods) than the background for clear cell renal cell carcinoma (Fig. 2D, p<0.001, permutation test). Conversely, gains on chromosome 5 with a major copy number of four were more likely parsimonious (*i.e.* earlier, Fig. S12C, Fig. 2D, p=0.059, permutation test). This effect was also observed when a non-parsimony penalty term was not applied (Fig. S12D-E). More generally, we found that the frequency of pre-WGD gains on chromosomes was highly correlated (Pearson’s, p<0.001) across cancer types between major copy number 3 and 4 states, even though such gains were non-parsimonious for major copy number 3 (Fig. S15).

Applying a penalty to the number of events provides a useful lower bound on the non-parsimonious evolution of complex gains. However, we found that, as expected, this causes the inferred probability of non-parsimonious evolution to be very inaccurate for cohorts simulated to contain non-parsimonious routes (Fig. S9). Therefore, for all subsequent analyses, we show the results of applying GRITIC without this non-parsimony penalty term, and display the results with a penalty term in the Supplementary Information. In general, both results are highly consistent.

We find that the initial gains that occur independently of the WGD tend to occur earlier as the major copy number (up to 8) increases. They are typically among the first gains in the tumor genome (Methods, Fig. 2E, S16). Moreover, when considering all copy number gains that occur independently of WGD, segments with high copy numbers have gains dispersed across tumor development (Fig. 2F, S16). This suggests that moderate-level amplifications in tumor development generally begin early but can accumulate gains throughout the clonal evolutionary period.

We speculated that elevated CIN post-WGD may lead to an increase in copy-neutral events, where the loss of one parental allele is combined with a gain of the other. Post-WGD copy-neutral events would cause a 2+2 state to become a 3+1 state, which may then go on to undergo further gains. Indeed, we found that regions in WGD tumors with a minor allelic copy number of one were significantly more likely to have a first gain post-WGD than other copy number states (Fig. S17A-B), much more than in the cohorts simulated without this effect (Fig. S17C-D).

### The effect of genome doubling on the rate of chromosomal instability

Next, we evaluated the effect of genome doubling on the rate of CIN. We analyzed the copy number profiles of 260 individual tumor cells in an undifferentiated sarcoma that had undergone consecutive subclonal WGDs, as shown experimentally (Methods). We found progressively higher rates of inter-copy number diversity in cells with each round of WGD (Fig. 3A, S18, p<0.001, Mann-Whitney U Test, Methods), suggesting that WGD increases CIN even within the same tumor.

**Fig. 3.**
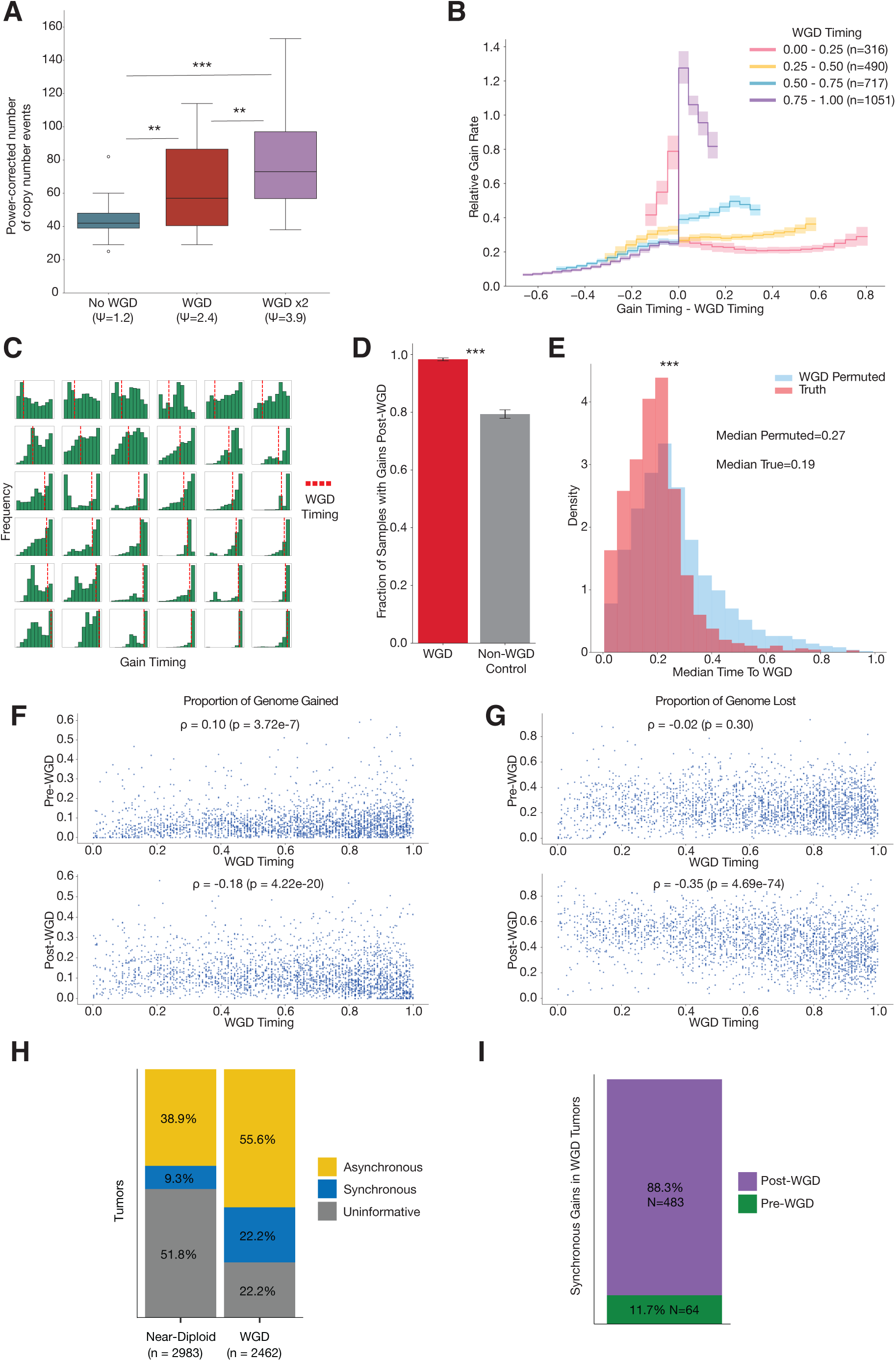
The history of chromosomal instability over tumor evolution. **A**, The number of copy number events required to reach the final copy number state from the most recent common ancestor from a collection of tumor cells with different ploidy states from a single undifferentiated sarcoma. The number of events is normalized for ploidy and statistical significance is calculated with a Mann-Whitney U test. Ψ denotes population ploidy as determined by FACS. **B**, The normalized rate of gains relative to the WGD timing across cancer types. Statistical significance is calculated with a permutation test and 95% confidence intervals are calculated by bootstrapping over samples. **C**, Histograms of gain timing in a random selection of tumors with a WGD. Samples are ordered by independent gain timing from left to right and WGD timing from top to bottom. **D**, Proportion of samples with gains post-WGD for WGD tumors and a cohort of control non-WGD tumors with a pseudo-WGD timing randomly sampled from WGD tumors with the same cancer type. Statistical significance is calculated with a permutation test and 95% confidence intervals are calculated by bootstrapping over samples. **E**, Distribution of the mean mutation time between the timing of all independent gains and the median WGD timing for each genome duplicated sample. Two cohorts are displayed, one with the correct WGD timing and another where the WGD timing is permuted between samples of the same cancer type. Statistical significance calculated by Mann-Whitney U test. **F**, Proportion of genome gained before and after WGD against WGD timing for genome doubled tumors. **G**, Proportion of genome lost before and after WGD against WGD timing for genome doubled tumors. **H**, Proportion of tumors with clonal gains identified as occurring synchronously or asynchronously, or uninformative where the number of gains was too low to classify, split by WGD status. **I**, Proportion of synchronous gains occurring in WGD samples, classified by whether they occurred pre- or post-WGD.

We used GRITIC to calculate the rate of independent gains, a proxy for the rate of CIN, relative to the occurrence of WGD across our cohort. We found that the gain rate increased after WGD but only in tumors with late genome doubling (Fig. 3B). While this increase in the rate of gains remains upon the application of the non-parsimony penalty, owing to uncertainties in placing gains directly before or after a WGD, it leads to a greater immediate increase in the post-WGD gain rate (Fig. S19).

We observed that, regardless of the penalty on non-parsimony, the tumors with the earliest genome doubling have a different trend in gain rate relative to WGD. They exhibited a high rate before early WGD and a subsequent steady post-WGD gain frequency, which was lower than that before WGD (Fig. 3B). This behavior may be driven primarily by breast tumors, which were enriched in this group (Fig. S19), which is consistent with reports of highly aneuploid karyotypes of early precursor lesions of breast cancers (27). While only a few individual tumors followed such clear patterns (Fig. 3C), the effect of WGD on CIN in aggregate was clear.

There was also a general increase in CIN over the clonal evolutionary period in non-WGD tumors (Fig. S20), there was a significant further excess of gains in WGD tumors: the proportion of WGD tumors that gained post-WGD (98.3%) was significantly higher than that in a control cohort of non-WGD tumors with pseudo-WGD timing (79.4%) (p<0.001, permutation test, Fig. 3D, S21A-B). Moreover, the mean mutational time between independent gains and WGD was significantly lower than expected from permuting WGD timing (p<0.001, Mann-Whitney U Test, Fig. 3E, S21C-D, S22, Methods), suggesting that this excess is subsequent to WGD.

Although the later a WGD occurs, the less time there is for gains to accumulate post-WGD, the amount of genome gained post-WGD was only weakly negatively correlated with WGD timing (Spearman’s ρ=-0.18, p<0.001, Fig. 3F, S23). Similarly, the genomic material gained pre-WGD was weakly positively correlated with WGD timing (ρ=0.10, p<0.001, Fig. 3E, S24). In contrast, there was a much stronger negative correlation between WGD timing and the proportion of genome lost post-WGD (ρ=-0.35, p<0.001, Fig. 3G, S25A). Generally, these trends are conserved when cancer types are considered separately and after applying the non-parsimony penalty (Fig S23, S24, S25A).

In contrast, there was no correlation between WGD timing and the proportion of the genome that was lost pre-WGD. This suggests that such losses pre-WGD do not accumulate steadily with respect to mutation time or are linked to WGD (Fig. 3G, S25B). This could support a model whereby a major advantage of WGDs is the mitigation of the effect of deleterious mutations in regions with loss of heterozygosity (LOH), as reported by Lopez *et al.* (25). If so, the greater the proportion of the genome lost in a non-WGD sample, the more likely it is that a WGD has a positive selective impact. This selective impact would remove the relationship between the amount of genome lost pre-WGD and the timing of WGD. This is supported by observations of a higher proportion of LOH in the genomes of WGD tumors than in non-WGD tumors (25). The high frequency of post-WGD losses is consistent with the widespread hypothesis that, in many cancers, WGD serves as an evolutionary intermediate to fitter sub-tetraploid karyotypes (28).

### Punctuated gain evolution in WGD tumors

Next, we sought to better understand the distribution of gain timing in WGD samples. We previously found that copy number gains in non-WGD tumors occurred in bursts that were closer than expected by chance (17). Indeed, using GRITIC, we found that gains in 19.3% of informative non-WGD samples (those with gains affecting at least three separate chromosomes) occurred over significantly shorter timespans than expected under a permutation model (Fig. 3H, S26A-D). Methods).

Using GRITIC, we can now study the timing of gains that arise independently of the WGD. We found that gains in 28.5% of informative WGD tumors occurred significantly closer in time than expected from permutations (Fig. 3H, S26A-D). Most of these (88.3%) occurred post-WGD (Fig. 3I, S26E), suggesting that WGD may increase the likelihood of or tolerance to punctuated bursts of gains (24). Together, these results suggest that copy number gains occur frequently in punctuated bursts, even in the most chromosomally unstable samples.

### Measuring the impact of a WGD on the copy number landscape

The landscape of gains and losses along the genome is similar for WGD and non-WGD tumors across cancer types (9). Therefore, we sought to determine whether the selection landscape of copy number events is different before and after a WGD. First, we compared the relative frequency of arm-level copy number gains across different cancer types pre- and post-WGD and found that they were highly correlated (Spearman’s ρ=0.81, p<0.001, Fig. 4A, S27-S28).

**Fig. 4.**
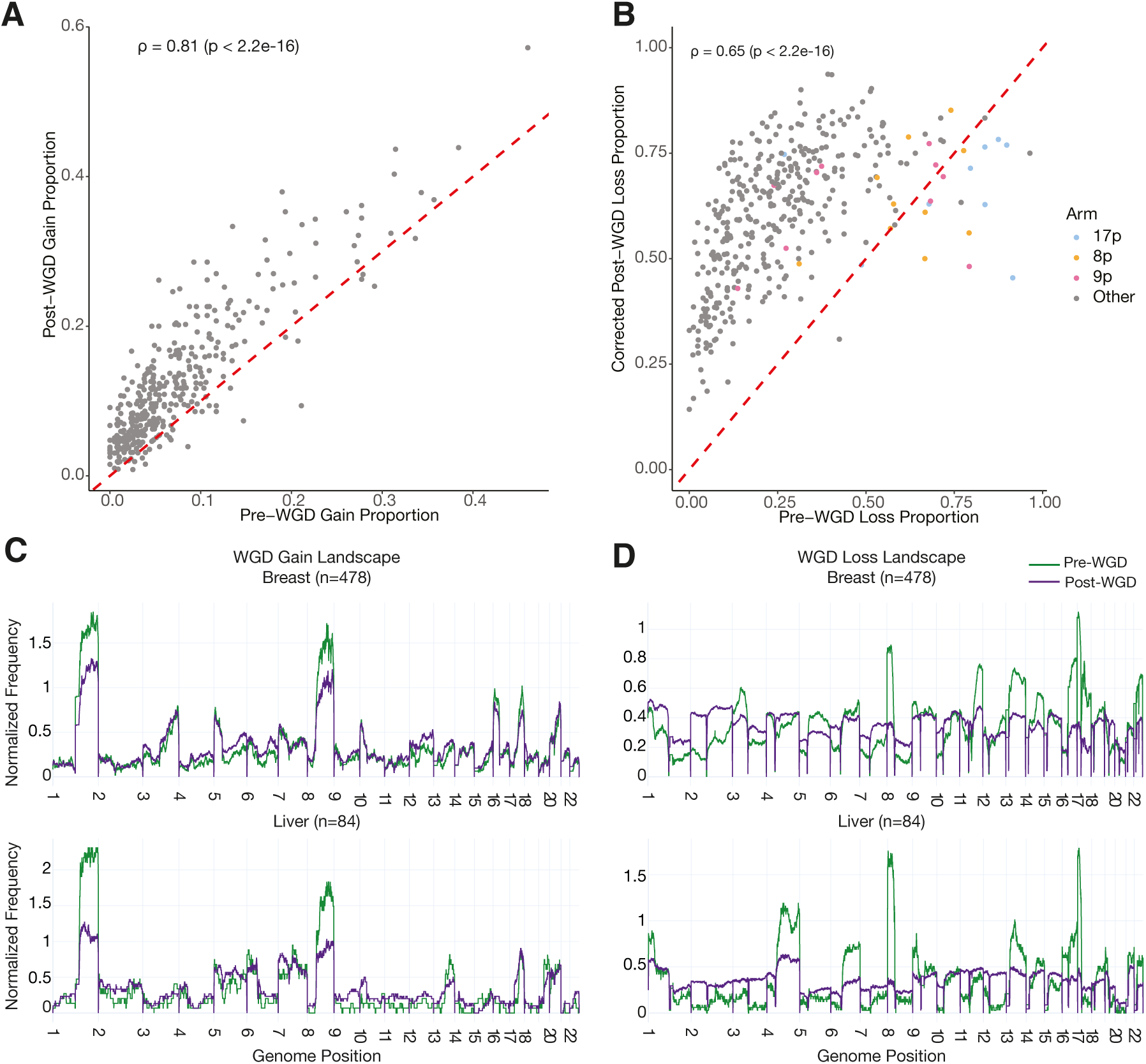
The landscape of pre- and post-WGD copy number events. **A**, Frequency of arm-level gain events occurring before and after a WGD in different cancer types. **B**, Frequency of arm-level gain events occurring before and after a WGD in different cancer types. The post-WGD arm loss frequency is corrected for mutual exclusivity in measuring pre-WGD and post-WGD arm losses in the same region. **C**, Frequency of pre- and post-WGD gains for breast and liver tumors. **D**, Frequency of pre- and post-WGD losses for breast and liver tumors. The post-WGD loss frequency is corrected for mutual exclusivity. Both frequencies are normalized so that the pre- and post-WGD frequencies integrate to the same arbitrary constant.

We then investigated how arm loss frequencies were affected by WGD. After correcting for mutual exclusivity in measuring the rates of loss before and after WGD (Methods), we also observed clear positive correlations (Spearman’s ρ=0.65, p<0.001, Fig. 4B, S29). This suggests that losses without loss of heterozygosity (LOH) observed post-WGD are mostly derived from a continuation of the same processes that lead to pre-WGD LOH losses. However, there are clear outliers where the pre-WGD loss frequency is much higher than expected given the corresponding post-WGD loss frequency.

Of the 21 arms across different cancer types that had a higher loss proportion pre-WGD than post-WGD, 16 were 8p (n=4), 9p (n=3), or 17p (n=9) (Fig. 4B). Interestingly, chromosome arms 9p and 17p contain well-known and frequently hit tumor suppressor genes, such as *CDKN2A* and *TP53*, respectively, whereas 8p harbors the helicase-encoding gene *WRN*, which has recently been identified as a likely tumor suppressor (29). Thus, while gains and losses are broadly unaffected by a WGD, we find specific arm-level losses that occur disproportionately pre-WGD. We note that most arms had higher event rates post-WGD than pre-WGD, reflective of increased CIN post-WGD. In agreement with previous findings (10,30), we found that the frequencies of chromosome arm events maintained similar levels of correlation/anticorrelation with the overall tumor suppressor gene and oncogene density (30) across pre-WGD, post-WGD, and non-WGD copy number events (Fig. S30-31). Despite certain arms having significantly higher pre-than post-WGD loss, the frequency of both event types showed a similar correlation with driver gene density across chromosome arms. This suggests that the disproportionate pre-WGD loss frequency of certain chromosome arms may be due to specific genes, perhaps in the context of a second inactivating hit, rather than the overall tumor suppressor gene density.

We then normalized the relative rates of pre- and post-WGD events for both gain and loss across the genome. We found that the rates of pre- and post-WGD gains were similar when aggregated across cancer types (Fig. S32). However, there are notable differences in individual tumor types. For example, gains on 1q and chromosome 8 were disproportionately likely to occur pre-WGD relative to other events (Fig. 4C, S32-S36). Gains on chromosome 3 are often pre-WGD for small-cell lung cancers and upper respiratory tract carcinomas (Fig. S35-S36). Chromosome 7 is commonly gained pre-WGD in glioblastomas, in agreement with previous observations that these events occur very early (17) (Fig. S33).

Interestingly, the differences were much larger pre- *vs.* post-WGD for losses, both at the aggregate level and in individual cancer types (Fig. 4D, S37-S39). As discussed earlier, chromosome arms 8p, 9p, and 17p are frequently lost pre-WGD in several cancer types (Fig. S37-S39). We observed many cancer-type specific patterns, such as high levels of pre-WGD loss of chromosome 18 in colorectal carcinomas (Fig. S37). Together, our results suggest a model in which the selection landscape for copy number changes remains broadly similar post-WGD, yet the selective pressure on certain chromosome arms is reduced if not removed, particularly for losses.

## Discussion

GRITIC is a Bayesian framework for genome-wide timing of both simple and complex gains in cancer evolution. It leverages the relationship between alternate read counts, copy number, and purity to determine the sequence and timing of gains. GRITIC theoretically allows the timing of gains from any copy number state, although in practice for computational efficiency and timing accuracy, we focus on autosomal segments with a maximum total copy number of nine and at least 20 SNVs, respectively.

By applying GRITIC to the PCAWG and Hartwig datasets, we measured the genome-wide timing of gains relative to WGD. Therefore, we describe the effect of WGD on chromosomal instability. We found that late WGDs tended to induce a spike in gain activity, which remained elevated compared with the pre-WGD gain rate. Conversely, early WGDs were preceded by an elevated gain rate, which then decreased post-WGD. This increase in genomic instability likely enables WGD tumors to have greater adaptivity in response to therapy, contributing to their poor prognosis (28). Because losses cannot be quantitatively timed, the gain rate over mutation time was not corrected for total genomic content. However, the contrasting patterns observed for tumors with different WGD timings indicate that an increase in CIN post-WGD does not solely result from more chromosomes missegregating at the same rate as before the WGD.

We found that the landscapes of gains occurring before and after genome duplication were similar. This parallels the similarity in gain landscape between tumors with *vs.* without WGD. We hypothesize that this is because WGD only has a very moderate impact on the gain fitness landscape of tumors. Although the pre- and post-WGD landscapes are also broadly similar for losses, there are selective pressures to lose certain chromosome arms encoding well-known tumor suppressor genes before WGD, leading to LOH. Post-WGD losses accumulated much more linearly with respect to the overall mutation burden than other copy number events. This could suggest a lower selective impact relative to other copy number events and a role in evolution that is more akin to the accumulation of passenger mutations.

We discovered that many segments across different copy number states in WGD tumors arose through routes that violate the principle of maximum parsimony even after this was substantially penalized. It is worth noting that we inferred the most parsimonious routes from single biopsies only. Event histories that appear non-parsimonious, as measured in one biopsy, may be parsimonious when the full heterogeneity of copy number in the tumor is considered. While GRITIC can only identify non-parsimony in gain evolution in individual segments, it calls into question the validity of the assumption of maximum parsimony in cancer evolution, particularly in the context of inference from a single biopsy.

GRITIC considers the gains in different segments separately. Future work incorporating structural variant information into timing analyses would enable more integrated evolutionary analyses of copy number changes across the genome (23). Similarly, as our inference is limited by ambiguities in resolving routes from SNV read counts, phasing SNVs either to haplotypes or ideally to specific alleles will substantially reduce this ambiguity and allow greater resolution of evolution. Currently, we are unable to time events in tumors with multiple WGDs, which we estimate to be 4.9% of the patients in the PCAWG and Hartwig cohorts (Methods). While this is theoretically possible in our framework, more work is required to build an inference pipeline that can link the complex gained states across segments.

In summary, GRITIC is a novel computational framework for inferring copy number gain evolution from a single bulk whole-genome sequencing experiment. It can also be applied across tumors from patient cohorts to reconstruct more complete evolutionary timelines (17) and to better understand CIN, particularly in relation to WGD.

## Methods

### Data collection

Whole genome sequencing, alignment, mutation calling, and copy number data were obtained from Pan-Cancer Analysis of Whole Genomes (19) and Hartwig Medical Foundation datasets (20). Clonal consensus copy number calls were used for PCAWG samples, and PURPLE copy numbers were rounded to the nearest integer for each allele in each Hartwig sample. The cancer type classifications for each sample were obtained from a combined set of PCAWG and Hartwig classifications (21). SNV clustering information was obtained from consensus clustering for PCAWG (31) and by using the default settings of DPClust (15) for Hartwig. DPClust was run for 2,000 iterations with 1,000 burn-in steps.

### Kataegis Filtering

SNVs that were identified as being likely to have arisen through kataegis were identified using an existing analysis pipeline (19). These were filtered from the data because kateagis is a localized hypermutation process that leads to sets of SNVs that violate the assumption of constant relative mutation rates across the genome.

### WGD Calling

To identify WGDs, we calculated the cumulative number of base pairs spanned by each clonal major copy number state and identified the major copy number state spanning the highest number of total base pairs, *i.e.* the mode of the major copy allele. If the mode was one, the sample was identified as non-WGD. If it was two, we calculated the individual timing of all two segments of the major copy number. A core principle of GRITIC is that WGD causes simultaneous gain across the genome. Therefore, if at least 60% of the base pairs spanned by the major copy number two segments had posterior gain timing distributions with overlapping 89^th^ percentiles, a reasonable choice for posterior credible intervals (32), then the sample was identified as WGD.

This provided WGD calls consistent with those provided by the datasets using copy number profiles alone (Fig. S40A-B). Notably, the samples with a major copy number mode of two but with less than 60% timing overlap were enriched in lung and skin tumors, which are tumor types known to have late copy number gains (Fig. S40C). Indeed, the timing of the maximum overlap was later in mutation time compared to tumors with greater than 60% overlap in the timing of major copy number two gains (p<0.001, Mann-Whitney U Test, Fig. S40D).

### Sample filtering

Across PCAWG and Hartwig, whole-genome sequencing information was obtained for 7,502 tumors. We then applied several quality filters to this cohort.

To ensure sufficient coverage for SNV multiplicity inference, we required samples with an NRPCC of at least 5. NRPCC is the expected number of reads in a tumor for a clonal SNV with a multiplicity of one (33). It is calculated as:

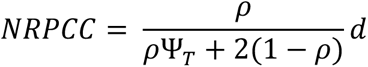

Here, ⍴ is the tumor purity, Ψ_T_ is the tumor ploidy and *d* is the depth of sequencing. In total, 62 tumors had an NRPCC of less than 5. For patients with multiple tumor samples, we selected one representative sample that had the highest NRPCC. This led to the removal of 431 tumor samples.

We filtered samples in which the major copy number mode was greater than two. This is because these tumors likely had more than one WGD, an evolutionary history that is not supported by GRITIC. This encompassed 381 tumors in both cohorts. The 209 tumors with a major copy number mode of two but with less than 60% overlap of the timing of major copy number two segments by total base pairs were also excluded, as they were likely genome duplicated, but violated the principle of simultaneous gains required by GRITIC.

Only clonal copy number gains in segments with at least 20 SNVs and a total copy number of no more than 9 were timed. A gained region was defined as a copy number segment with a major copy number of >2 for WGD tumors and >1 for non-WGD tumors. Of all the gained segments in the PCAWG and Hartwig cohorts with a major copy number of three or greater, 97.0% had a total copy number of nine or less. Of the tumors with a major copy number mode of one or two, 917 tumors from the combined cohort were not included as they contained no independently gained autosomal regions with sufficient SNVs.

After applying all filters together, 1,846 tumors were removed. This left 5,656 tumors for the analysis.

### GRITIC

GRITIC enumerates the binary tree structures that represent all routes to a given gained copy number state for a particular segment, accounting for the presence of up to one WGD. These representations are used to sample the SNV multiplicity proportions that correspond to the range of possible gain and WGD timing for each route. The relative probability of each route was calculated using the likelihood of the SNV read counts for the segment given each sampled multiplicity proportion. A full description of the GRITIC method and its principles is provided in the Supplementary Methods file.

### Sample simulation

We tested GRITIC using simulated samples with known ground-truth gain timing and route histories. We aimed to make the simulations as representative of true samples as possible. We randomly selected a WGD tumor from Hartwig and the PCAWG to form the basis of each simulated sample. The original copy number profile and purity of the selected base tumors were maintained in the simulated sample. As we were primarily interested in the copy number profile of the base tumor, the WGD status was taken from classifications provided by the original datasets and not GRITIC.

The true WGD timing for the sample was randomly selected from the distribution of the WGD timings measured by applying GRITIC to real PCAWG and Hartwig samples. The multiplicity proportions were set to match the simulated ground-truth timing for all segments with sufficient SNVs. Segments with major two copy number were simulated to have multiplicity proportions corresponding to a gain occurring at the time of the WGD. The gain timing and corresponding clonal multiplicity proportions for higher copy number states were simulated by selecting the 250th sample from our MCMC sampler with a randomly selected route. The proportion of subclonal mutations was directly obtained from SNV clustering information for each sample. As in the main GRITIC method, all subclonal SNVs were assumed to have a multiplicity of one.

The number of SNVs in each multiplicity state was obtained from a multinomial draw using clonal and subclonal multiplicity proportions and the number of SNVs in the gained segment from the base tumor. Alternative read counts for each SNV were then drawn from a binomial distribution using the variant allele frequency equation (33) for the given multiplicity and total coverage drawn from a Poisson distribution with a mean equal to the average read coverage of the segment in the base sample.

Two cohorts were simulated, one where all routes for a given copy number state were selected equally, and one where only the most parsimonious routes were selected.

### Measuring performance of gain timing inference

We used a probabilistic approach to compare the gain timing inferred from GRITIC with the simulated ground truth. For each simulated segment and route, we sorted all inferred gain timings and all true timings and compared these timings pairwise. Although the number of independent gains differs between copy number states, the total number of gains, including those inferred to arise through WGD, is the same across all routes.

Only comparisons corresponding to independent gains in the sorted true cohort were collected. The pairwise comparisons for each route were stored in a histogram and each comparison from every route was weighted according to the inferred route probability from GRITIC.

### Assessing route probability calibration

We assessed the calibration of the gain route probabilities inferred from GRITIC using a binning approach. We placed the probability of each gain route for all simulated segments in 20 evenly spaced bins from zero to one. The true probability for each bin was then calculated from the mean number of routes in each bin that were the true simulated route for the corresponding segment. The model was judged to be well calibrated if this true probability fell within the corresponding bin.

### Assessing non-parsimony from cell line data

We used existing cell line data with provided copy number calls (25). We examined the copy number of tetraploid clones at passages four and 50 and measured any change in copy number at each base pair between passages. We plotted the transitions weighted by segment width between major copy number two, three and four segments at passage four, and major copy number three and four at passage 50. Each transition was labeled as parsimonious if the major copy number at passage four would have been expected under the most parsimonious route history expected from the major copy number at passage 50.

### Measuring non-parsimony probabilities

We assessed non-parsimony across the simulated and PCAWG and Hartwig cohorts by examining the total probability assigned to routes for each gained segment that had more events than the route with the minimum number of events for the segment. The average non-parsimony was aggregated across the cohort.

### Penalty on non-parsimony

As discussed in the main text, the model evidence used in the computation of the probability for each route provides a natural penalty against additional independent gains because the evidence is integrated over the extra parameters required to model the timing of these additional gains. However, this provides no penalty for the additional loss events required to make each route consistent.

Therefore, we applied a second penalty term P to the number of events n implied by each route.

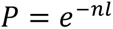

We tuned the penalty parameter *l* on a representative simulated cohort of simulated tumors to have only parsimonious routes. We set *l* such that the total probability of non-parsimony across the cohort was approximately 5%. As this will depend on the exact setup of the simulation, we did not tune l precisely; instead, we found that *l*=2.7 was a reasonable penalty, giving ∼5% (4.8%) total non-parsimonious probabilities.

### Measuring non-parsimony in clear cell renal cell carcinomas

We selected the 48 genome-duplicated clear cell renal cell carcinomas from PCAWG and Hartwig that contained gains with a major copy number of three or four and a minor copy number of no more than two. We filtered segments with at least a 50% probability on the most parsimonious or non-parsimonious route(s) with the fewest number of events. These copy number states and routes were selected because they have a clear relationship between parsimony and gain timing. For segments with a major copy number of four, the most parsimonious route had the first gain pre-WGD, and the two non-parsimonious routes with the joint lowest number of events had the first gain post-WGD. The reverse is true for major copy number three segments, where the sole non-parsimonious route with the fewest number of events has a single pre-WGD gain.

We aggregated the average non-parsimonious probability for each chromosome weighted by segment base pair length, normalizing the route probabilities within the selected routes. Statistical significance was calculated using a permutation test and 95% confidence intervals by bootstrapping samples.

This analysis was restricted to the two most parsimonious routes for copy number states with major copy numbers of three and four and a minor copy number of two. There are more complex non-parsimonious routes that exist for these copy number states, but the situation is more complex because the combination of gains and losses means that there is no clear relationship between non-parsimony and gain timing. We only included segments that had at least 50% of their posterior probability on the two most parsimonious routes. With the non-parsimony penalty, this accounted for 96.9% by base pair length of the gains in the clear cell renal cell carcinomas. Without the penalty, the fraction was reduced to 74.9%, but similar patterns in terms of non-parsimony were observed.

### Measuring Timing of Major Copy Number Gains

We sought to compare the timing of gains in different major copy number states across tumors. We calculated the relative percentile rank of each posterior sample of gain timing across the combined posterior distribution over all segments for a given tumor. The percentile ranks were directly compared between the samples. We computed and compared two percentile rank distributions: one with only the initial gain that leads to each complex state and a second with all gains.

### Measuring CIN in single cells

We utilized 10x single-cell whole genome sequencing data from a single undifferentiated sarcoma (34). The tumor contained three distinct tumor cell sub-populations with average ploidies of 1.2, 2.4, and 3.9, respectively, which were determined by fluorescence-activated cell sorting. Based on these ploidies and their copy number profiles, the diploid and tetraploid populations are likely to represent cells resulting from two rounds of WGD following an initial haploidization event. We ran ASCAT.sc (https://github.com/VanLoo-lab/ASCAT.sc) using phased germline SNPs generated by Battenberg (15) on a matched normal to obtain allele-specific copy number profiles for all tumor cells. Noisy cells were removed based on the standard filtering metrics in ASCAT.sc.

Cells with a higher ploidy can have genomes with a greater number of potential copy number states. To obtain an estimate of CIN corrected for this bias, all cell populations were fitted with copy number profiles with ploidy between 3.7 and 5.5.

We then used MEDICC2 (35) to build copy number phylogenies for each subpopulation separately, with a minimum segment length of 2.5Mb. The copy number heterogeneity of each population was inferred by measuring the number of copy number events required to transform the common ancestral profile (as inferred by MEDICC2) into each of the cells in the subpopulation.

To account for a small number of cells with noisy copy number profiles which were not removed by ASCAT.sc, the ancestral profile was defined as the profile inferred by MEDICC2 to be the ancestor of 90% (or the closest possible fraction) of cells in the population. Only cells that were measured to be descendants of this ancestral profile were considered.

### Calculating the rate of gains relative to WGD

The rate of gain relative to WGD was measured by summing the posterior density multiplied by the segment base pair length across 20 bins of *gain timing – WGD timing* across the cohort. The bins were evenly distributed through the central 90^th^ percentile of the *gain timing – WGD timing* distribution with the smallest possible offset added, such that one bin started at zero.

The binned gain-timing distribution needs to be normalized to account for the distribution of WGD timing in each cohort, as the maximum possible time for a gain to occur before and after the WGD is dependent on the WGD timing. Therefore, we normalized the binned posterior density by dividing it by a second binned posterior distribution that used the same samples and the corresponding WGD timing distribution. However, in this normalizing distribution, the gain timing was uniformly distributed across mutation time, and the gains in each sample were weighted by segment base pair length such that each sample contributed equally. This normalizing distribution therefore represented the change in *gain timing – WGD timing* that would occur purely from differences in WGD timing alone. The scale of this distribution was standardized such that the mean of the bins was one.

### Measuring the fraction of samples with gains post-WGD

A tumor was identified as having a gain post-WGD if at least 50% of the posterior gain timing samples for any segment had at least one gain post-WGD. A control cohort of non-WGD tumors with a pseudo-WGD timing distribution was constructed by randomly choosing a WGD timing distribution for each non-WGD tumor from WGD tumors with the same cancer type. Only cancer types with at least ten WGD and ten non-WGD samples were considered.

### Measuring difference in timing between gains and WGD

We measured the median difference in timing between the posterior gain timing distribution over all the gained segments and the WGD timing sampled from the WGD timing distribution for the tumor. We repeated this process for 25 samples of the WGD distribution for each tumor and calculated the average median difference in timing. This resulted in a distribution of the average difference between WGD and gain timing for all WGD tumors in our cohort.

This distribution was compared to a control distribution calculated identically, except that all WGD distributions were randomly permuted between WGD tumors of the same cancer type.

### Measuring gain synchronicity

We aimed to identify tumors that had gains in different chromosomes that occurred closer together in mutation time than would be expected under independent draws from the distribution of gain timing. Such samples were classified as having a synchronous or punctuated set of gains. Following an approach similar to that of Gerstung *et al.* (17), we performed a permutation-based procedure at the tumor level to obtain a threshold to define synchronous gains for each sample. We restricted this analysis to tumors that had at least 15 samples of the same primary tumor type. Additionally, tumors with fewer than five gained chromosomes were excluded and classified as uninformative.

To identify synchronous gains in each sample, we defined a synchronicity score for each sample, which we term the median weighted gain timing distance. First, we created pseudo-samples with exact gain timing for each segment by drawing from the posterior timing distribution of each gained segment. We calculate the point in the mutation time that has the minimum distance to the timing of all segments weighted by the segment width. For segments with multiple gains, only the distance between the gain and closest timing was measured. A total of 100 pseudo-samples were drawn for each sample, and the median minimum weighted gain timing distance across the samples was computed.

We then compared the median distance to that obtained under a null distribution, obtained with 1,000 independent permutations performed with different random seeds for all samples using the curveball method (36), which randomly swaps chromosomes across patients within the same cancer type, while maintaining the number of chromosomes gained per patient and the number of gains per chromosome across the cohort. Each tumor was independently defined as synchronous if its median weighted gain timing distance was lower than 95% of the permuted samples for that tumor. This can be interpreted as gains within a sample occurring closer in mutational time than expected by chance, for a given cancer type.

### Inferring the pre- and post-WGD landscape of copy-number gains and losses

We assessed a chromosome arm as having a pre- or post-WGD gain in a WGD tumor if at least 50% of the total base pairs in the arm belonged to segments with at least 50% of posterior gain timing samples and at least one gain before or post-WGD, respectively. An arm was classified as having a pre-WGD loss if at least 50% of the arm had a minor copy number of zero and as a post-WGD loss if at least 50% of the arm had a minor copy number of one.

This involves a weak assumption of parsimony as a minor copy number of zero could arise from two post-WGD losses, though without any other gains, a minor copy number of one cannot arise from a pre-WGD loss.

To correct for mutual exclusivity when measuring pre- and post-WGD losses, a corrected post-WGD loss proportion was calculated by dividing the post-WGD loss proportion by 1 – the pre-WGD loss proportion, thereby changing the denominator in proportion to only the samples that did not have a pre-WGD loss.

We also produced pre- and post-WGD gain and loss proportions at base pair resolution across the genome. Each segment was rounded to the nearest 1 kb. For certain cancer types with a low number of samples, the pre-WGD loss proportion was 1.0, leading to division by zero when calculating the corrected post-WGD loss proportion. Therefore, we clipped the pre-WGD proportion to a maximum of 0.95 when calculating the corrected post-WGD loss proportion. To compare relative rates, pan-genome event proportions were normalized such that the integral of the proportions over the genome was equal to 10^9^, chosen to provide a normalized event frequency with a magnitude of approximately one.

## Data availability

An access request for sequencing data and metadata from the Hartwig Medical Foundation can be found at https://www.hartwigmedicalfoundation.nl/en/data/data-acces-request/. Details of applications for PCAWG data are available at https://docs.icgc.org/pcawg/data.

## Code availability

GRITIC is available at https://github.com/VanLoo-lab/gritic.

## Supplementary data

**Supplementary Methods** – Additional description and justifications for the GRITIC method

**Supplementary Figures S1 to S40**

S1: WGD frequencies across cancer types and stage

S2: Effect of WGD constraint on timing accuracy

S3: Measuring timing accuracy on simulated data

S4-7: Measuring timing accuracy on simulated data by copy number state

S8-11: Measuring inferred route probabilities on simulated data

S12: Non-parsimony in copy number gain evolution

S13-14: Calibrating a penalty on non-parsimony

S15: Probability of pre-WGD gains in different chromosomes and copy number states

S16: Distribution of gain timing by major copy number

S17: Probability of first gain post-WGD by copy number state

S18: Single copy number profiles of an undifferentiated sarcoma

S19: Distribution of gain rates relative to WGD by cancer type

S20: Combined distribution over gain timing by WGD status

S21: The timing of gains relative to WGD

S22: The timing of gains relative to WGD by cancer type

S23: The relationship between genome gained post-WGD and WGD timing by cancer type

S24: The relationship between genome gained pre-WGD and WGD timing by cancer type

S25: The relationship between the fraction of genome lost pre and post-WGD and WGD timing by cancer type

S26: Punctuated gains in WGD tumors

S27: Frequency of arm gains pre and post-WGD and in non-WGD tumors

S28: Frequency of arm gains pre and post-WGD and in non-WGD tumors by cancer type

S29: Frequency of arm losses pre and post-WGD and in non-WGD tumors by cancer type

S30: Effect of oncogene and tumor suppressor gene density on arm gain rates

S31: Effect of oncogene and tumor suppressor gene density on arm loss rates

S32-36: Pan-genome frequencies of pre and post-WGD gains by cancer type S37-39: Pan-genome frequencies of pre and post-WGD losses by cancer type S40: WGD status calling in GRITIC

## Acknowledgments

This work was supported by the Francis Crick Institute which receives its core funding from Cancer Research UK (CC2008), the UK Medical Research Council (CC2008), and the Wellcome Trust (CC2008). For the purpose of Open Access, the authors have applied a CC BY public copyright license to any Author Accepted Manuscript version arising from this submission. T.M.B. was supported by a PhD fellowship from Boehringer Ingelheim Fonds. A.R.L. is a TRIUMPH Fellow in the CPRIT Training Program (RP210028). M.T. was supported as a postdoctoral researcher of the F.R.S.-FNRS. C.S. is Royal Society Napier Research Professor (RP150154). His work is supported by the Francis Crick Institute, which receives its core funding from Cancer Research UK (CC2041), the UK Medical Research Council (CC2041), and the Wellcome Trust (CC2041). C.S. is funded by Cancer Research UK (TRACERx, C11496/A17786; Cancer Research UK Lung Cancer Centre of Excellence C11496/A30025), the Rosetrees Trust, Butterfield and Stoneygate Trusts, NovoNordisk Foundation (ID16584), a Royal Society Research Professorship Enhancement Award (RP/EA/180007), the National Institute for Health Research (NIHR) Biomedical Research Centre at University College London Hospitals, the Breast Cancer Research Foundation (BCRF 20-157), the CRUK-UCL Centre, Experimental Cancer Medicine Centre. His research is supported by a Stand Up To Cancer-LUNGevity-American Lung Association Lung Cancer Interception Dream Team Translational Research Grant (SU2C-AACR-DT23-17). C.S. also receives funding from the European Research Council (FP7-THESEUS-617844, FP7-PloidyNet 607722, PROTEUS 835297 and Chromavision 665233). P.V.L. is a Winton Group Leader in recognition of the Winton Charitable Foundation’s support towards the establishment of The Francis Crick Institute. P.V.L. is a CPRIT Scholar in Cancer Research and acknowledges CPRIT grant support (RR210006). The results shown here are in part based upon data generated by the TCGA Research Network: https://www.cancer.gov/tcga. This publication and the underlying study have been made possible partly based on data that the Hartwig Medical Foundation and the Center of Personalised Cancer Treatment (CPCT) have made available to the study through the Hartwig Medical Database. We thank Paul Spellman for his helpful comments on our manuscript.

## Author contributions

T.M.B. conceptualized and developed GRITIC under the supervision of M.T. and P.V.L.; T.M.B. carried out the majority of the analyses; S.L. conducted the punctuated gains analysis; T.L. and S.D. undertook the SNV clustering analysis on Hartwig. H.Y. and A.L.B. performed the computational single-cell analysis; A.V. performed wet-lab experiments on undifferentiated sarcomas. T.M.B., M.T., and P.V.L. wrote the manuscript with contributions from A.R.L.; N.P., A.M.F., and C.S. provided samples and expertise and contributed to manuscript writing. All authors have read and approved the final manuscript.

