## Supplementary Figures for "The history of chromosomal instability in genome doubled tumors"

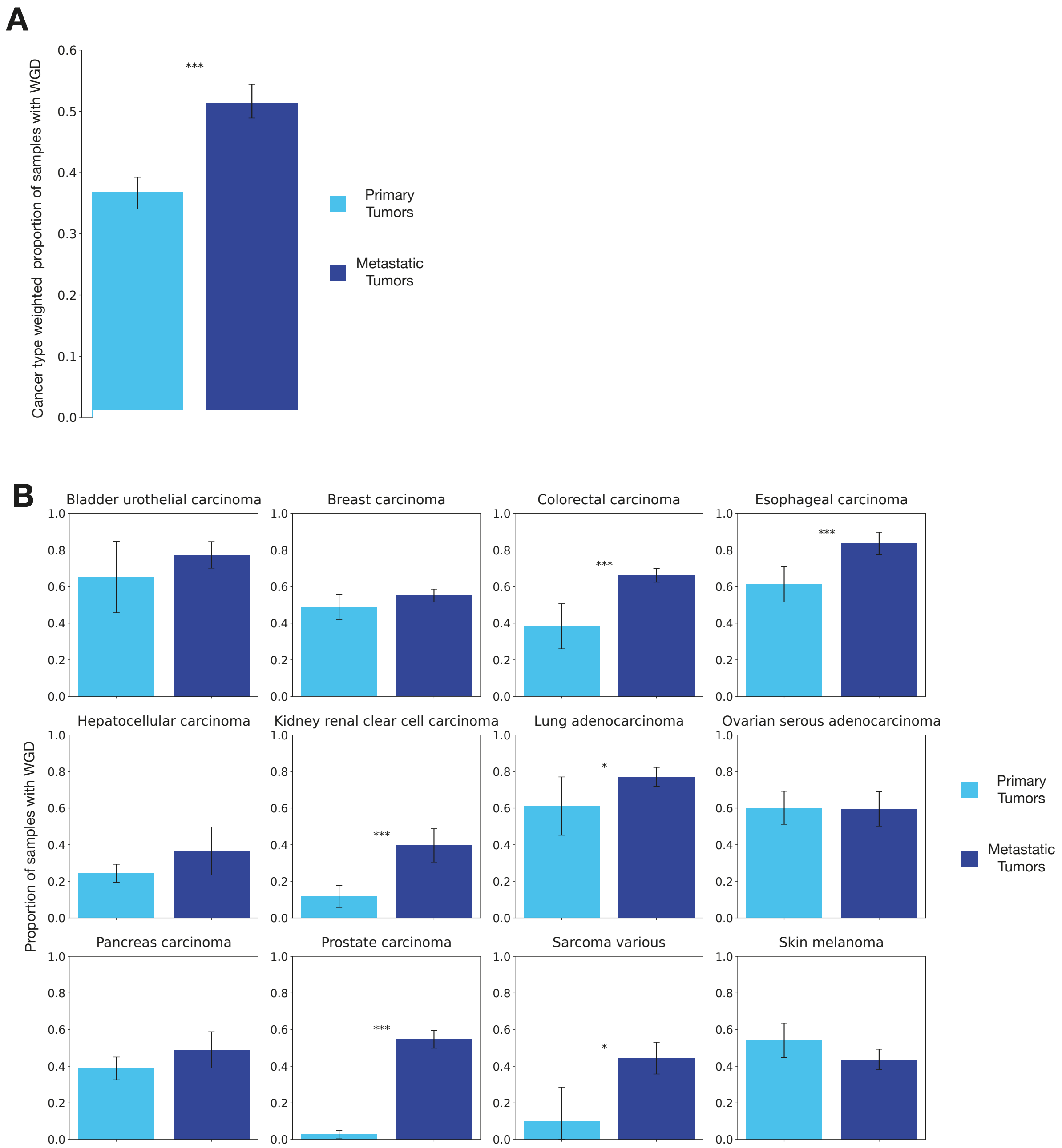


**Supplementary Figure 1. WGD frequencies across cancer types and stage. A,** The proportion of tumor samples identified as WGD in primary and metastatic cohorts, weighted by cancer type. Only primary tumors with at least 20 primary and metastatic tumors are included. Statistical significance was calculated by proportion test and 95% confidence intervals by bootstrapping over samples. **B,** The proportion of tumor samples identified as WGD in primary and metastatic cohorts across different primary cancer types. Statistical significance was calculated by proportion test and 95% confidence intervals by normal approximation.

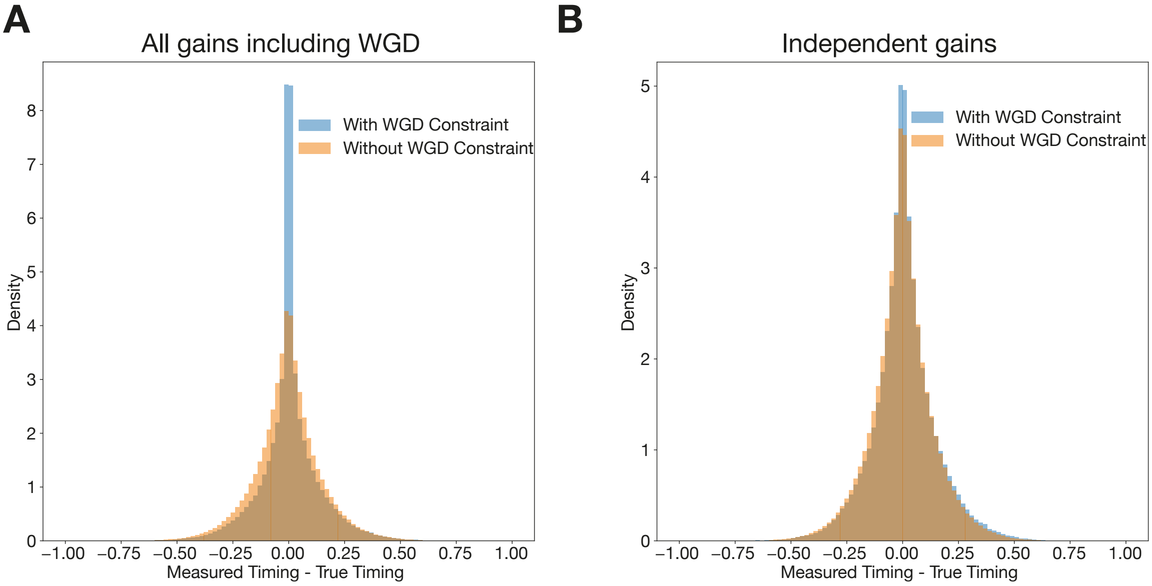


**Supplementary Figure 2. Effect of WGD constraint on timing accuracy. A,** The difference between the measured posterior timing distribution and the true timing for a cohort of representative simulated tumor gains in a WGD cohort. All copy number gains in complex states including those arising through a WGD are included in the distribution. **B,** The difference between the measured posterior timing distribution and the true timing for a cohort of representative simulated tumor gains in a WGD cohort. Only the timing of copy number gains in complex states that arose independently of the WGD are included.

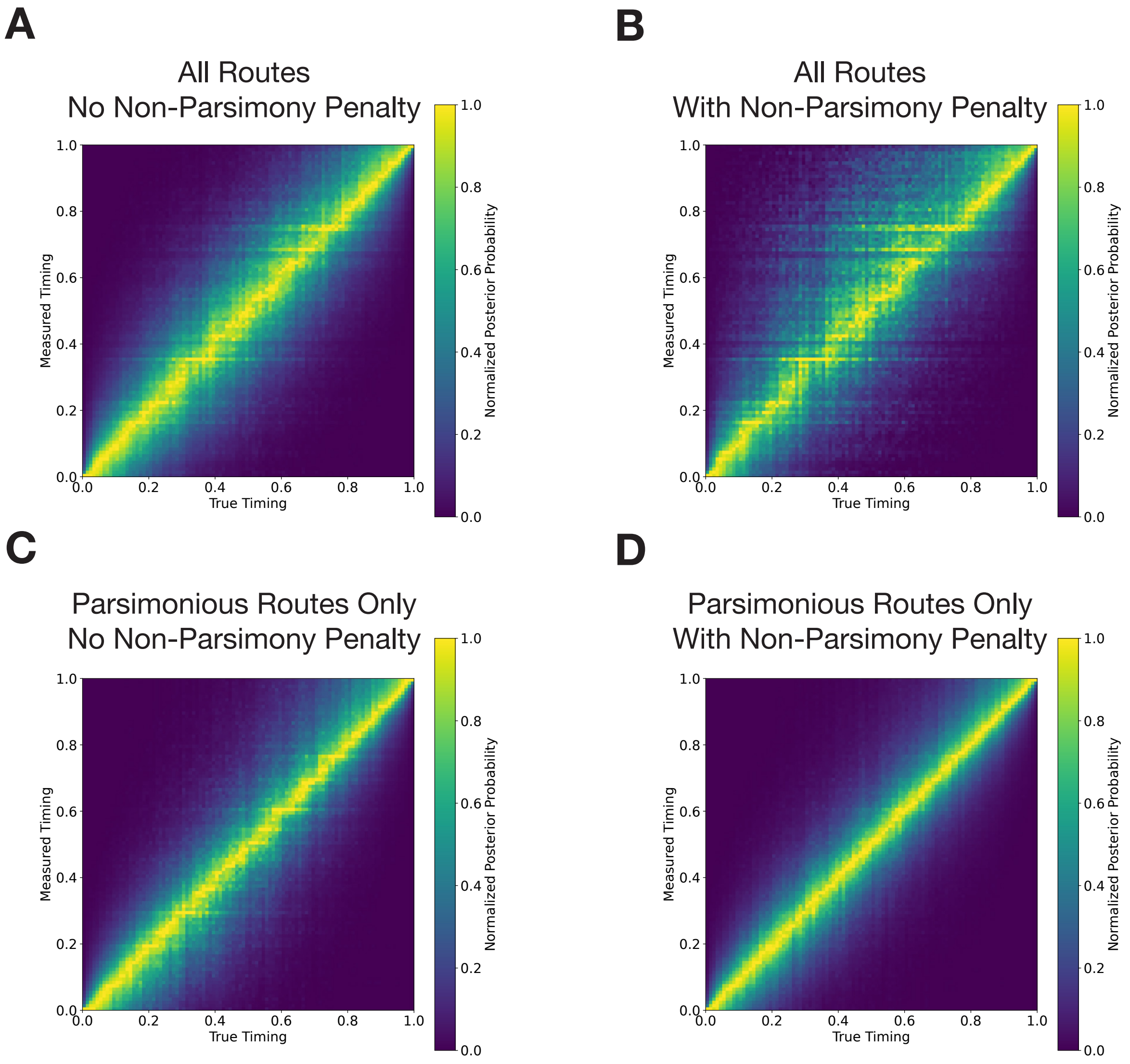


**Supplementary Figure 3. Measuring timing accuracy on simulated data.** Distribution of measured posterior probability on gain timing against true gain timing across several representative cohorts of complex gains simulated under different conditions. The histograms are normalized such that the posterior probability is set relative to the maximum posterior probability for each value for true timing.

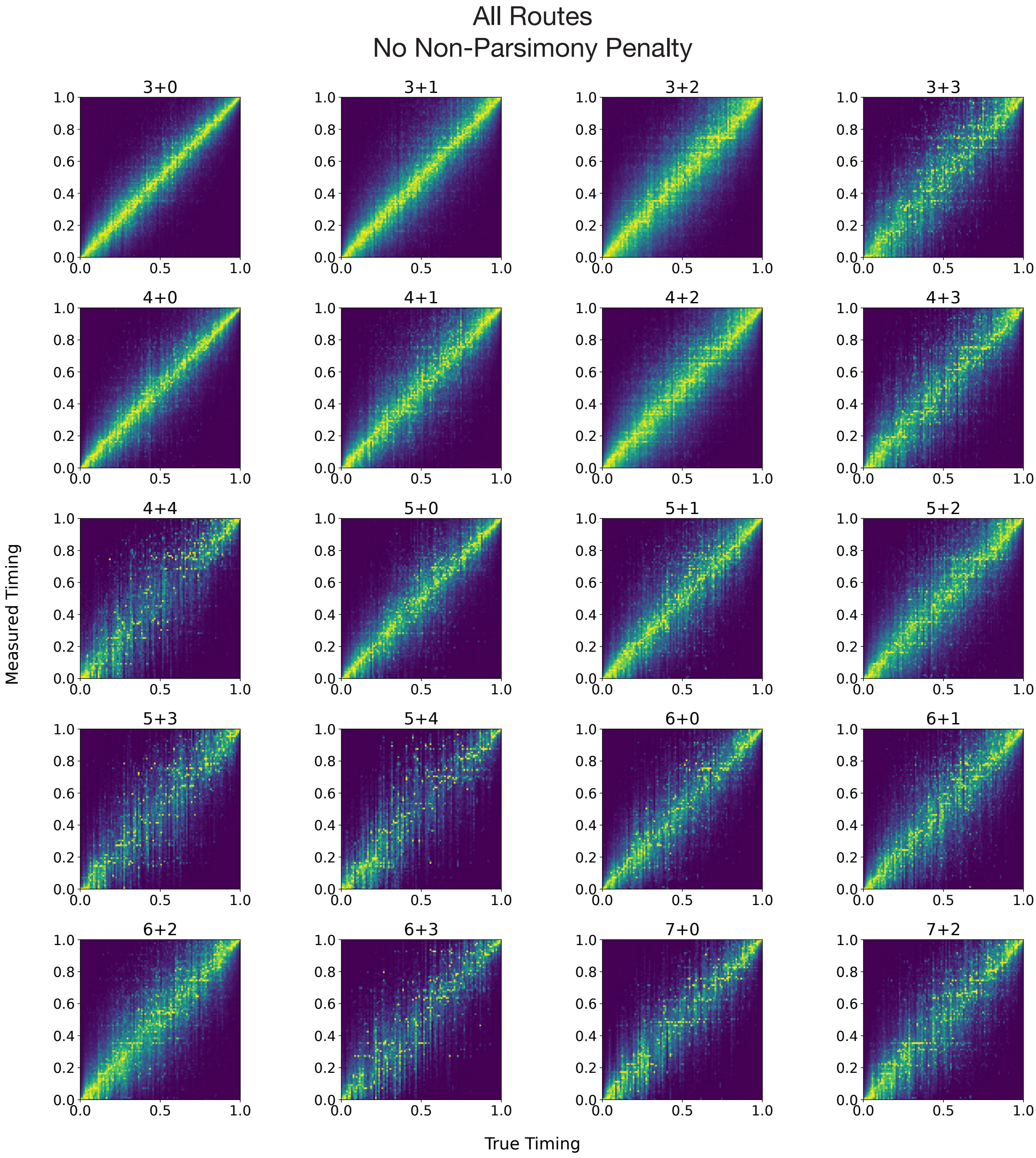


**Supplementary Figure 4. Measuring timing accuracy on simulated data by copy number state.** Distribution of measured posterior probability on gain timing against true gain timing for a representative cohort of tumors where all gains from all routes are simulated as the ground truth, and no penalty is applied to non-parsimonious routes during inference. The histograms are normalized such that the posterior probability is set relative to the maximum posterior probability for each value for true timing.

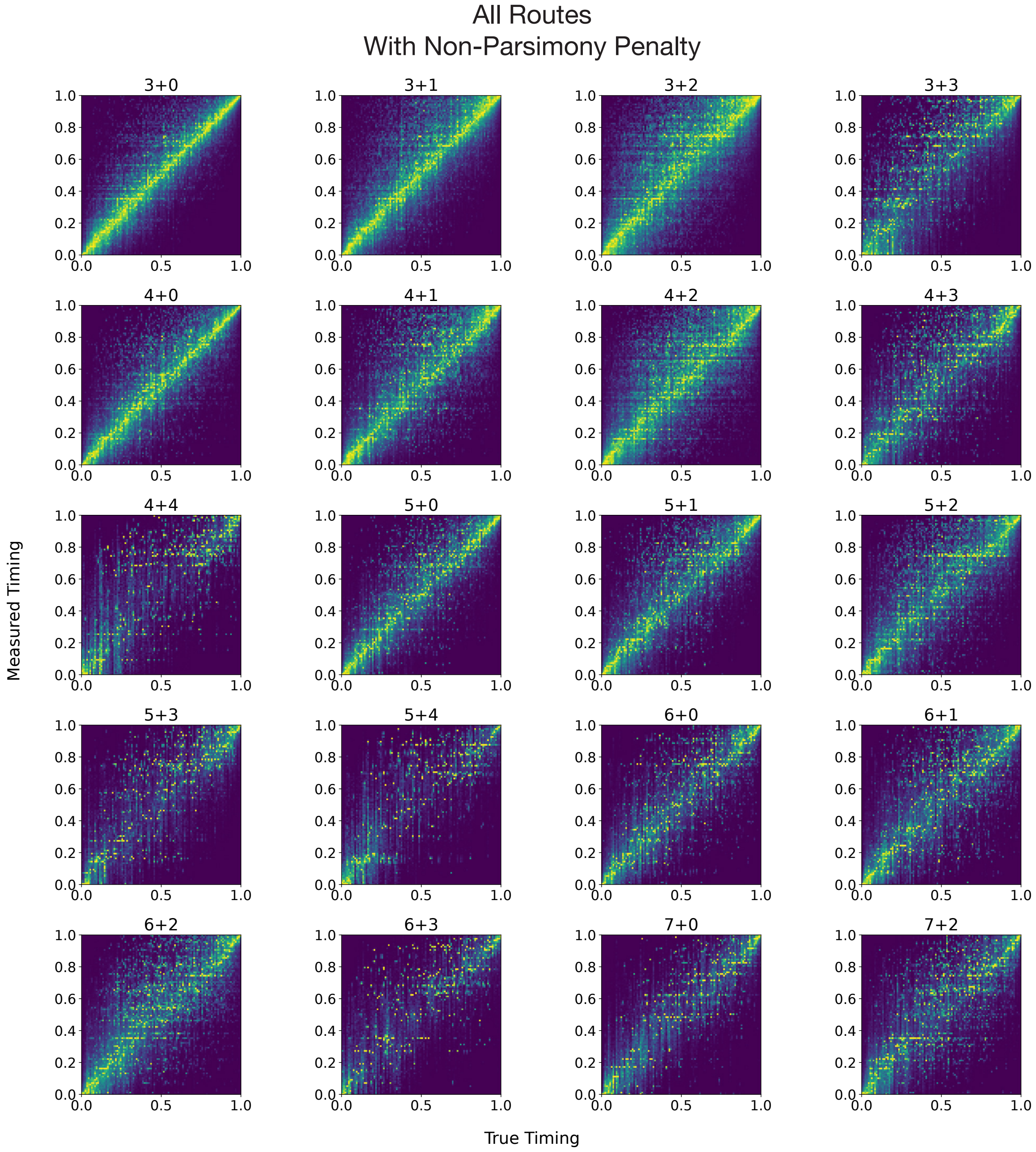


**Supplementary Figure 5. Measuring timing accuracy on simulated data by copy number state.** Distribution of measured posterior probability on gain timing against true gain timing for a representative cohort of tumors where all gains from all routes are simulated as the ground truth, and a penalty is applied to non-parsimonious routes during inference (see Methods). The histograms are normalized such that the posterior probability is set relative to the maximum posterior probability for each value for true timing.

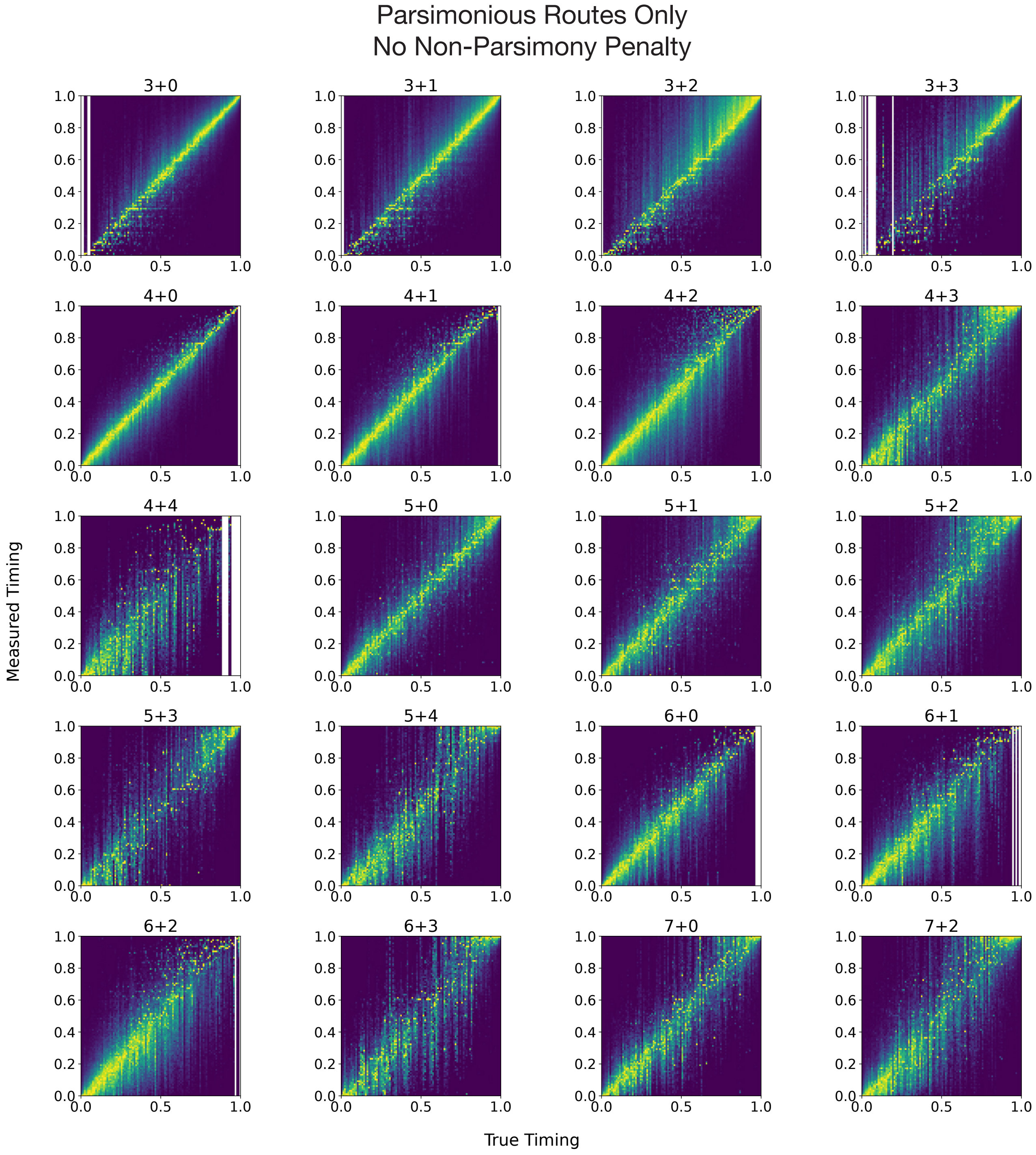


**Supplementary Figure 6. Measuring timing accuracy on simulated data by copy number state.** Distribution of measured posterior probability on gain timing against true gain timing for a representative cohort of tumors where all gains from solely parsimonious routes are simulated as the ground truth, with no penalty applied to non-parsimonious routes during inference. The histograms are normalized such that the posterior probability is set relative to the maximum posterior probability for each value for true timing. True timing values where there was insufficient data for inference are indicated by white columns.

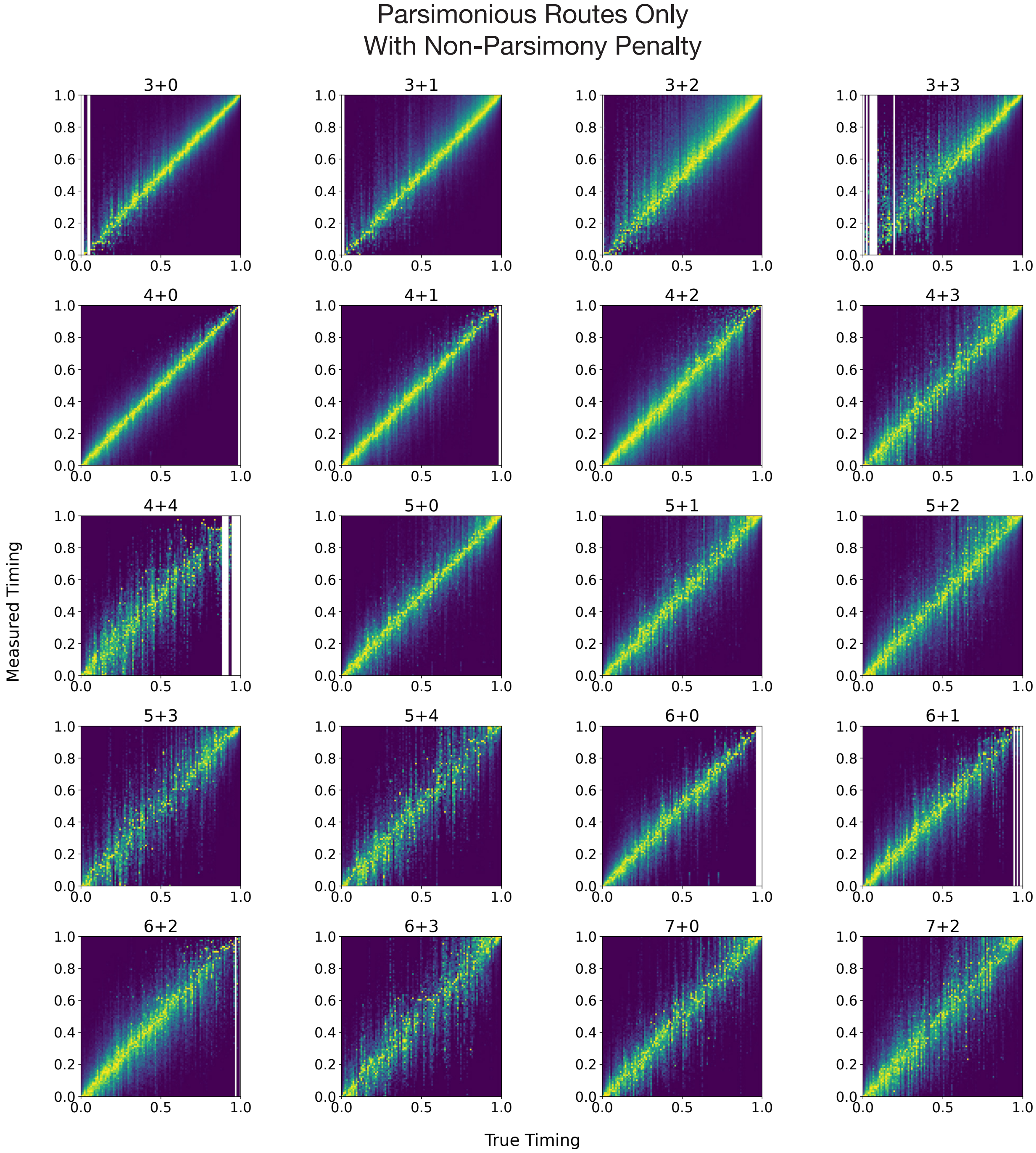


**Supplementary Figure 7. Measuring timing accuracy on simulated data by copy number state.** Distribution of measured posterior probability on gain timing against true gain timing for a representative cohort of tumors where all gains from solely parsimonious routes are simulated as the ground truth, with a penalty applied to non-parsimonious routes during inference (see Methods). The histograms are normalized such that the posterior probability is set relative to the maximum posterior probability for each value for true timing. True timing values where there was insufficient data for inference are indicated by white columns.

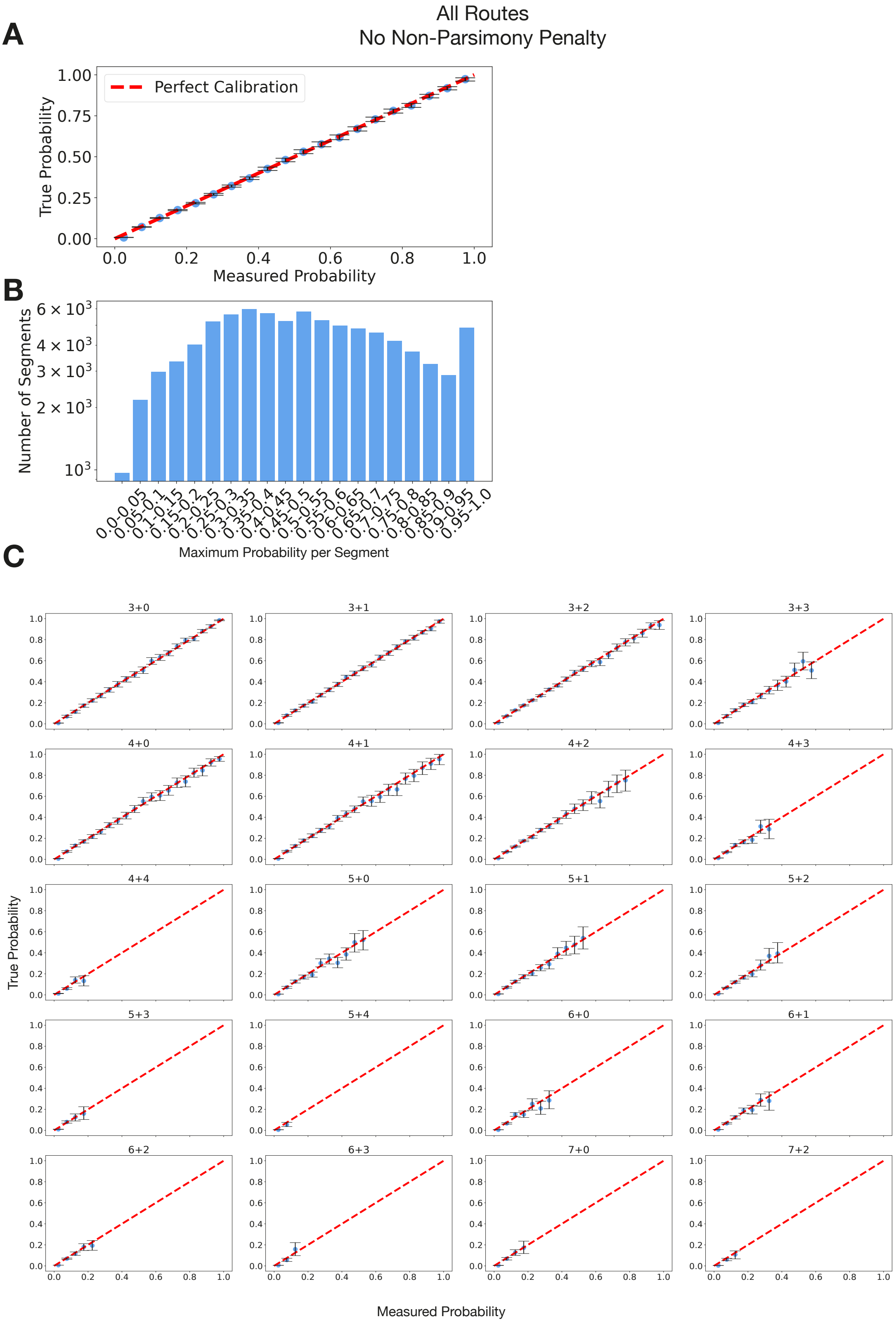


**Supplementary Figure 8. Measuring inferred route probabilities on simulated data.** Probability calibration plots for a cohort simulated with all routes and no penalty on non-parsimony applied during inference. **A,** Binned measured probability of different route assignments against true probability calculated as the proportion of segments within each bin of measured probability that have the route assignment corresponding to the probability. 95% confidence intervals were calculated by bootstrapping over samples. **B,** Distribution of the maximum probability across all routes for the segments in the simulated cohort. **C,** Binned measured probability of different route assignments against true probability calculated as the proportion of segments within each bin of measured probability that have the route assignment corresponding to the probability, split by copy number state. 95% confidence intervals were calculated by bootstrapping over samples.

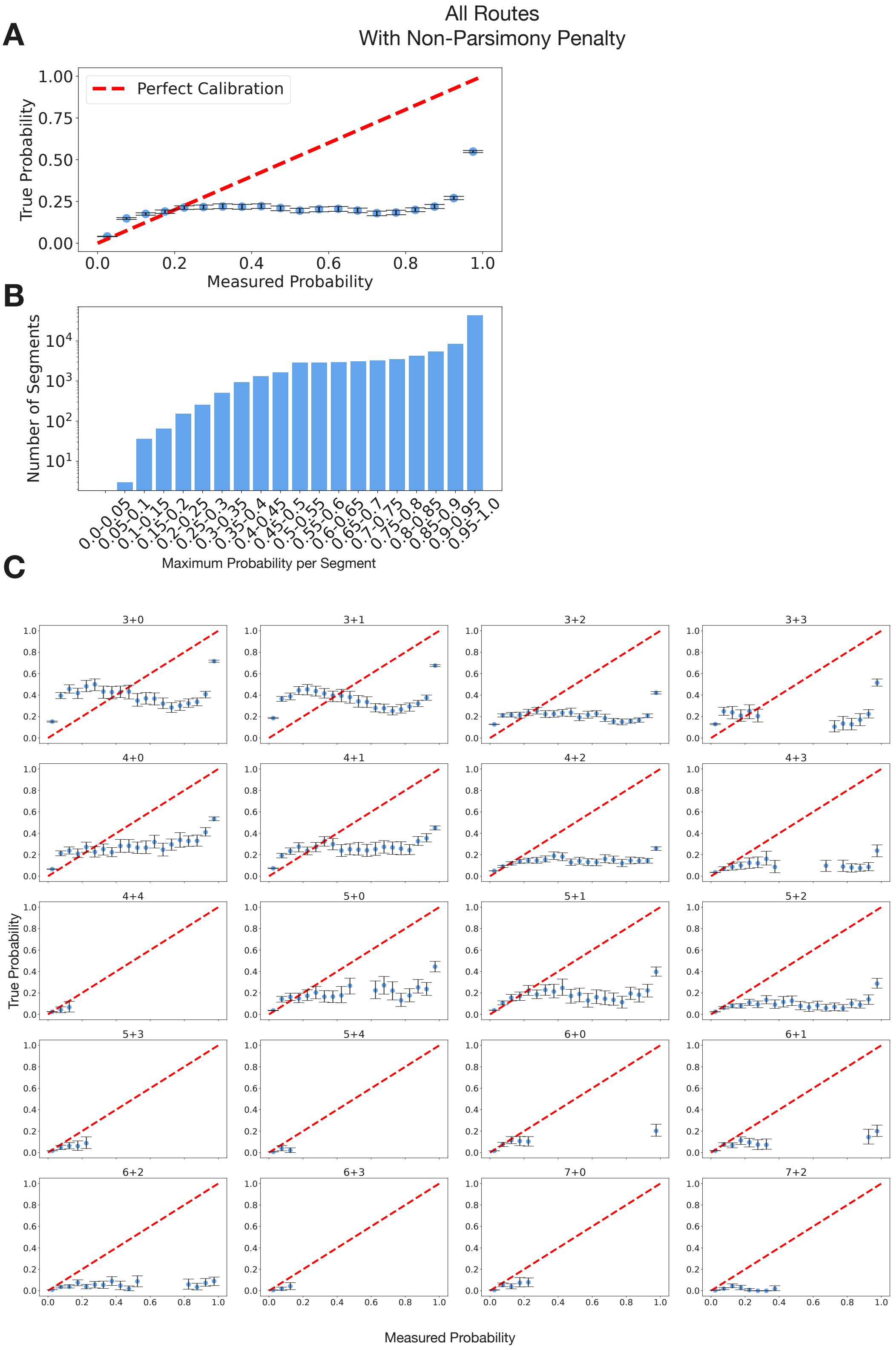


**Supplementary Figure 9. Measuring inferred route probabilities on simulated data.** Probability calibration plots for a cohort simulated with all routes and a penalty on non-parsimony applied during inference. **A,** Binned measured probability of different route assignments against true probability calculated as the proportion of segments within each bin of measured probability that have the route assignment corresponding to the probability. 95% confidence intervals were calculated by bootstrapping over samples. **B,** Distribution of the maximum probability across all routes for the segments in the simulated cohort. **C,** Binned measured probability of different route assignments against true probability calculated as the proportion of segments within each bin of measured probability that have the route assignment corresponding to the probability, split by copy number state. 95% confidence intervals were calculated by bootstrapping over samples.

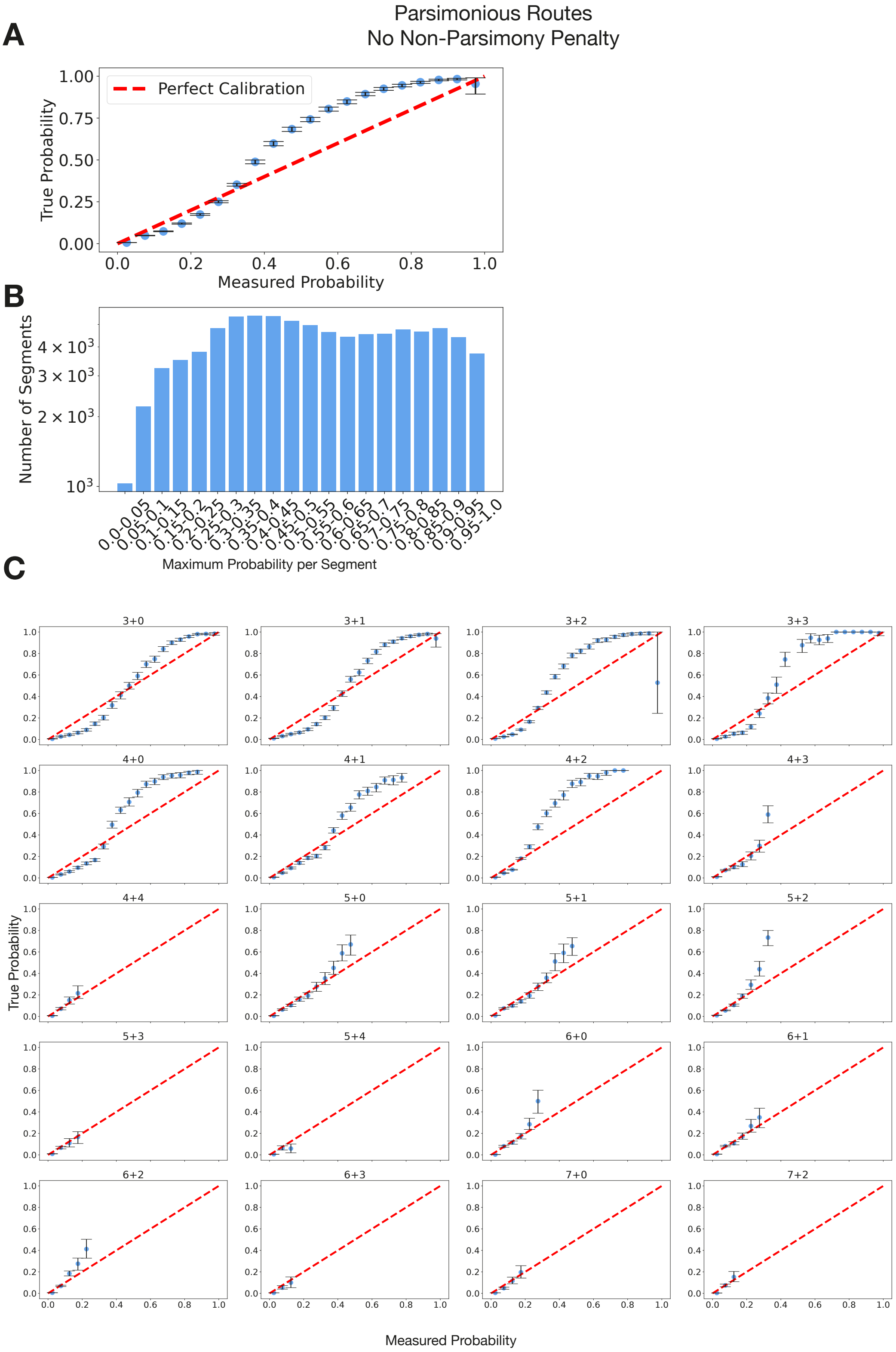


**Supplementary Figure 10. Measuring inferred route probabilities on simulated data.** Probability calibration plots for a cohort simulated with parsimonious routes only and no penalty on non-parsimony applied during inference. **A,** Binned measured probability of different route assignments against true probability calculated as the proportion of segments within each bin of measured probability that have the route assignment corresponding to the probability. 95% confidence intervals were calculated by bootstrapping over samples. **B,** Distribution of the maximum probability across all routes for the segments in the simulated cohort. **C,** Binned measured probability of different route assignments against true probability calculated as the proportion of segments within each bin of measured probability that have the route assignment corresponding to the probability, split by copy number state. 95% confidence intervals were calculated by bootstrapping over samples.

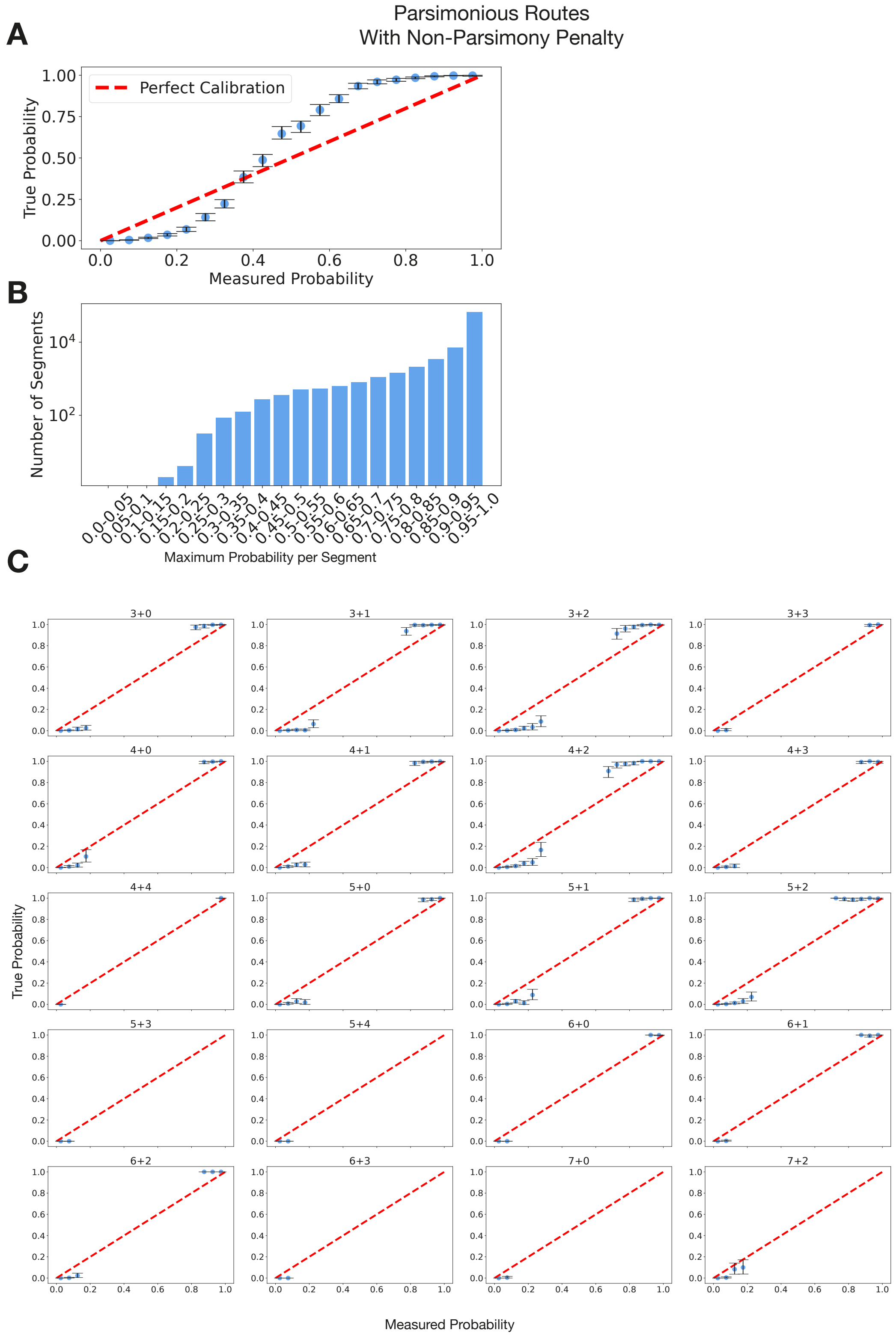


**Supplementary Figure 11. Measuring inferred route probabilities on simulated data.** Probability calibration plots for a cohort simulated with parsimonious routes only and a penalty on non-parsimony applied during inference. **A,** Binned measured probability of different route assignments against true probability calculated as the proportion of segments within each bin of measured probability that have the route assignment corresponding to the probability. 95% confidence intervals were calculated by bootstrapping over samples. **B,** Distribution of the maximum probability across all routes for the segments in the simulated cohort. **C,** Binned measured probability of different route assignments against true probability calculated as the proportion of segments within each bin of measured probability that have the route assignment corresponding to the probability, split by copy number state. 95% confidence intervals were calculated by bootstrapping over samples.

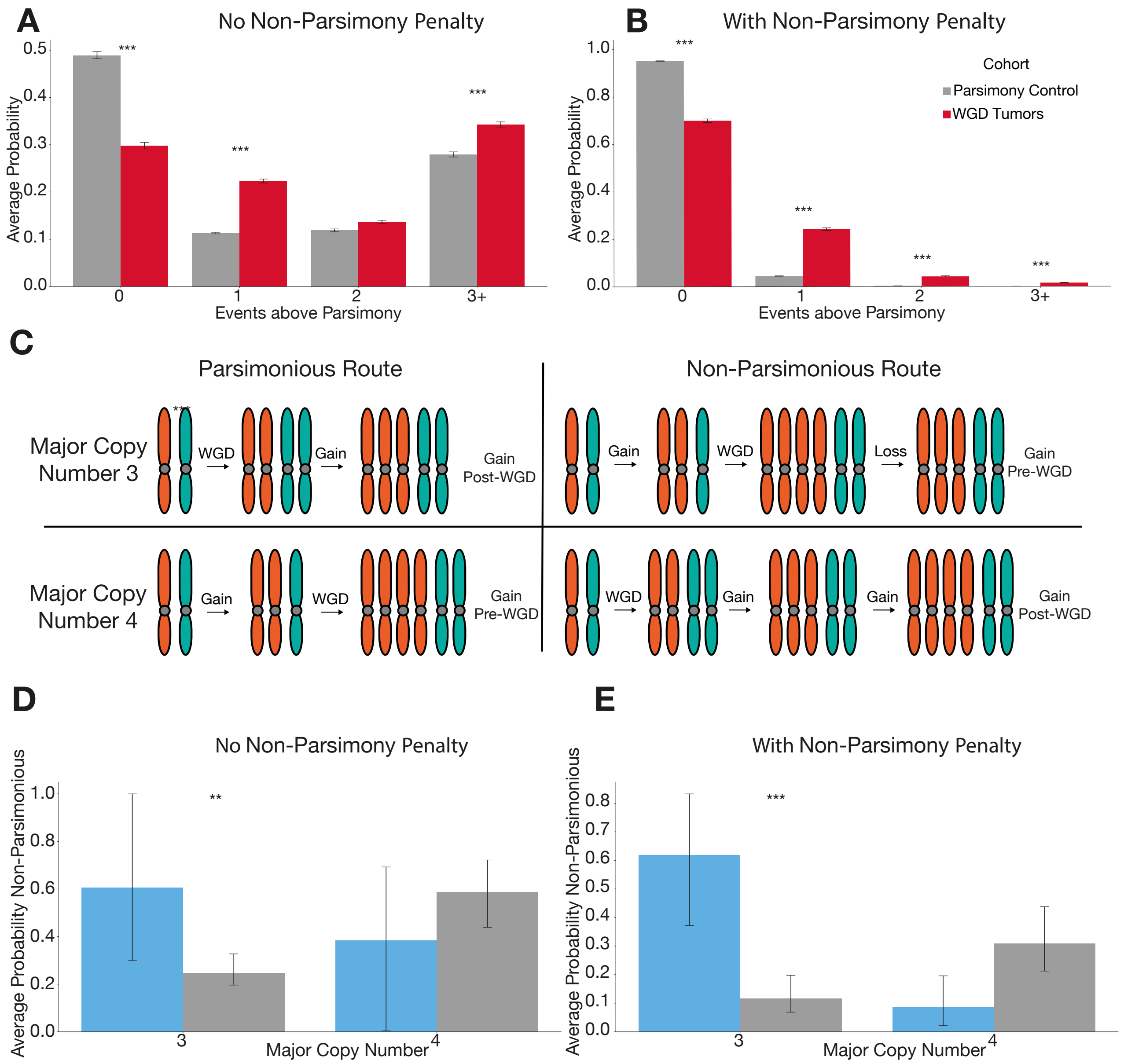


**Supplementary Figure 12. Non-parsimony in copy number gain evolution. A,B,** The average posterior probability on number of additional events required to reach the final state over the most parsimonious route for complex gained states in the PCAWG and Hartwig cohort compared to a simulated control where only parsimonious routes were included. Measured with **(A)** and without a penalty **(B)** on non-parsimony during inference. Statistical significance is calculated using a permutation test and 95% confidence intervals by bootstrapping over samples. **C**, Schematic demonstrating that parsimonious routes have an earlier independent gain for major copy number three copy number segments in WGD tumors and vice versa for major copy number four segments. **D-E,** The average probability on non-parsimonious routes for gained segments in kidney renal cell carcinoma split by major copy number and gain location with **(D)** and without a penalty **(E)** on non-parsimony during inference. Statistical significance is calculated using a permutation test and 95% confidence intervals by bootstrapping over samples.

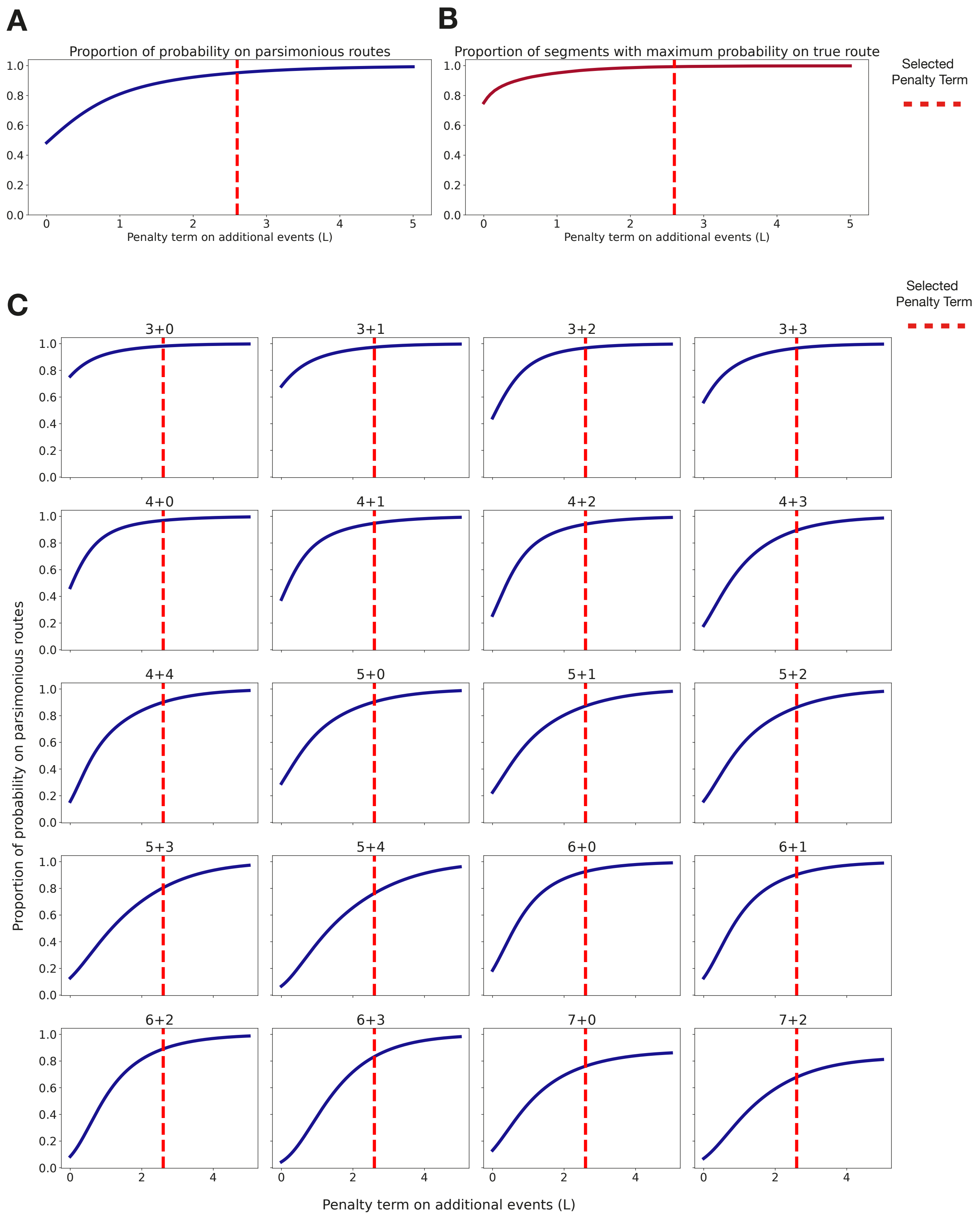


**Supplementary Figure 13. Calibrating a penalty on non-parsimony. A,** The average probability assigned to parsimonious routes on a representative simulated cohort of parsimonious only copy number routes against a non-parsimony penalty term. The non-parsimony penalty is exp(-NL) where N is the number of extra events over the minimum number of possible events in all possible routes for the segment. **B,** The average number of segments with maximum a- posteriori probability assigned to parsimonious routes on a representative simulated cohort of parsimonious only copy number routes against a non-parsimony penalty term. **C,** The average probability assigned to parsimonious routes on a representative simulated cohort of parsimonious only copy number routes against a non-parsimony penalty term, split by copy number state.

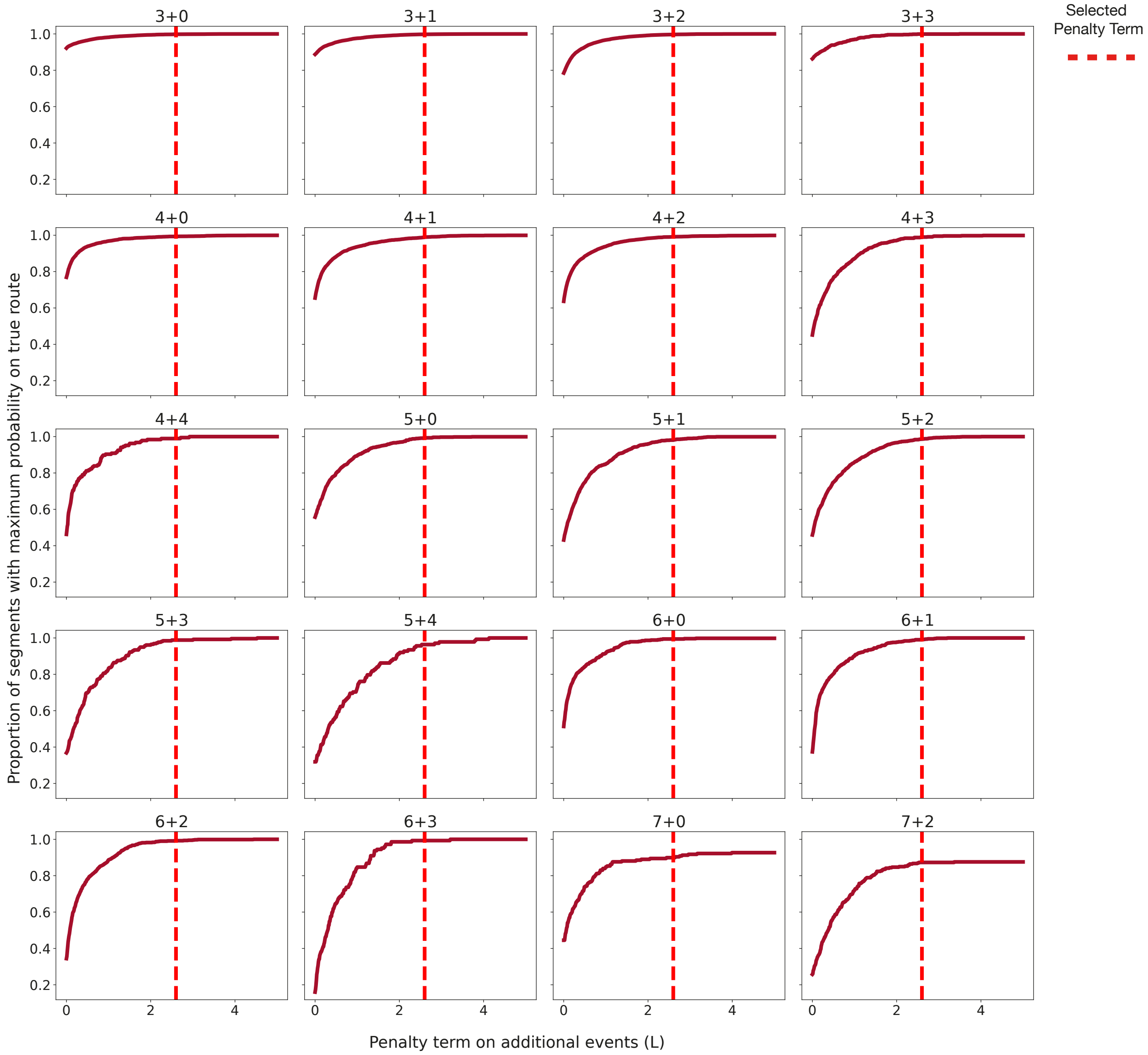


**Supplementary Figure 14. Calibrating a penalty on non-parsimony.** The average number of segments with maximum a-posteriori probability assigned to parsimonious routes on a representative simulated cohort of parsimonious only copy number routes against a non-parsimony penalty term, split by copy number state. The non-parsimony penalty is exp(-NL) where N is the number of extra events over the minimum number of possible events in all possible routes for the segment.

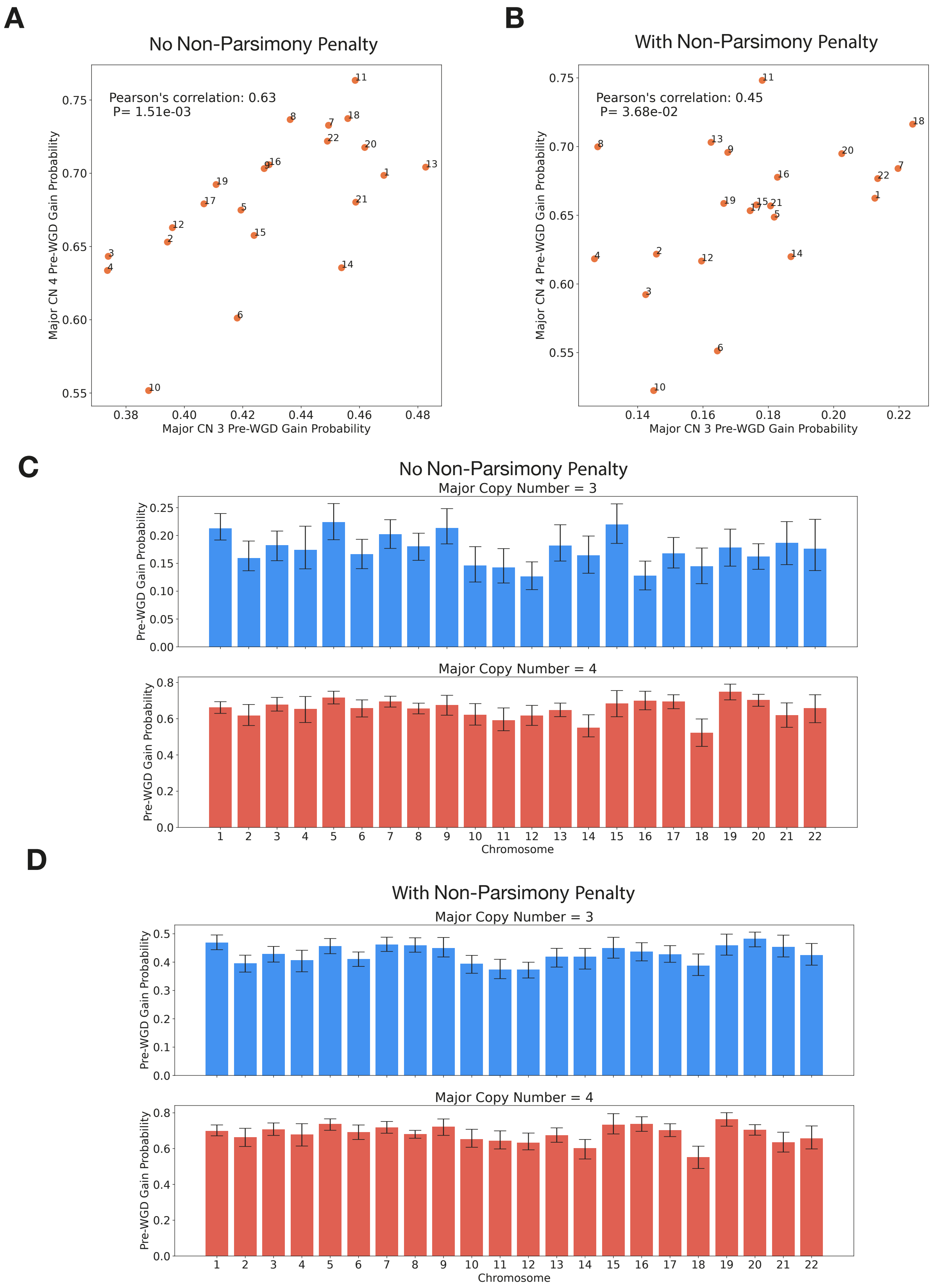


**Supplementary Figure 15**. **Probability of pre-WGD gains in different chromosomes and copy number states. A,B** The average probability the first gain arises pre-WGD in different chromosomes in major copy number 3 and 4 states as measured the PCAWG and Hartwig datasets. Measured without **(A)** and with **(B)** a penalty on non-parsimony. **C,** The average probability the first gain arises pre-WGD in different chromosomes split by major copy number in the PCAWG and Hartwig datasets. Measured without **(C)** and with **(D)** a penalty on non-parsimony. 95% confidence intervals were calculated by bootstrapping over samples.

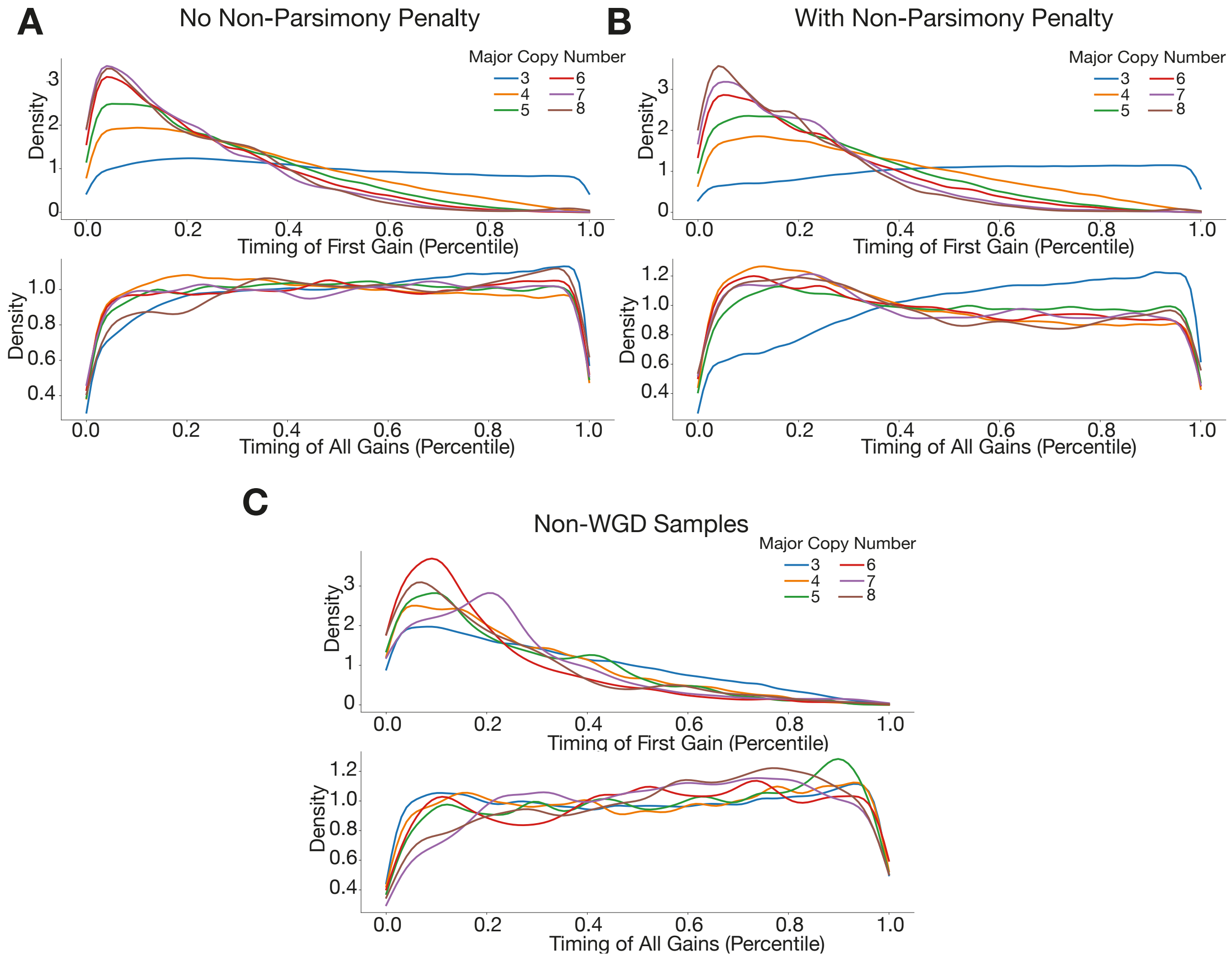


**Supplementary Figure 16. Distribution of gain timing by major copy number. A-C.** The distribution of the percentile timing within samples of the first gains and all gains in complex segments relative to other gains in the same sample, split by major copy number, with **(A)** and without a penalty **(B)** on non-parsimony during inference in WGD samples and **(C)** in non-WGD samples.

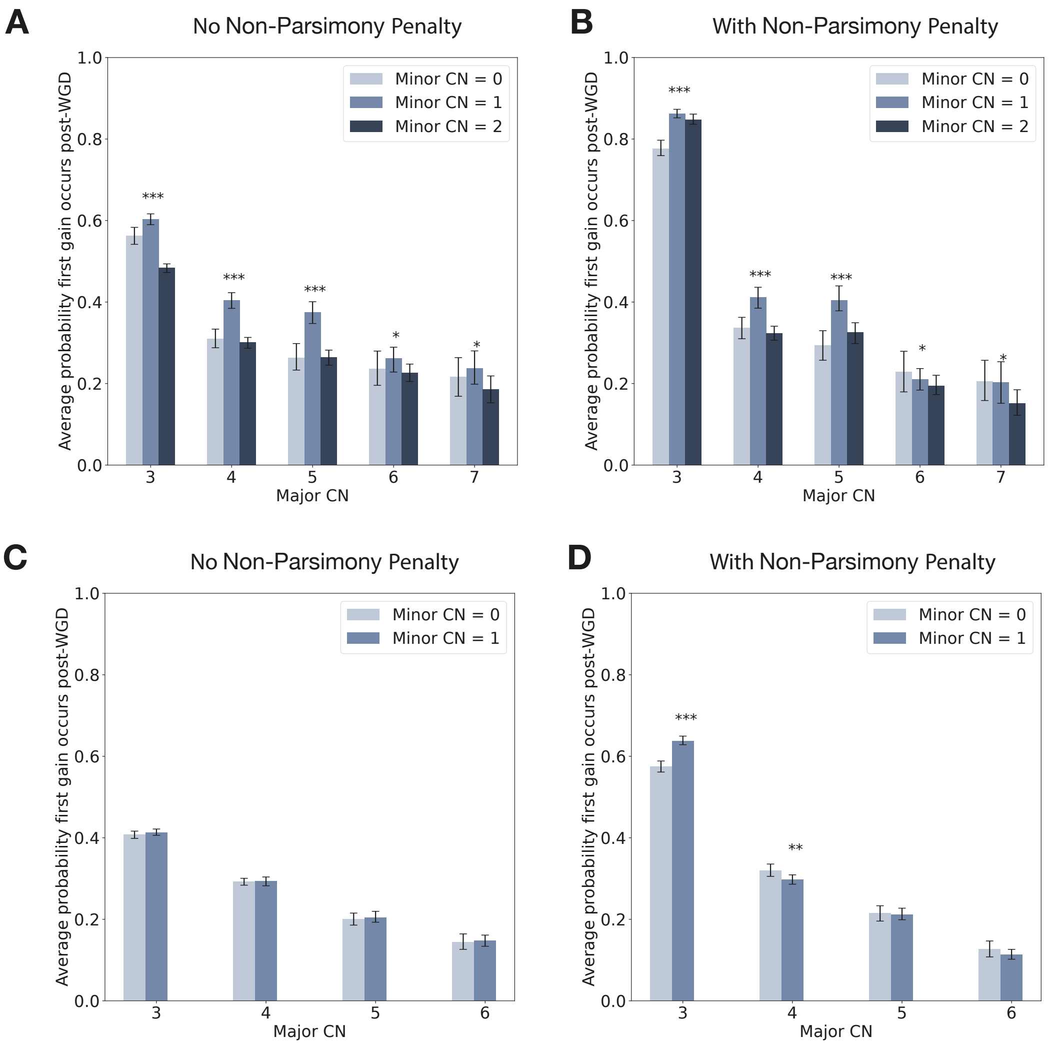


**Supplementary Figure 17. Probability of first gain post-WGD by copy number state**. **A-B,** The average probability of all gains occurring post-WGD weighted by segment width for different complex copy number states with (A) and without a penalty (B) on non-parsimony during inference on the PCAWG and Hartwig cohorts. Statistical significance was calculated by bootstrapping. **C-D,** The average probability of all gains occurring post-WGD weighted by segment width for different complex copy number states with (C) and without a penalty (D) on non-parsimony during inference on a representative simulated cohort. The cohort was simulated with equal likelihood of post-WGD gains for all minor copy number states for a given major copy number. States with minor CN two were excluded as they were simulated with a greater number of pre-WGD gains than the other minor copy number states. Statistical significance is calculated using a permutation test and 95% confidence intervals by bootstrapping over samples.

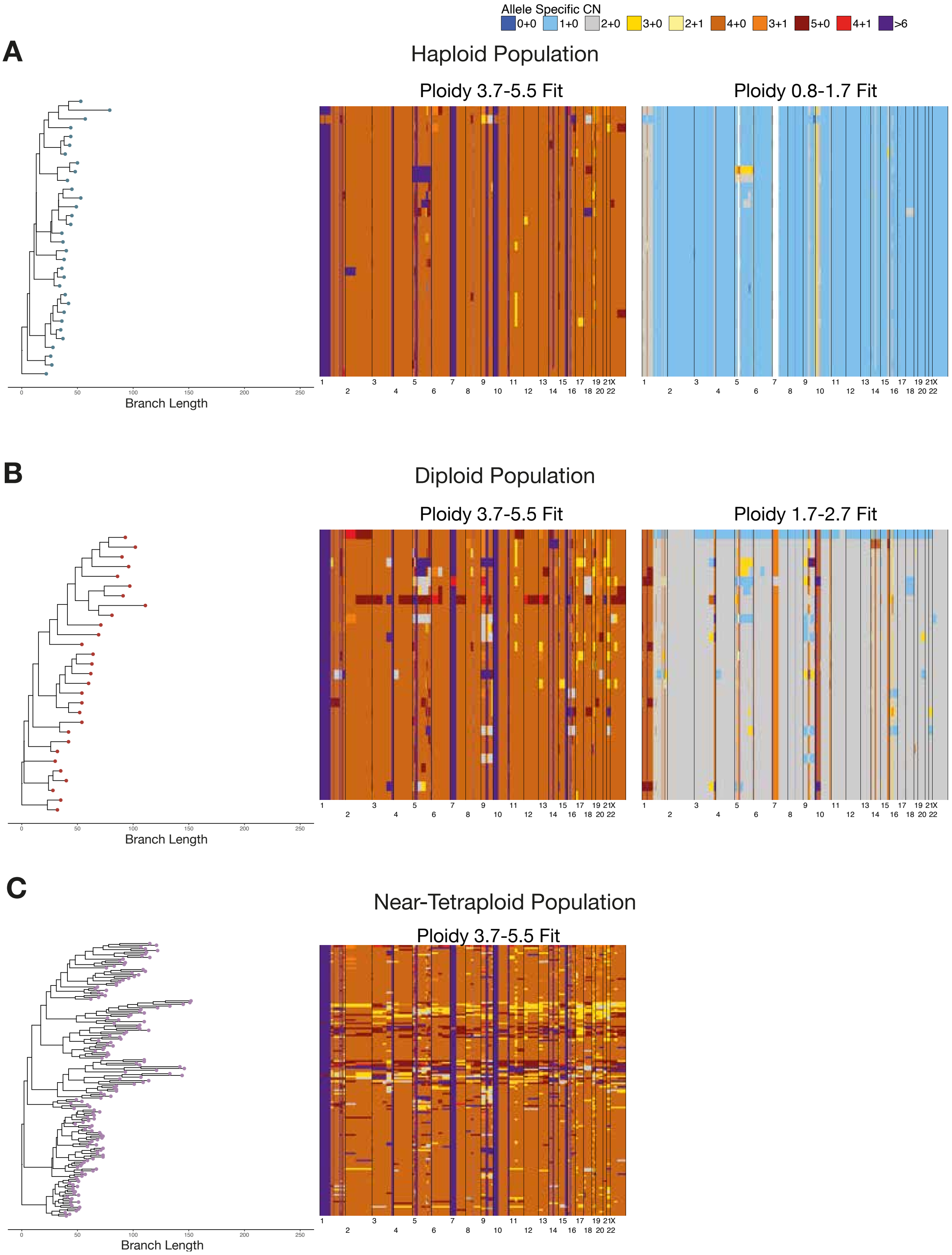


**Supplementary Figure 18. Single copy number profiles of an undifferentiated sarcoma** Single-cell copy number phylogenies and associated allele-specific copy number profiles from an individual undifferentiated sarcoma. These are split by different ploidy populations within the same tumor (**A-C**). One set of copy number profiles is fitted to true tumor cell ploidy as determined by fluorescence-activated cell sorting, the second is fitted to a near-tetraploid state to get an unbiased estimate of copy number heterogeneity.

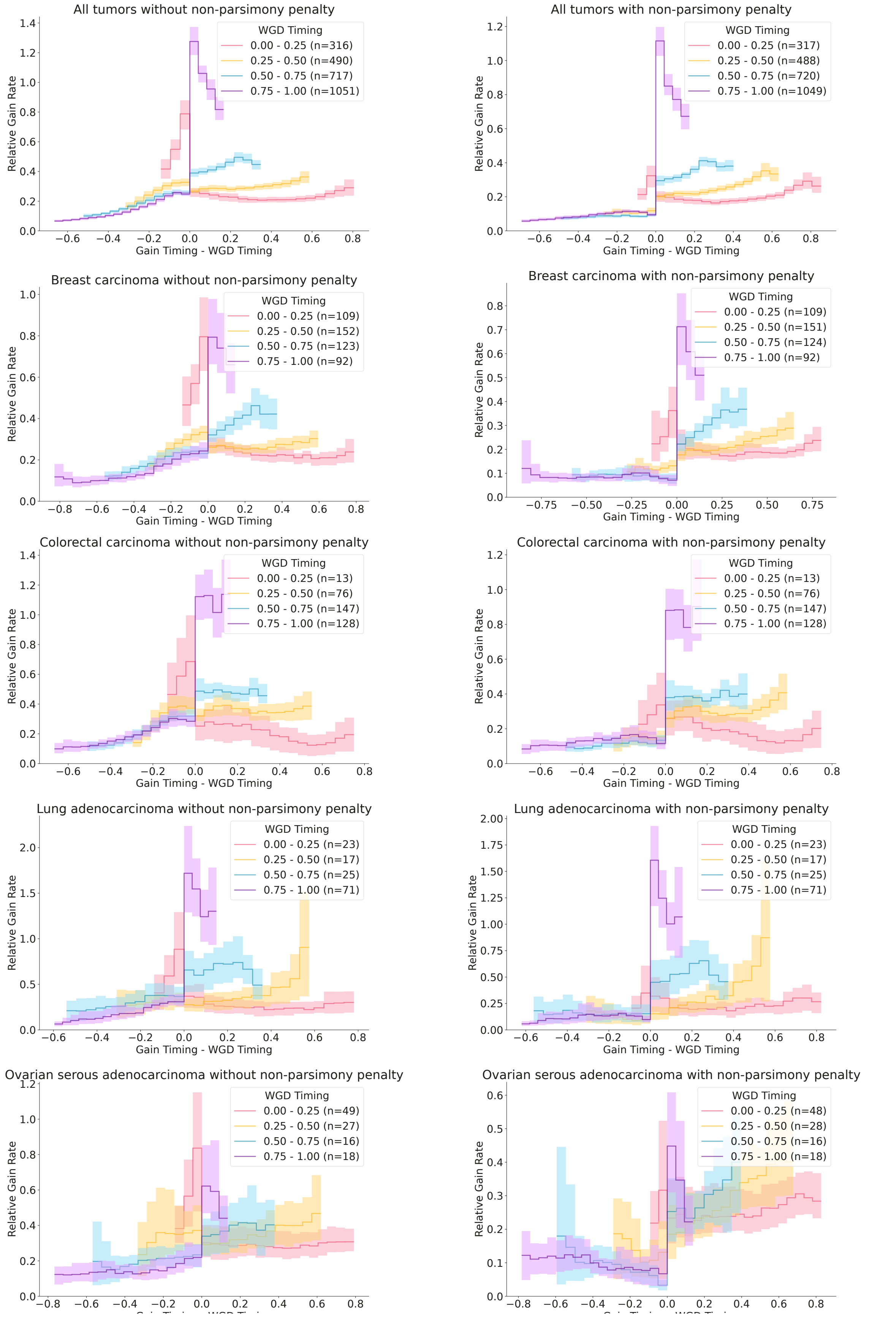


**Supplementary Figure 19. Distribution of gain rates relative to WGD by cancer type**. The normalized rate of gains relative to the WGD timing for a selection of cancer types with and without a parsimony penalty applied during inference. 95% confidence intervals are calculated by bootstrapping over samples.

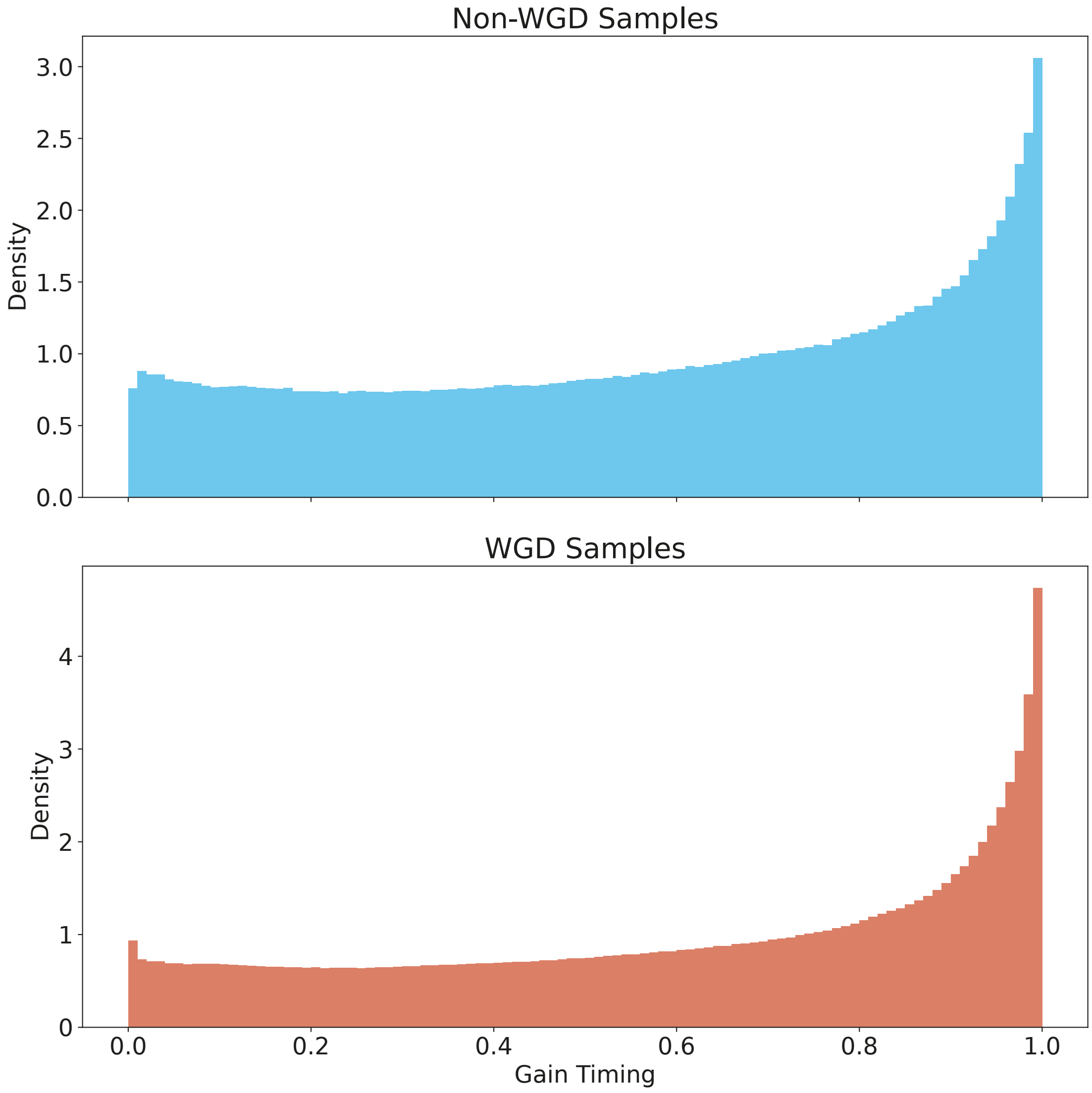


**Supplementary Figure 20. Combined distribution over gain timing by WGD status.** Distribution of gain timing across all segments and samples for non-WGD and WGD samples. Inference without penalty on non-parsimony.

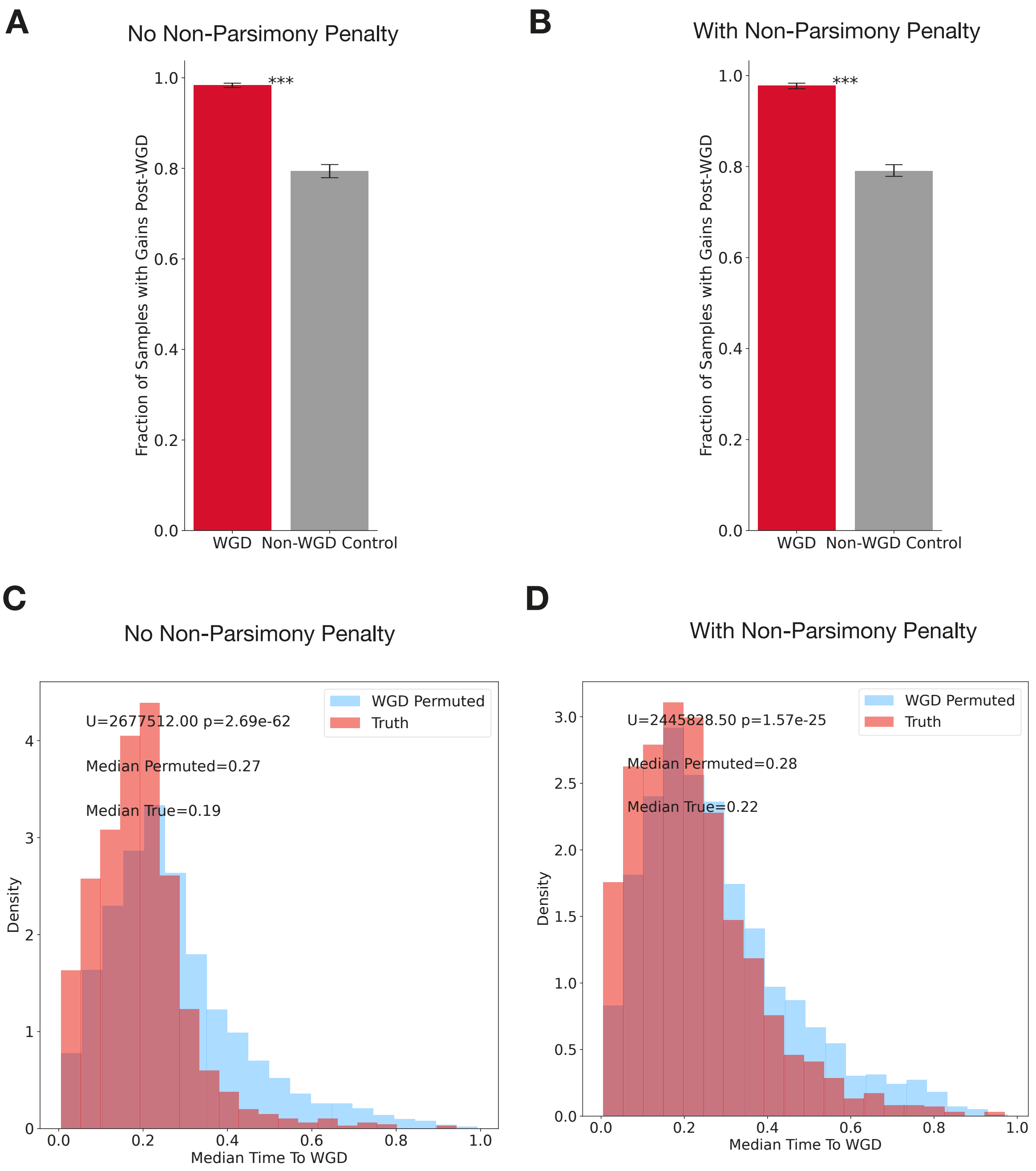


**Supplementary Figure 21. The timing of gains relative to WGD. A,B** Proportion of samples with gains post-WGD for WGD tumors and a cohort of control non-WGD tumors with a pseudo-WGD timing randomly sampled from WGD tumors with the same cancer type without (A) and with (B) a penalty on non-parsimony during inference. Statistical significance is calculated with a permutation test and 95% confidence intervals by bootstrapping over samples **C,D** Distribution of the median mutation time between the distribution of the timing of all independent gains and the median WGD timing for each sample for the correct WGD timing and a cohort where WGD timing is permuted between samples of the same cancer type. Applied without (C) and with (D) a penalty on non-parsimony during inference. Statistical significance was calculated by Mann Whitney U test.

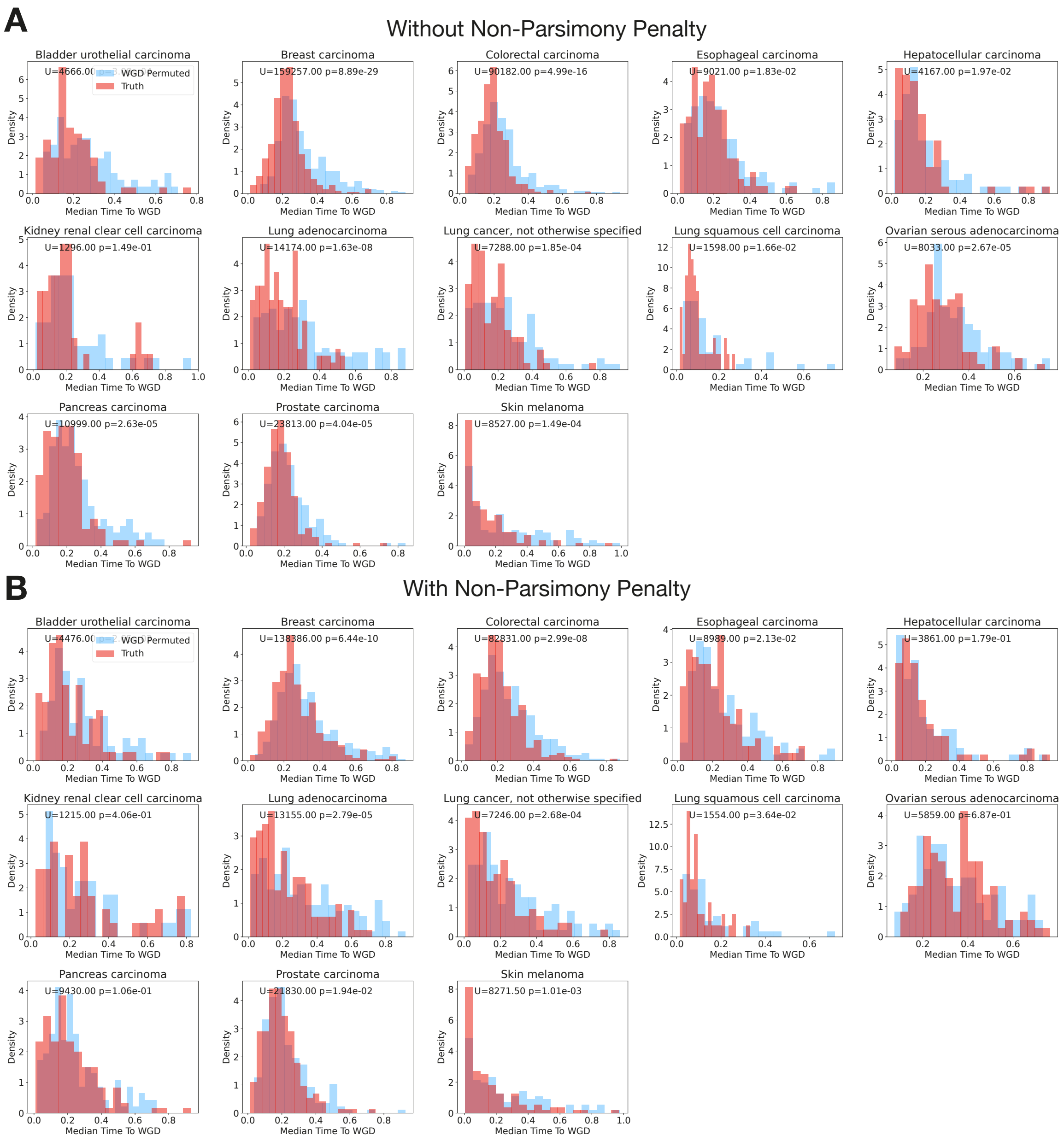


**Supplementary Figure 22. The timing of gains relative to WGD by cancer type. A,B** Distribution of the median mutation time between the distribution of the timing of all independent gains and the median WGD timing for each sample for the correct WGD timing and a cohort where WGD timing is permuted between samples of the same cancer type. Split by cancer type. Measured without (A) and with (B) a penalty on non-parsimony during inference. Statistical significance was calculated by Mann Whitney U test.

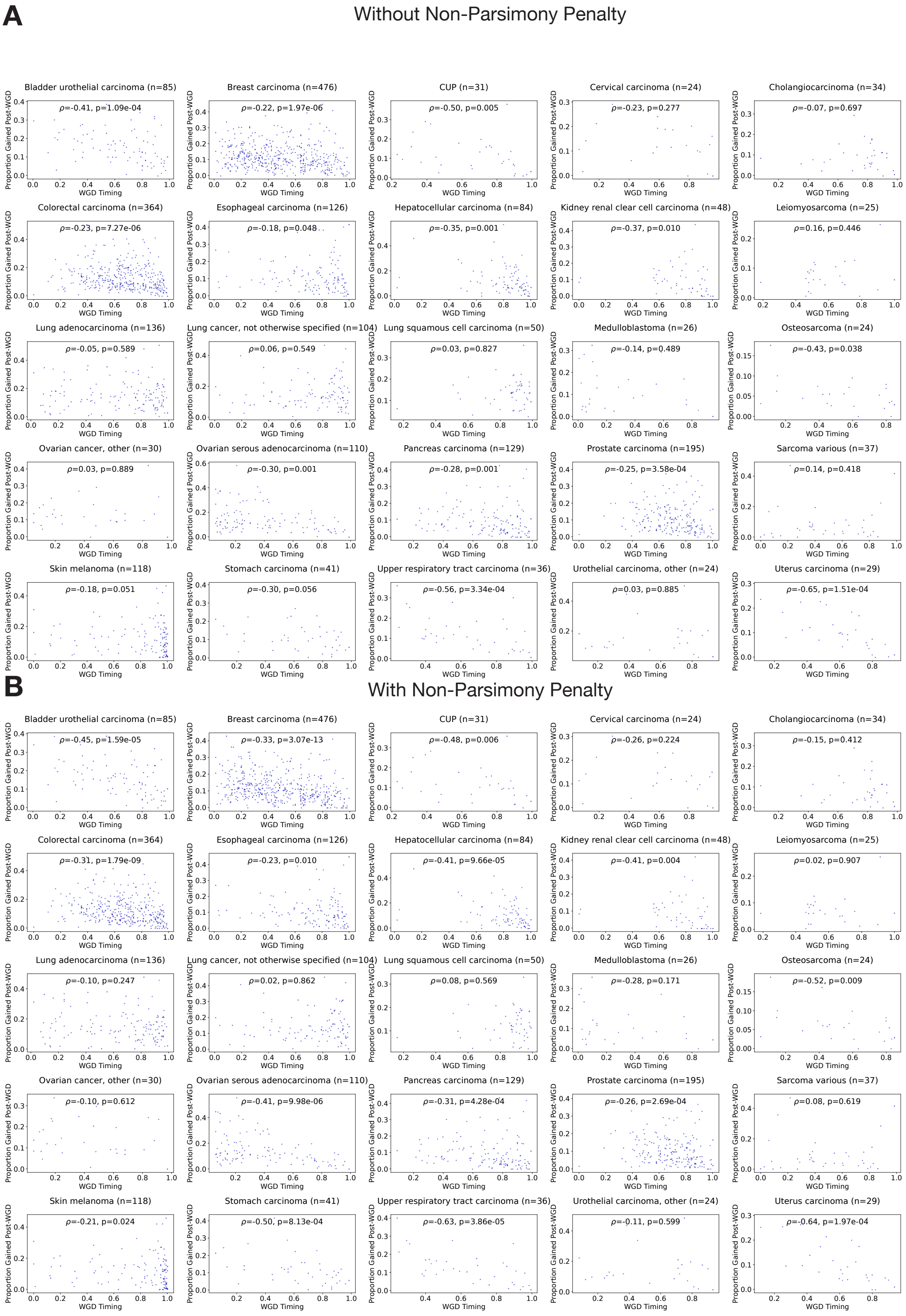


**Supplementary Figure 23. The relationship between genome gained post-WGD and WGD timing by cancer type.** Proportion of genome gained after the WGD against WGD timing for WGD tumors in PCAWG and Hartwig. Measured without (A) and with (B) a penalty on non-parsimony during inference. Correlation coefficient measured by Spearman's.

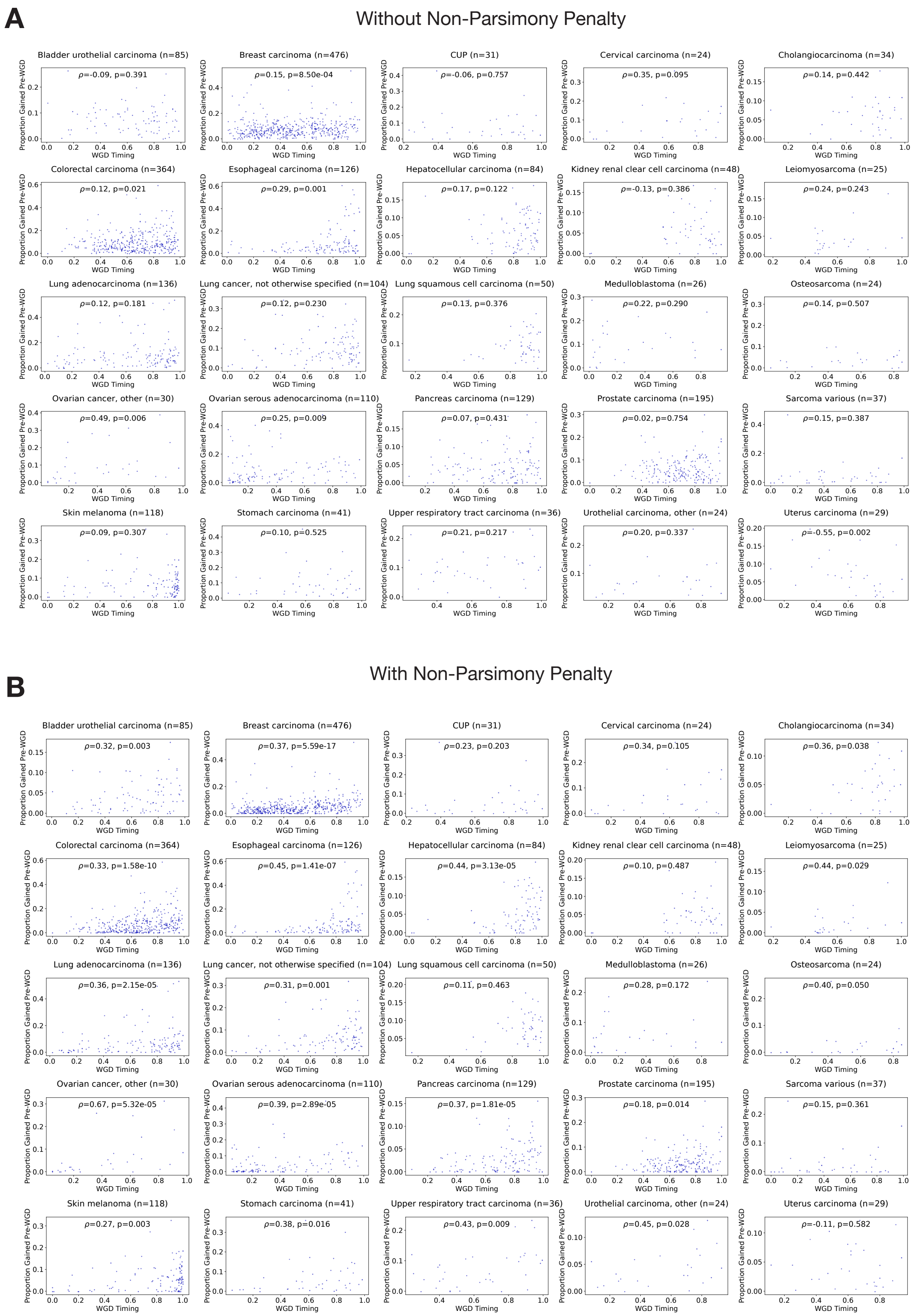


**Supplementary Figure 24. The relationship between genome gained pre-WGD and WGD timing by cancer type.** Proportion of genome gained before the WGD against WGD timing for WGD tumors in PCAWG and Hartwig. Measured without (A) and with (B) a penalty on non-parsimony during inference. Correlation coefficient measured by Spearman's.

**A**

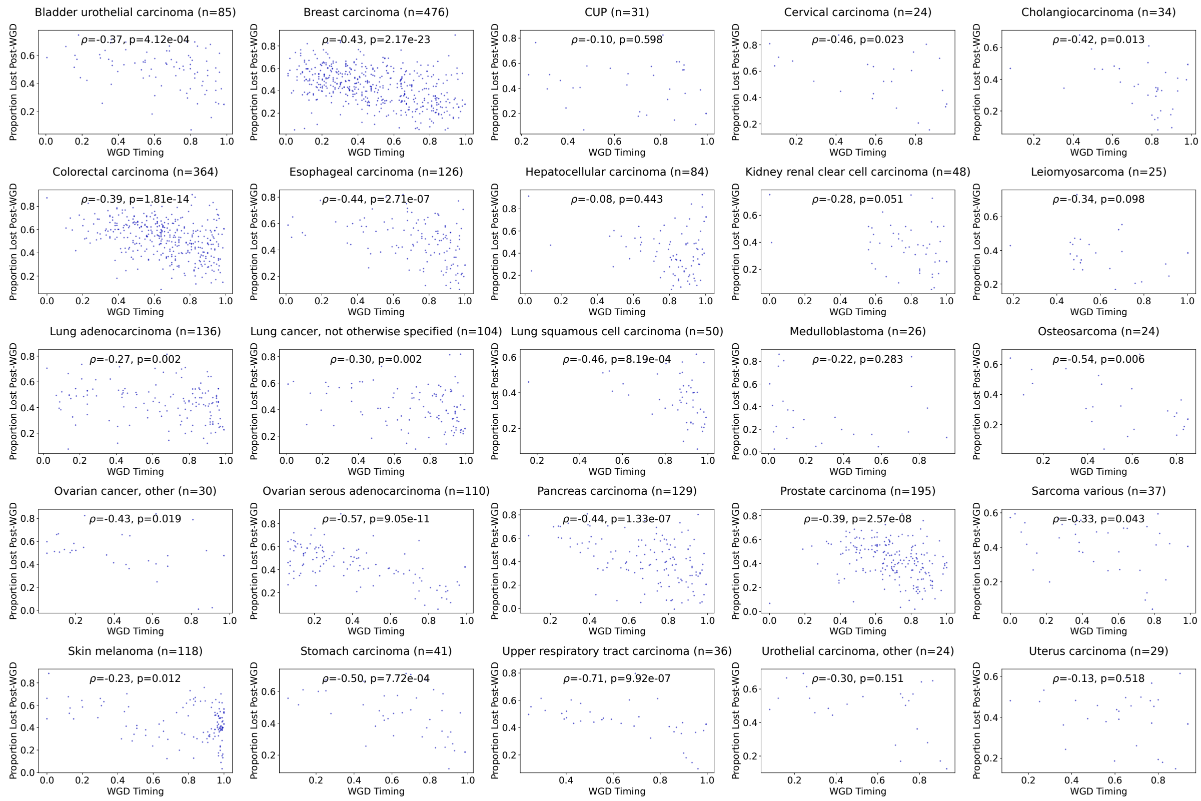


**B**

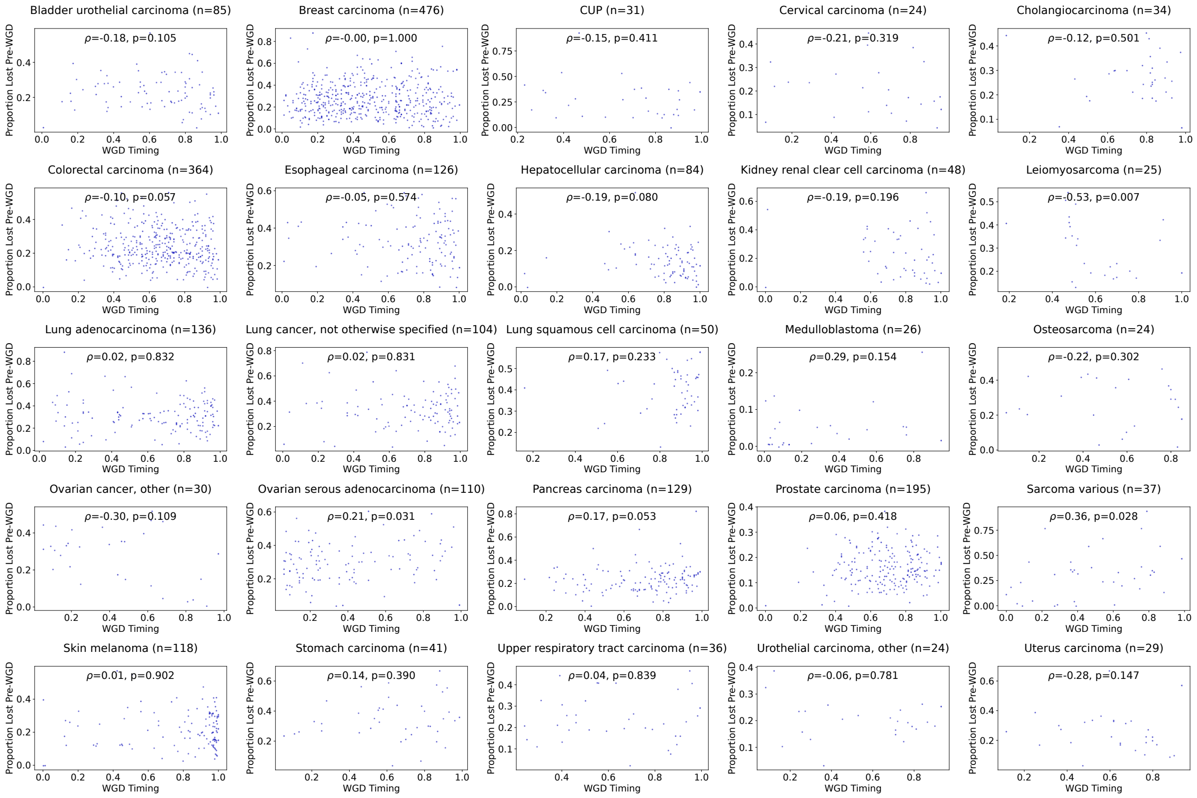


**Supplementary Figure 25. The relationship between fraction of genome lost pre and post-WGD and WGD timing by Cancer Type. A** Proportion of genome lost before and after the WGD against WGD timing for WGD tumors in PCAWG and Hartwig. Correlation coefficient measured by Spearman's.

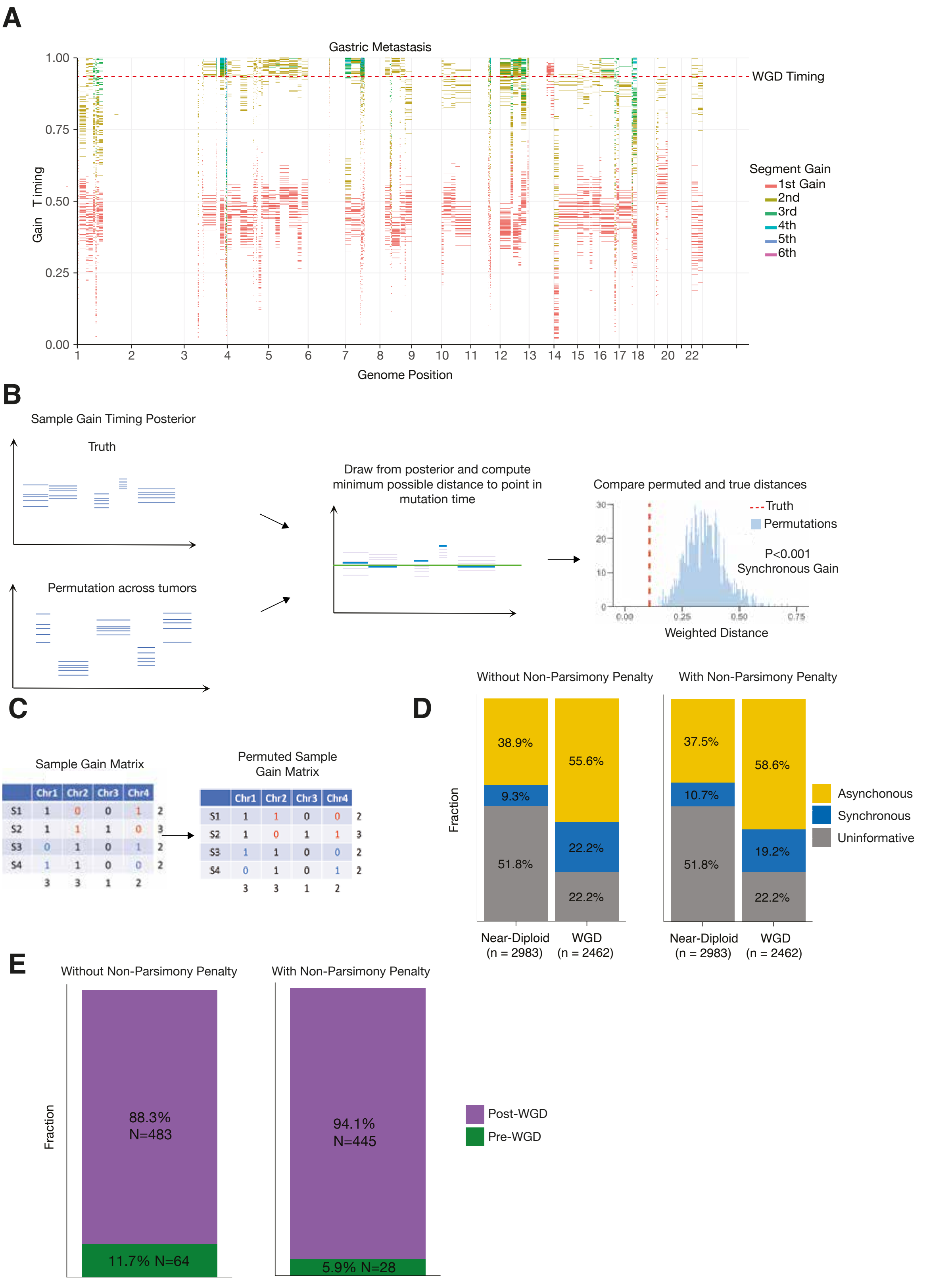


**Supplementary Figure 26. Punctuated gains in WGD tumors. A,** Example posterior distribution of copy number gain timing in a genome duplicated sample with a punctuated burst of gains. **B,** Schematic of procedure to identify samples with copy number gains occurring over a significantly shorter time period than expected under a permutation model. **C,** Schematic of a permutation scheme that maintains the number of gain chromosomes per sample and the number of times a chromosome is gained across the cohort. **D,** The proportion of tumors that have clonal gains identified as *occurring* synchronously or asynchronously, or uninformative where the number of gains was too low to classify, split by WGD status, with and without a penalty on non-parsimony. **e,** The proportion of synchronous gains occurring in WGD samples classified by whether they occurred pre- or post-WGD, with and without a penalty on non-parsimony.

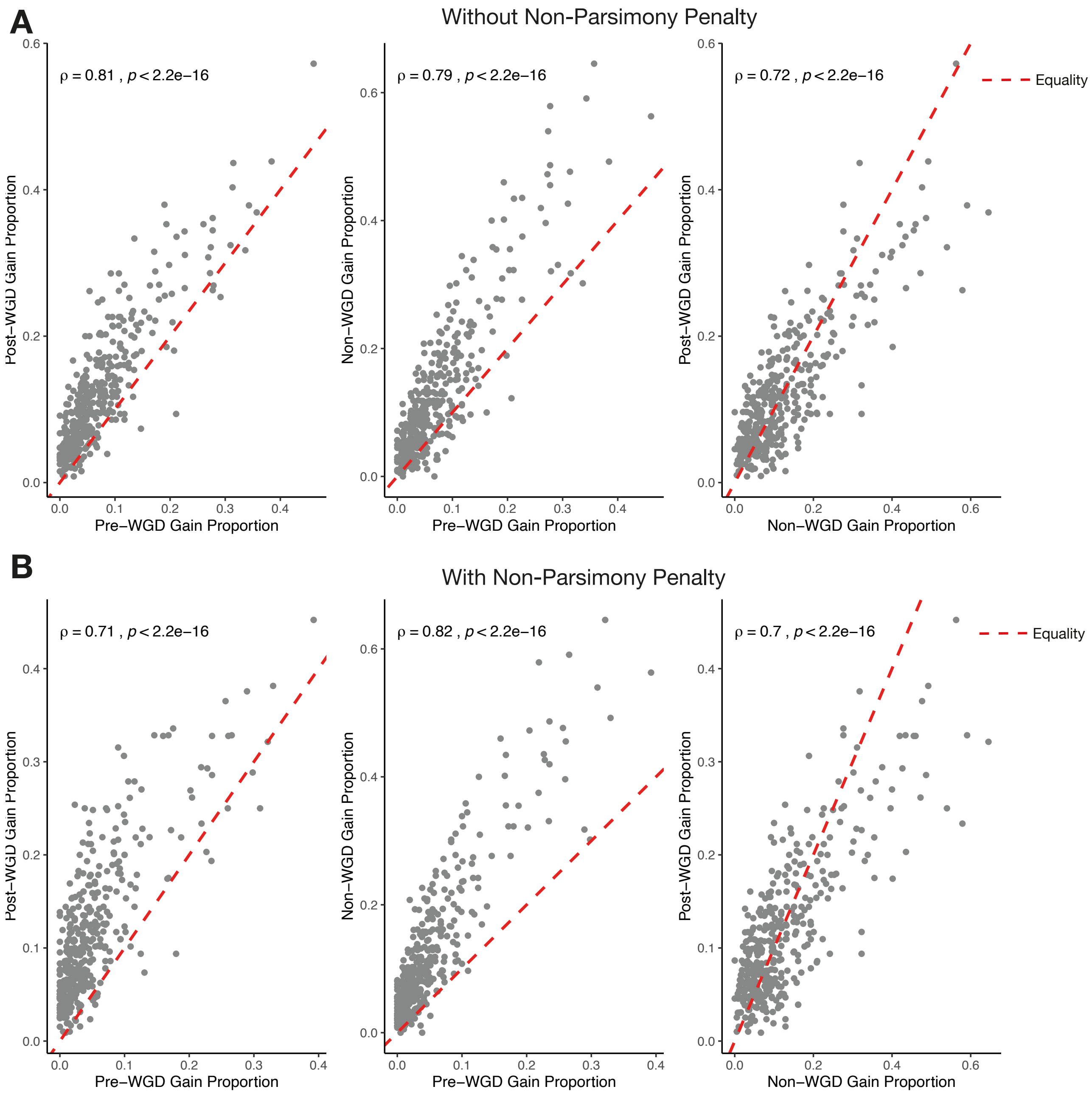


**Supplementary Figure 27. Frequency of arm gains pre and post-WGD and in non-WGD tumors.** The proportion of samples in different cancer types that have gained different chromosome arms pre- and post-WGD and in non-WGD samples. Measured without **(A)** and with **(B)** a penalty on non-parsimony during inference. Correlation is measured using Spearman's correlation coefficient.

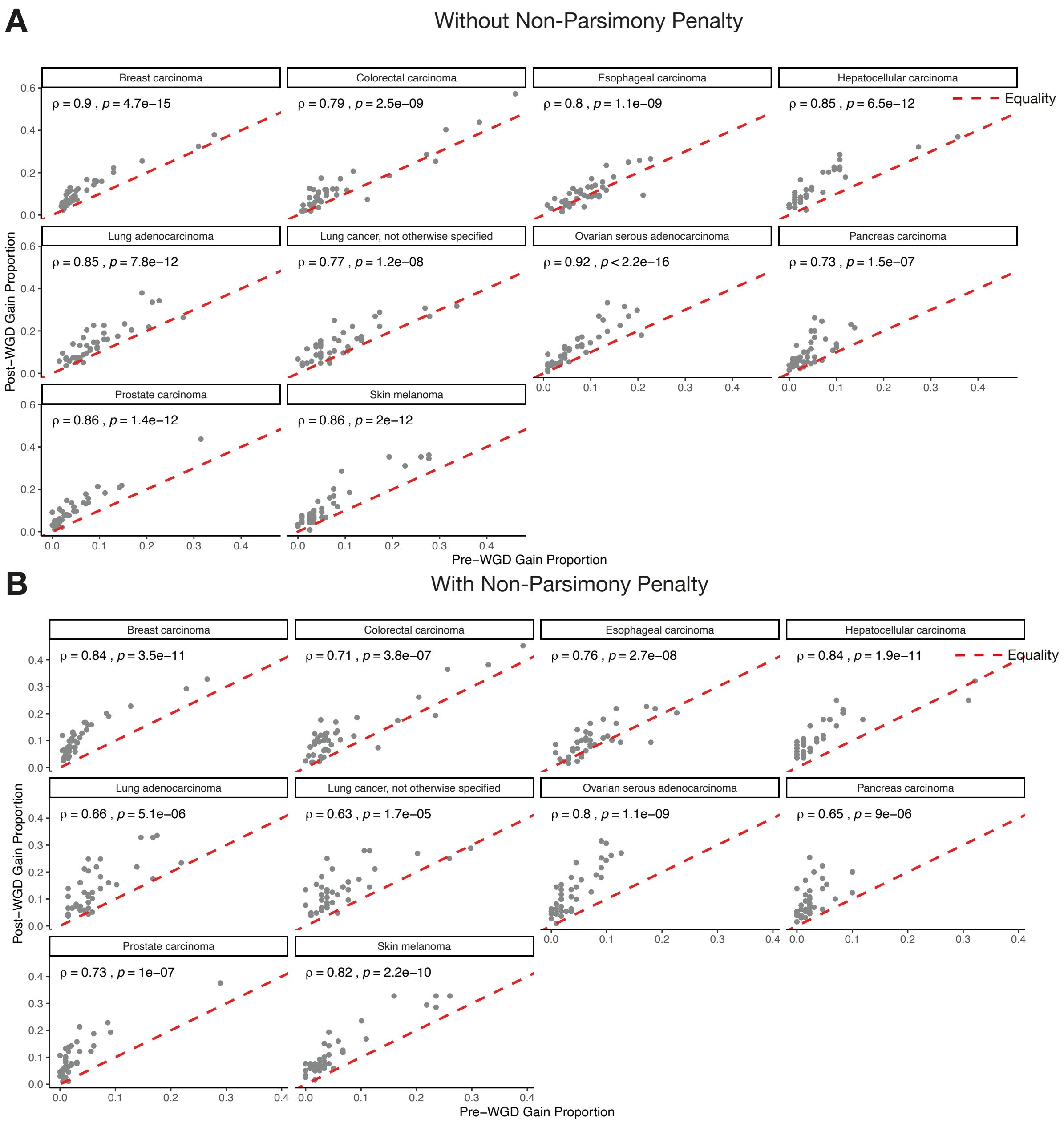


**Supplementary Figure 28. Frequency of arm gains pre and post-WGD and in non-WGD tumors by cancer type.** The proportion of samples that have gained different chromosome arms pre- and post-WGD split by cancer type. Measured without **(A)** and with **(B)** a penalty on non-parsimony during inference. Correlation is measured using Spearman's correlation coefficient.

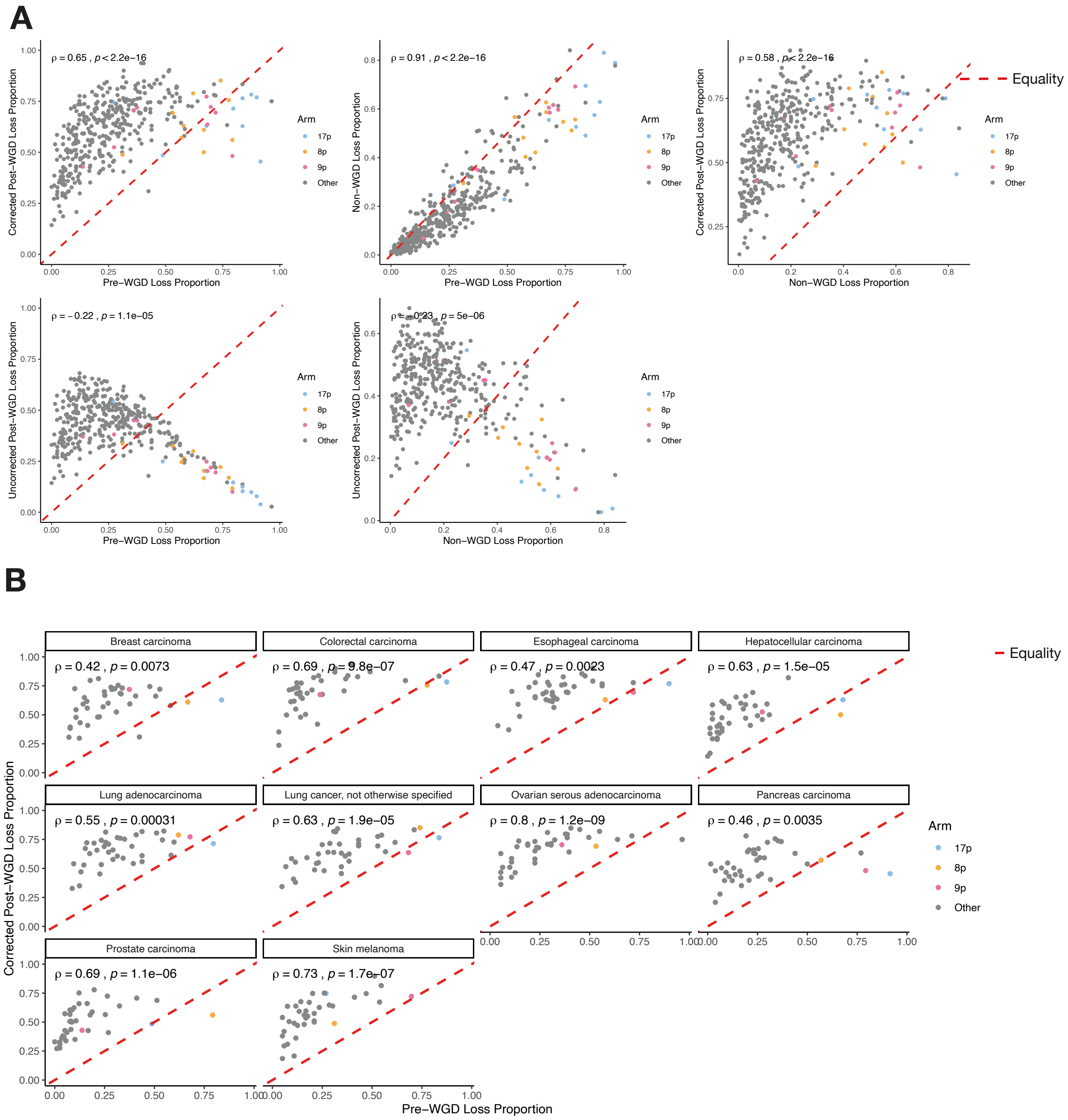


**Supplementary Figure 29. Frequency of arm losses pre and post-WGD and in non-WGD tumors by cancer type. A,** The proportion of samples in different cancer types that have lost different chromosome arms pre- and post-WGD and in non-WGD samples. **B,** The proportion of samples that have lost different chromosome arms pre- and post-WGD split by cancer type. Correlation is measured using Spearman's correlation coefficient.

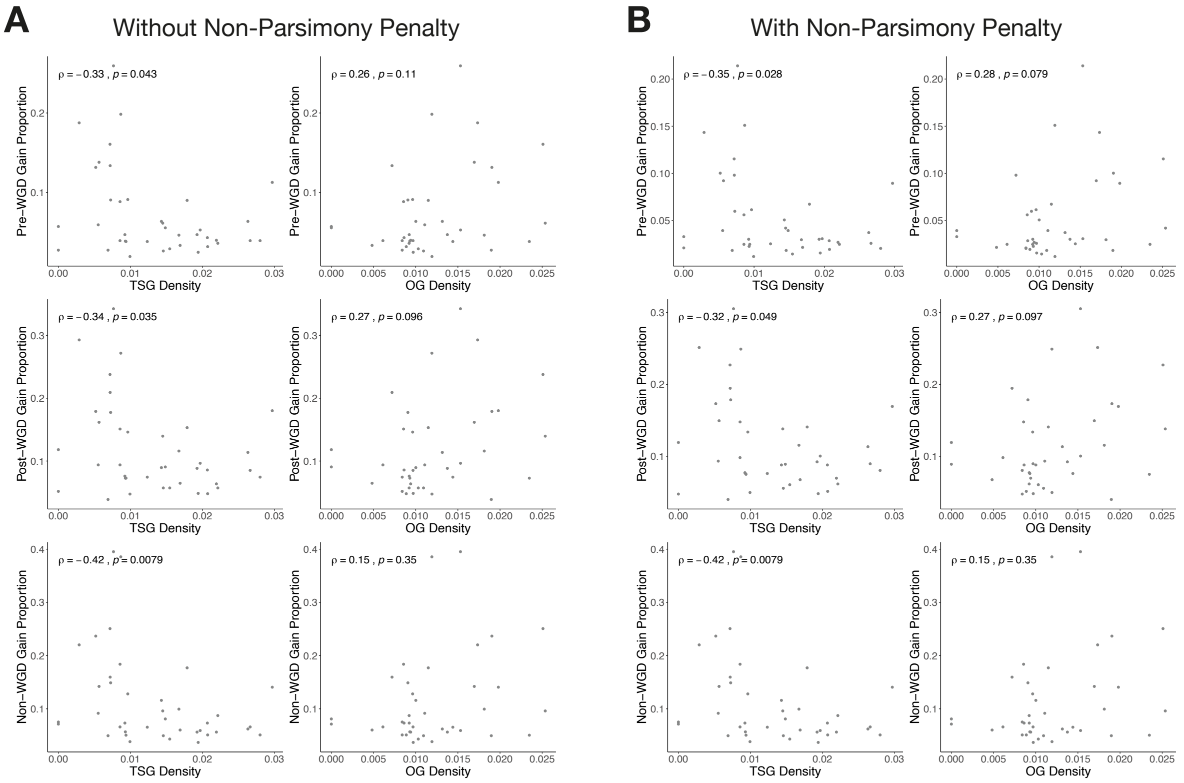


**Supplementary Figure 30. Effect of oncogene and tumor suppressor gene density on arm gain rates.** The proportion of samples combined across cancer types that have gained different chromosome arms pre- and post-WGD and in non-WGD samples against arm oncogene (OG) and tumor suppressor gene (TSG) density. Measured without **(A)** and with **(B)** a penalty on non-parsimony during inference. Correlation is measured using Spearman's correlation coefficient.

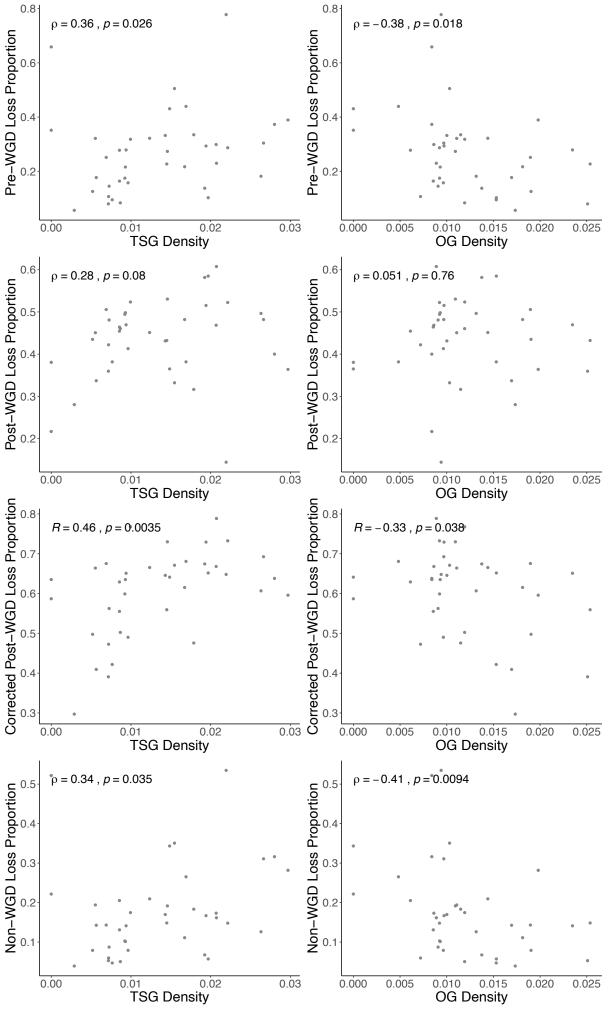


**Supplementary Figure 31. Effect of oncogene and tumor suppressor gene density on arm loss rates.** The proportion of samples combined across cancer types that have lost different chromosome arms pre- and post-WGD and in non-WGD samples against arm oncogene (OG) and tumor suppressor gene (TSG) density. Correlation is measured using Spearman's correlation coefficient.

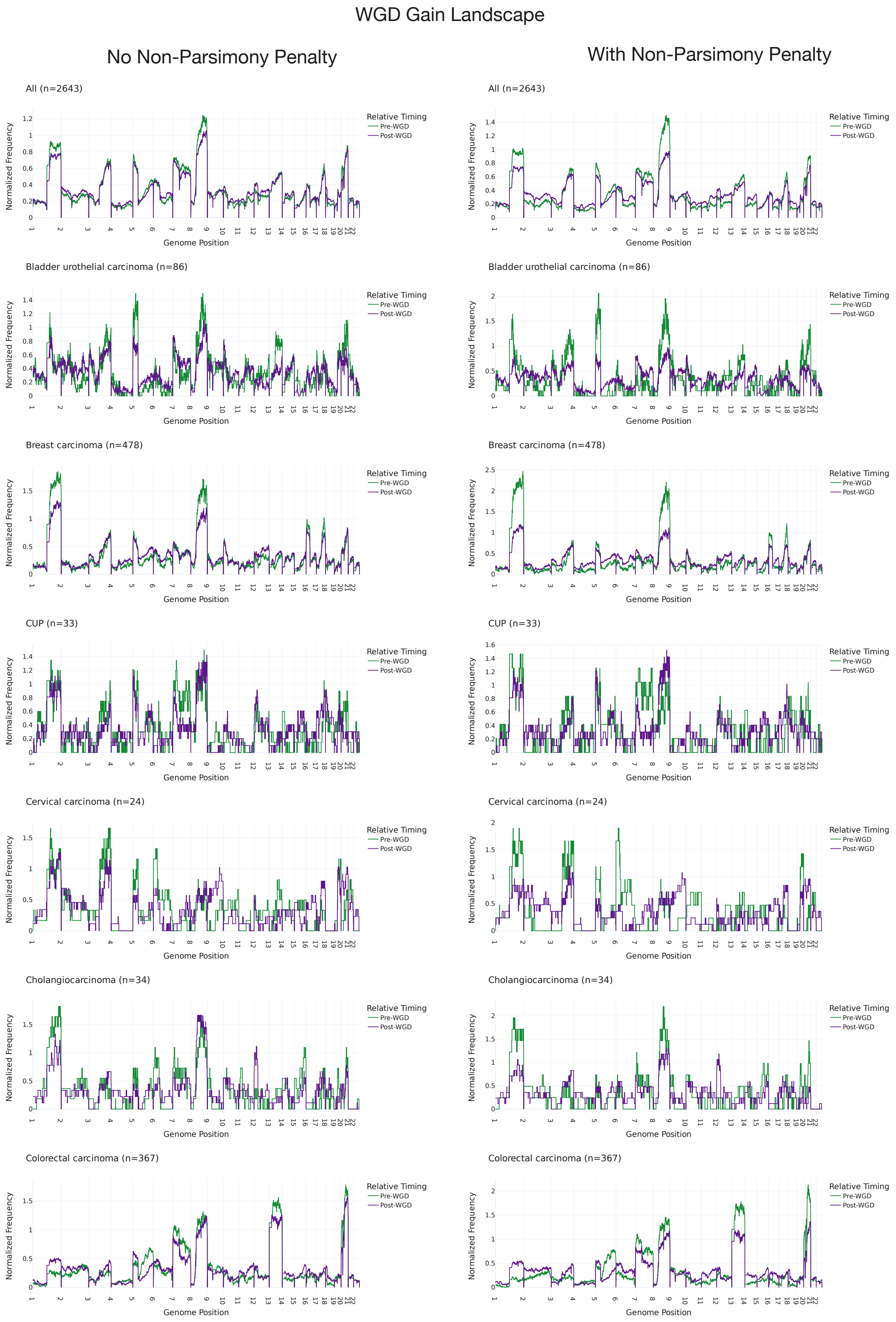


**Supplementary Figure 32. Pan genome frequencies of pre and post-WGD gains by cancer type.** Frequency of pre- and post-WGD gains for different cancer types. The frequencies are normalized so that the pre- and post-WGD frequency integrate to the same constant. Measured with and without a penalty on non-parsimonious routes during inference.

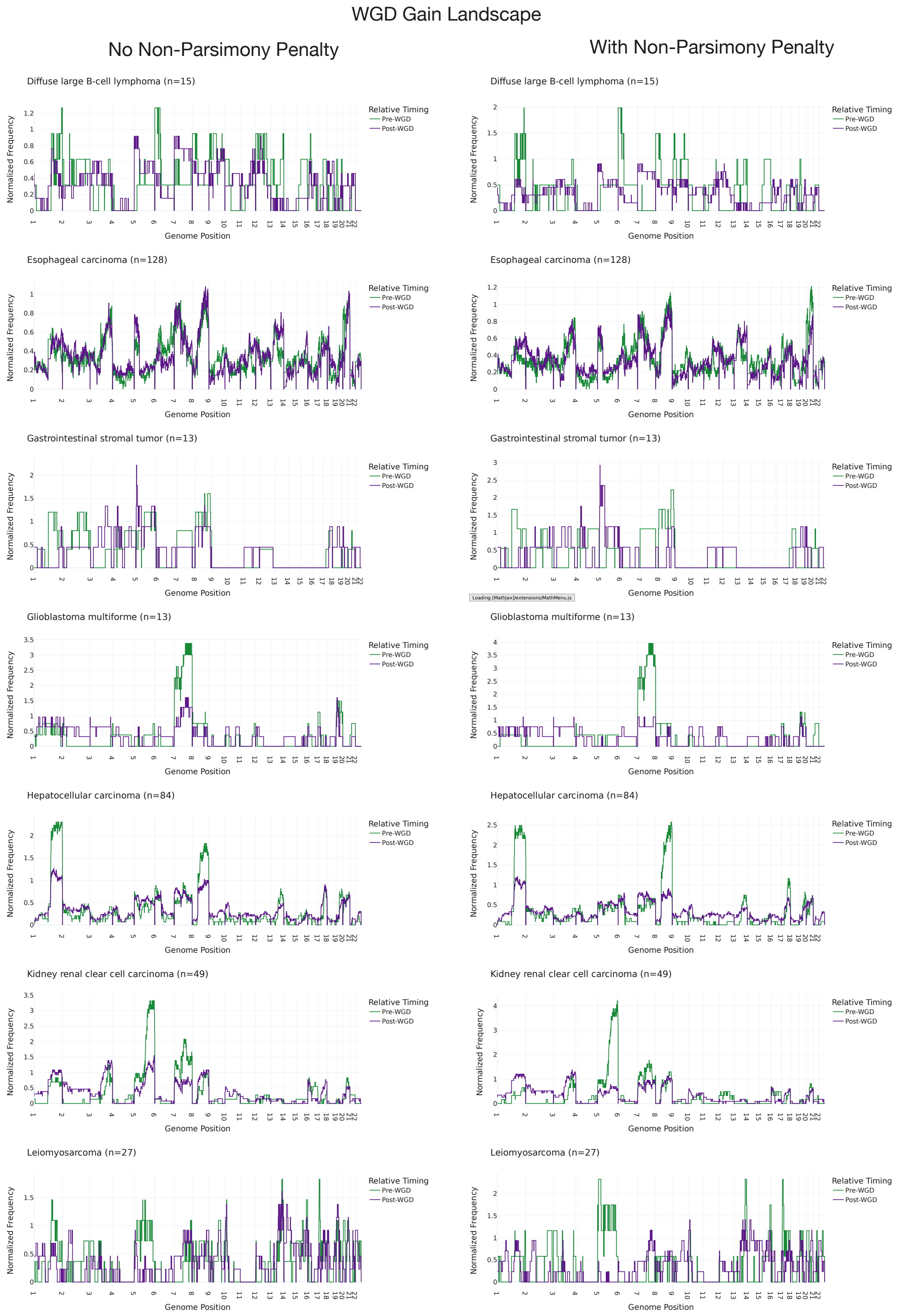


**Supplementary Figure 33. Pan genome frequencies of pre and post-WGD gains by cancer type.** Frequency of pre- and post-WGD gains for different cancer types. The frequencies are normalized so that the pre- and post-WGD frequency integrate to the same constant. Measured with (right column) and without (left column) a penalty on non-parsimonious routes during inference.

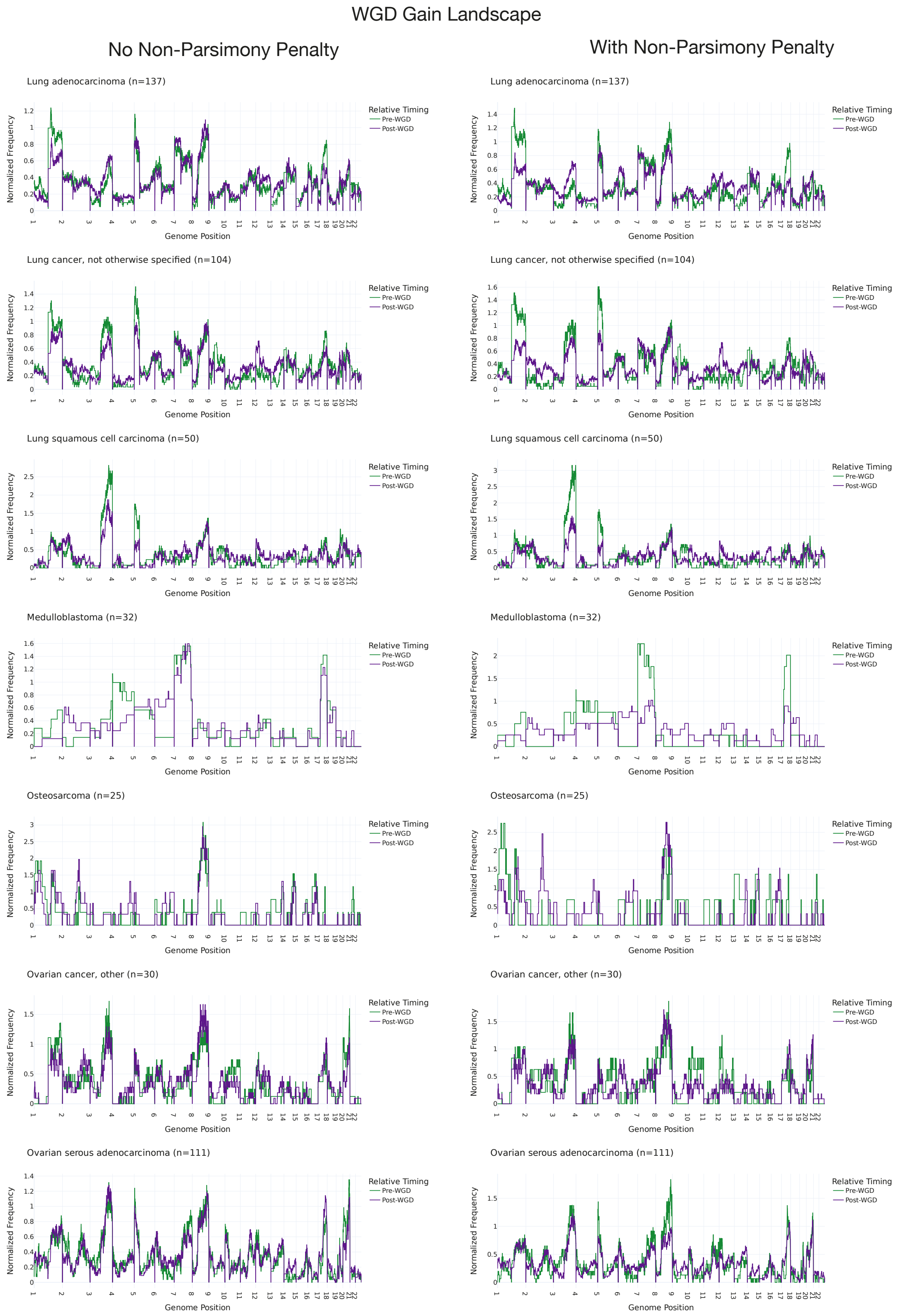


**Supplementary Figure 34. Pan genome frequencies of pre and post-WGD gains by cancer type.** Frequency of pre- and post-WGD gains for different cancer types. The frequencies are normalized so that the pre- and post-WGD frequency integrate to the same constant. Measured with (right column) and without (left column) a penalty on non-parsimonious routes during inference.

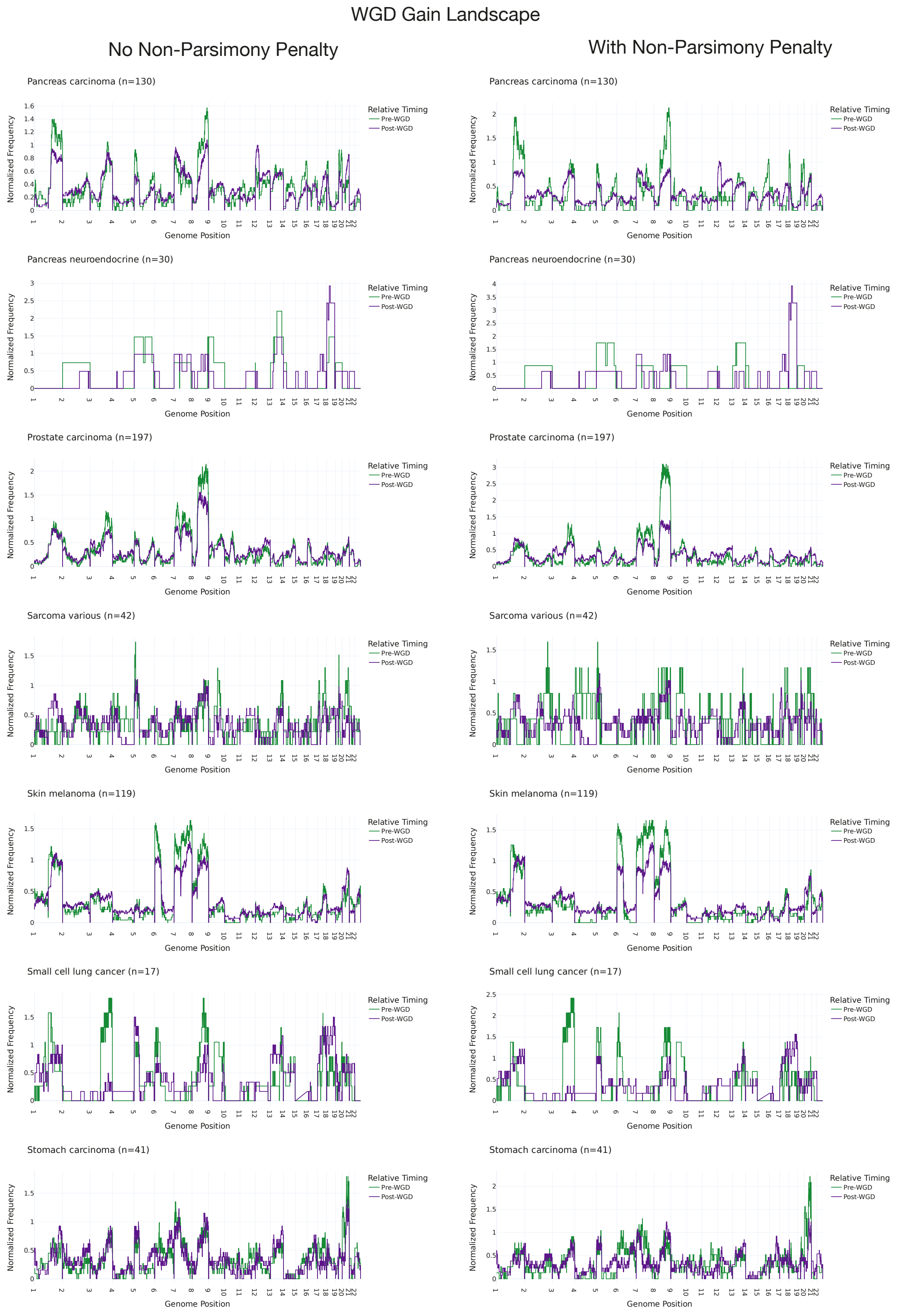


**Supplementary Figure 35. Pan genome frequencies of pre and post-WGD gains by cancer type.** Frequency of pre- and post-WGD gains for different cancer types. The frequencies are normalized so that the pre- and post-WGD frequency integrate to the same constant. Measured with (right column) and without (left column) a penalty on non-parsimonious routes during inference.

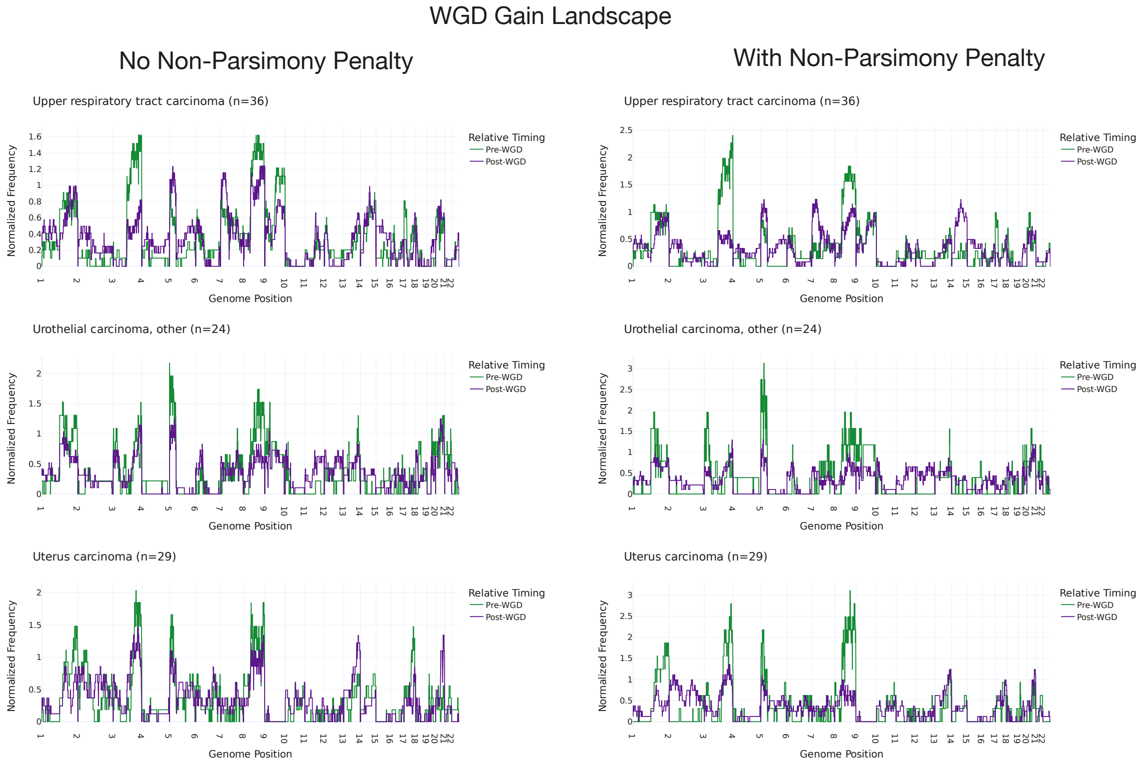


**Supplementary Figure 36. Pan genome frequencies of pre and post-WGD gains by cancer type.** Frequency of pre- and post-WGD gains for different cancer types. The frequencies are normalized so that the pre- and post-WGD frequency integrate to the same constant. Measured with (right column) and without (left column) a penalty on non-parsimonious routes during inference.

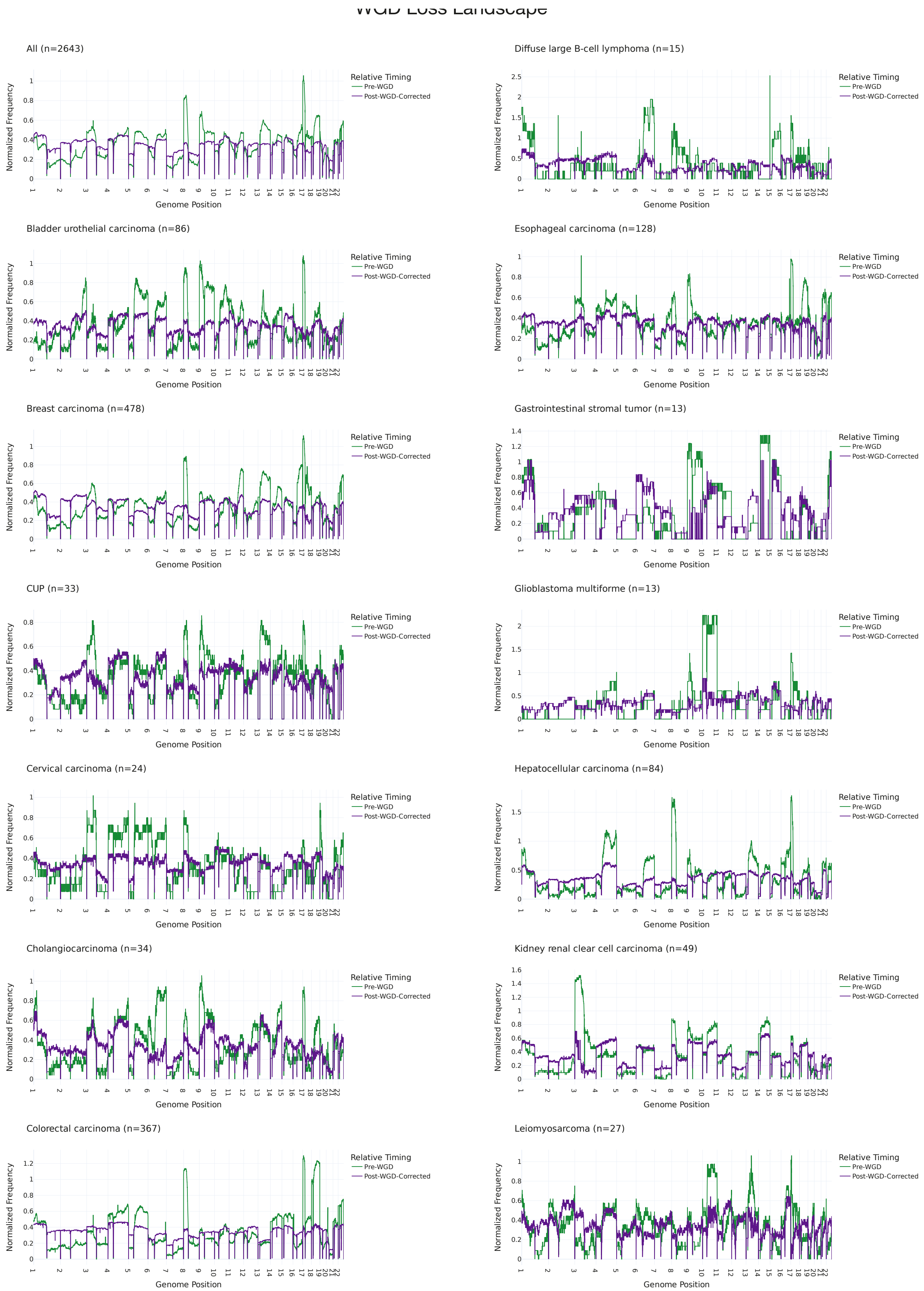


**Supplementary Figure 37. Pan genome frequencies of pre and post-WGD losses by cancer type.** Frequency of pre- and post-WGD losses for different cancer types. The frequencies are normalized so that the pre- and post-WGD frequency integrate to the same constant. The post-WGD loss frequency is corrected to account for mutual exclusivity when measuring pre- and post-WGD losses.

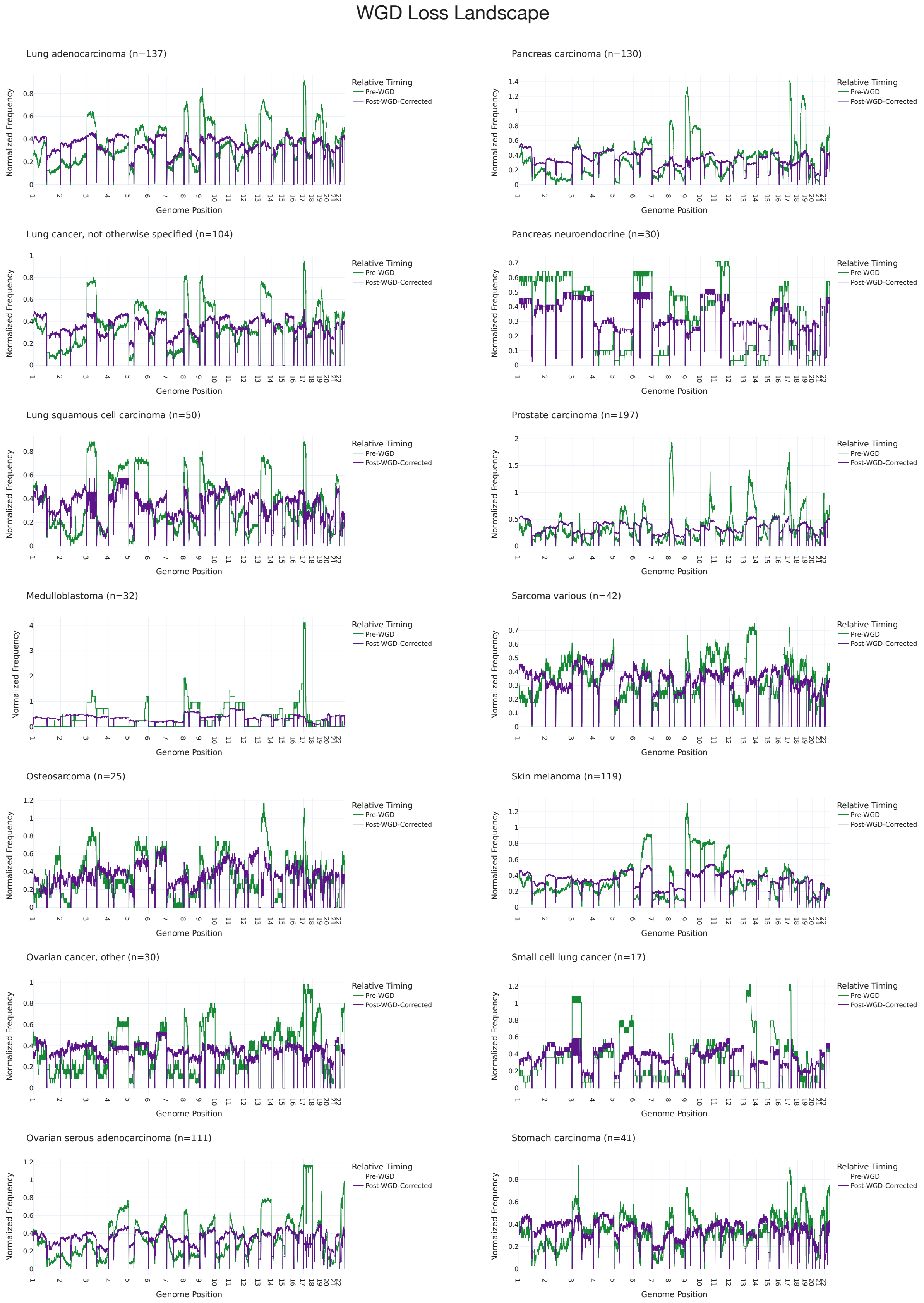


**Supplementary Figure 38. Pan genome frequencies of pre and post-WGD losses by cancer type.** Frequency of pre- and post-WGD losses for different cancer types. The frequencies are normalized so that the pre- and post-WGD frequency integrate to the same constant. The post-WGD loss frequency is corrected to account for mutual exclusivity when measuring pre- and post-WGD losses.

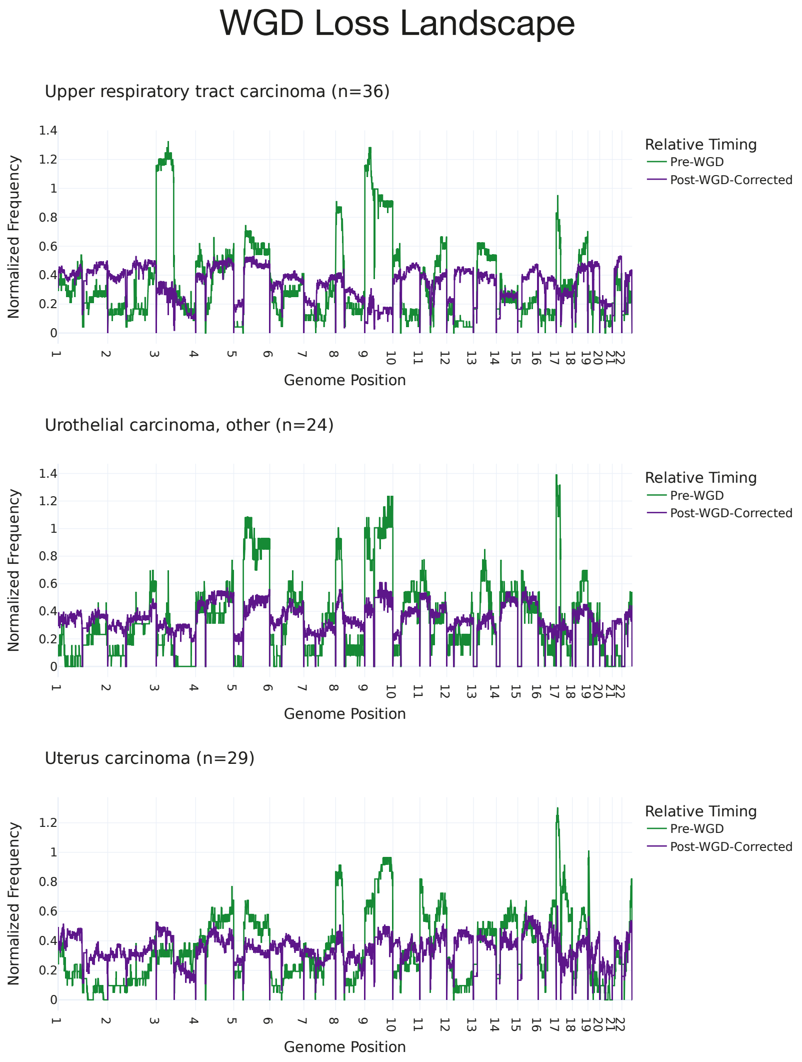


**Supplementary Figure 39. Pan genome frequencies of pre and post-WGD losses by cancer type.** Frequency of pre- and post-WGD losses for different cancer types. The frequencies are normalized so that the pre- and post-WGD frequency integrate to the same constant. The post-WGD loss frequency is corrected to account for mutual exclusivity when measuring pre- and post-WGD losses.

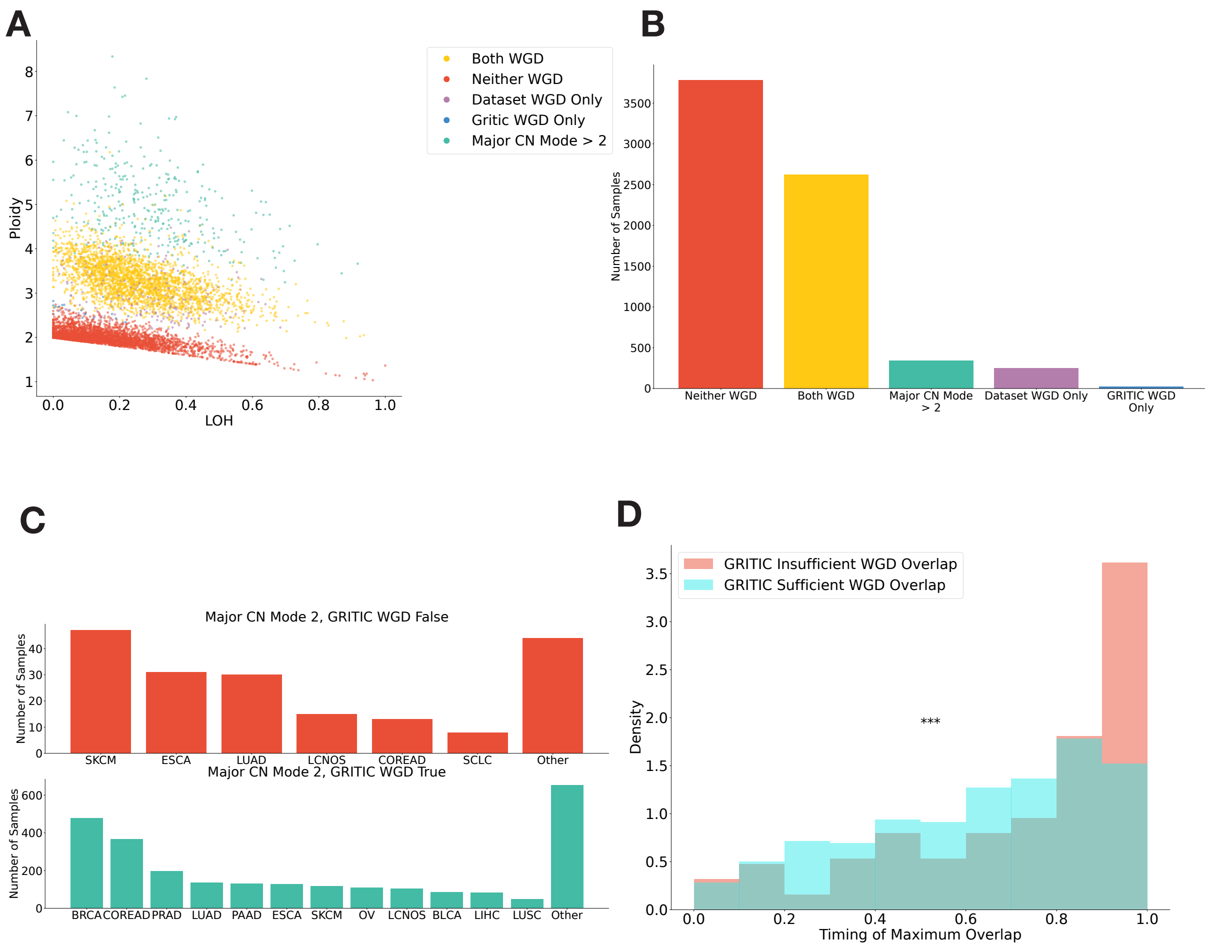


**Supplementary Figure 40. WGD status calling in GRITIC. A,** Proportion of genome with loss of heterozygosity and tumor ploidy for samples in the PCAWG and Hartwig cohorts. Samples are colored by whether they were called as WGD by one or both of GRITIC and from their copy number profiles only. Samples with major copy number mode greater than of two are categorised separately as they are not supported by GRITIC. **B,** Counts of samples with different WGD calling statuses across Hartwig and PCAWG cohorts. **C,** Counts of tumor with major copy number two in PCAWG and Hartwig by cancer type. Cohort split by WGD calling status in GRITIC. **D,** The timing that maximally intersected the timing confidence intervals of major copy number two states. The distributions are split by whether such timing intersected with 60% of major copy number two segments by width. Only tumors with a major copy number mode of two are displayed. Statistical significance was calculated by Mann Whitney U Test.
