## Supplementary Methods for "The history of chromosomal instability in genome doubled tumors"

#### Contents

**1 Summary** **3**

**2 Principles of mutation time** **3**

**3 Mutation time in practice** **5**

**4 Limitations and assumptions of mutation time** **9**

**5 Problems with timing more complex gains** **11**

**6 Representing routes as binary trees** **13**

**7 Using the representations to time complex gains** **16**

**8 Multiplicity spaces** **19**

**11 Nearest neighbor evaluation methods 37**

### 1 Summary

In these supplementary methods, we describe the theoretical basis for Gain Route Identification and Timing In Cancer (GRITIC), our novel method for timing complex copy number gains from single nucleotide variant (SNV) data. We first derive the quantitative relationship between SNV multiplicity proportions and the timing of simple copy number gains and discuss a number of assumptions and limitations that underpin mutation time.

Then, we introduce a tree-based representation of copy number event histories and use this to infer the relationship between SNV multiplicity proportions and gain timings for complex copy number states. We next describe how these principles are applied in GRITIC, a new generalized method for timing copy number gains. Finally, we describe the technical details of the GRITIC implementation.

### 2 Principles of mutation time

Using a limited number of assumptions, the timing of clonal copy number gains can be measured relative to the tumor SNV burden. When a copy number gain occurs, SNVs in the region of the gain are duplicated onto the newly gained allele and are therefore present on two copies in the cell. The number of allelic copies that an SNV is present on is known as its multiplicity.

Absent a subsequent copy number loss, SNVs that occur before the gain will have multiplicities greater than one. We can then apply the infinite sites assumption [1], which posits that the genome can be treated as infinite in size and, therefore, any base pair can be mutated at most once in the development of a tumor. Therefore, the only route for a mutation to have multiplicity greater than one is through duplication in a copy number gain.

Using the infinite sites assumption in segments with a copy number gain, all SNVs with multiplicity two on the gained parental allele must have occurred before the gain and all those with multiplicity one must have occurred after

(Fig. 1a). This can be used to infer the timing of the gain relative to the accumulation of SNVs. The greater the number of SNVs with multiplicity two, the later the copy number gain occurred (Fig. 1b). For a given copy number segment, the larger number of parental alleles is defined as the major copy number and the smaller the minor copy number. Under the infinite sites assumption, the multiplicity of an SNV can be no larger than the major copy number of the section of the genome where it is located.

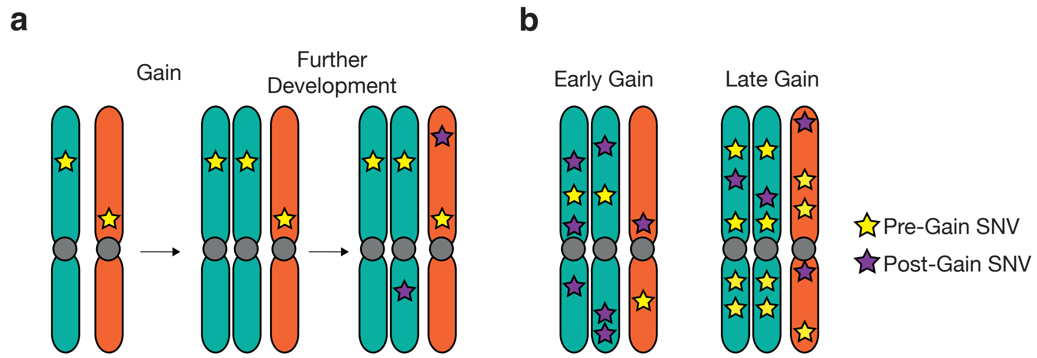

Figure 1: **a.** Schematic of the accumulation of SNVs during tumor development. SNVs on the gained allele are copied over to the new allele. **b.** Illustration of the concept that the earlier a copy number gain occurs, the greater the ratio of SNVs on single copies to multiple copies.

Given the above, gain timing can be assessed quantitatively by calculating the ratio of SNVs with multiplicity two to multiplicity one, after correcting for total genome content and the presence of SNVs on the non-gained allele. This gives the timing of the gain measured in mutation time. If only clonal SNVs are considered, mutation time has useful properties. It can be defined on a scale from zero to one, where zero is conception and one is the emergence of the progenitor cell to all cells in the tumor biopsy, the most recent common ancestor (MRCA).

Mutation time is defined to be linear with respect to SNV accumulation. As the mutation rate of a tumor typically increases over development, mutation time is non-linear with respect to real time. Equal intervals of mutation time likely correspond to shorter real time periods as the tumor develops (Fig. 2a). Correspondingly, equal intervals in real time correspond to larger periods of mutation time over tumor development (Fig. 2b).

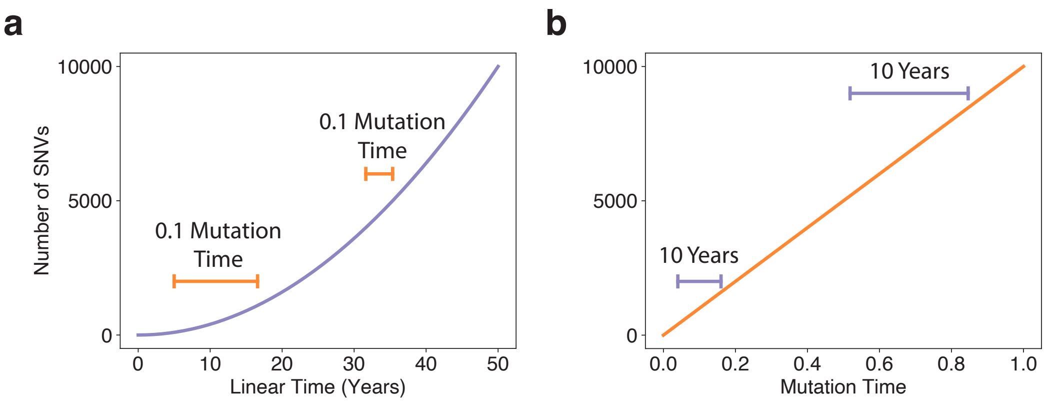

Figure 2: **a.** Schematic of a mutation accumulation in a tumor with a linearly increasing mutation rate. Equal units of mutation time are shown at different points in linear time. **b.** Mutation accumulation in the same tumor in mutation time. Equal units of real time are shown starting at different points in mutation time.

Although non-linear with respect to real time, mutation time is useful because it preserves order. If one gain occurs earlier than another in mutation time, it occurs earlier during tumor development. Mutation time can therefore be applied to build timelines for copy number gains in individual tumors, identifying which gains are the earliest to arise and which only occur when the tumor has already accumulated a number of SNVs and copy number gains.

### 3 Mutation time in practice

To illustrate how to use SNV multiplicities to time a simple gain, I consider the case of a genomic region with a single copy number gain (Fig. 3). The following derivation follows from the principles first outlined previous SNV-based gain timing methods [2–5].

Any mutations that occur on the gained parental allele in mutation time period $T_1$ will result in two copies. Those that occur on the non-gained allele at any time and on the gained allele during $T_2$ will only be present on one

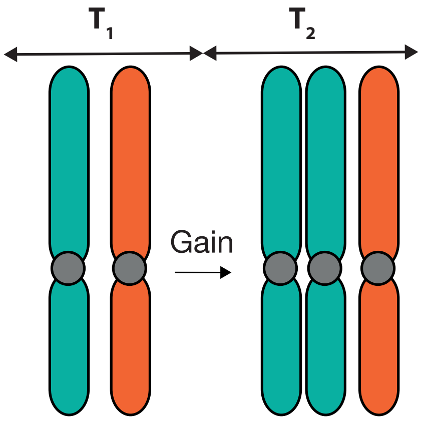

Figure 3: Schematic of a single copy number gain leading to a 2+1 state. The time period $T_1$ corresponds to the period between conception and the time of the gain and the period $T_2$ between the gain and the emergence of the most recent common ancestor of the tumor.

copy. We therefore obtain a set of equations for the number of mutations on $i$ copies $N_i$.

$$\begin{aligned} N_1 &= R(T_1 + 3T_2) \\ N_2 &= RT_1 \end{aligned} \tag{1}$$

Here $R$ is the mutation rate for the gained segment. $R$ is a constant as it is defined relative to mutation time. By the definition of mutation time, we also have that $T_1 + T_2 = 1$, and by reformulating the equations to use the proportion of SNVs on $i$ copies $m_i = \frac{N_i}{\sum_j N_j}$, we can solve for the time of the gain $T_1$.

$$T_1 = \frac{3m_2}{m_1 + 2m_2} \tag{2}$$

Note that the gain timing $T_1$ is independent of the mutation rate $R$. This is a very useful property of mutation time as it allows for gain timing to be compared within a tumor without directly considering the local mutation rate.

We now examine a second case where the non-gained allele is lost during development (Fig. 4). After considering the number of SNVs with different multiplicities at the final state, we arise at a similar set of equations to (1).

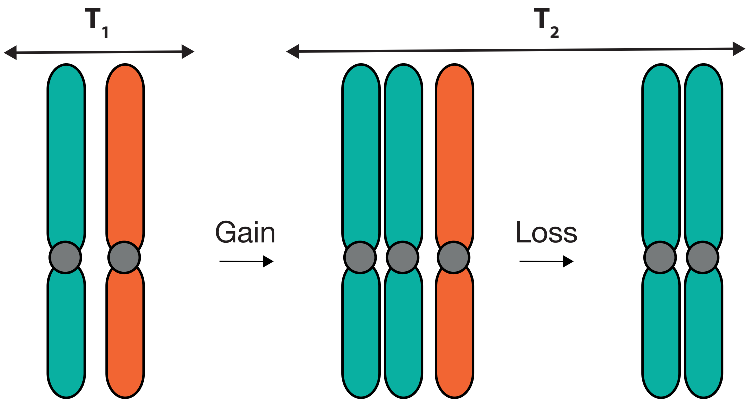

Figure 4: Schematic of a single copy number gain and loss leading to a 2+0 state. The time period $T_1$ corresponds to the period between conception and the time of the gain and the period $T_2$ between the gain and the emergence of the most recent common ancestor of the tumor. The loss occurs at an arbitrary time across the clonal period.

$$\begin{aligned} N_1 &= 2RT_2 \\ N_2 &= RT_1 \end{aligned} \tag{3}$$

The SNVs present on the allele that was lost cannot be observed, but this is not necessary to time the gain. The solved equation (4) is identical to the 2+1 case (2) up to a decrease in the multiplicative constant. The reduction in the multiplicative constant corrects for the lack of multiplicity one SNVs that would have occurred from the lost allele. This equation applies regardless of when the loss occurs during the clonal evolutionary period.

$$T_1 = \frac{2m_2}{m_1 + 2m_2} \tag{4}$$

Finally, consider the case of gains leading to a 2+2 copy number state. Similar to before, we have the following multiplicity equations:

$$
\begin{aligned}
N_1 &= R(2T_2 + 4T_3) \\
N_2 &= R(2T_1 + T_2)
\end{aligned} \tag{5}
$$

There are now four variables and three equations (including $\sum_i T_i = 1$) and thus this system of equations is under-determined. It can however, be solved for $T_1$ and $T_2$ together.

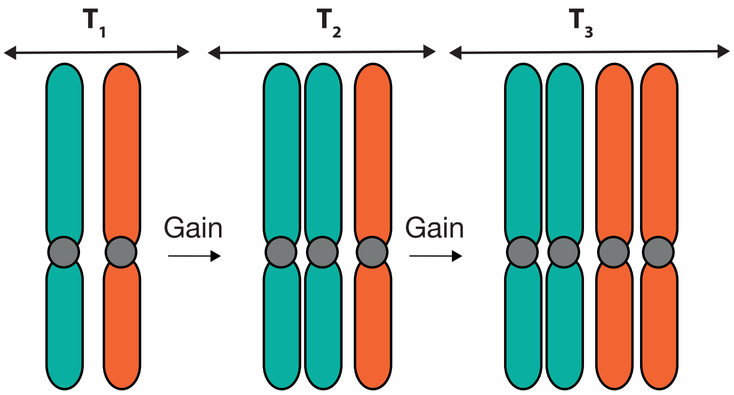

Figure 5: Schematic of the copy number gains leading to a 2+2 state. The time period $T_1$ corresponds to the period between conception and the time of the first gain, $T_2$ to the period between the first and second gain and the period $T_3$ to the period between the second gain and the emergence of the most recent common ancestor of the tumor.

$$
T_1 + \frac{T_2}{2} = \frac{2m_2}{m_1 + 2m_2} \tag{6}
$$

Further progress can be made if we assume that the two copy number gains are simultaneous, such as in a whole genome duplication (WGD). Then $T_2 = 0$ and $T_1$ is fully specified. This was the innovation first applied by Gerstung *et al.* to time gains leading to 2+2 states [6]. Note that under this condition, equation 6 is identical to the timing equation for a 2+0 gain, equation 4. This follows as both parental alleles in the 2+2 state can each be considered as a separate 2+0 state with identical gain timing.

### 4 Limitations and assumptions of mutation time

The principles of timing copy number gains outlined in the previous section rely on a number of assumptions and have limitations in order to obtain accurate timing.

#### 4.1 Timing in the clonal period

Mutation time is typically only extended to the clonal evolutionary period. This is because all observed SNVs that occurred during this period were all present in a single cell, the tumor's MRCA. This means that they accumulated sequentially and can therefore be used as a reference timescale.

In the subclonal period of the tumor, this is no longer the case. Different mutations can accumulate in parallel across different subclones, or sequentially in descendant subclones. While parallel SNV accumulation can be modelled in mutation timing frameworks [6], it relies on accurate subclonal phylogenies and SNV assignments to each subclone to obtain the correct accumulation models. As subclonal phylogenies obtained from single sample biopsies are often ambiguous [7], timing from such data is typically restricted to the clonal evolutionary period.

#### 4.2 Mutation rates

The SNV multiplicity ratios are normalized for the overall mutation burden of each segment. This means that a uniform mutation rate across the genome is not necessary to compare the timing of gains across different segments within a single tumor.

Instead, a less stringent assumption is required, that the relative mutation rate remains proportional across all regions during tumor development, at least at the length scales (typically $\approx$ Mb) of the gained segments that are timed. Processes that lead to local hyper-mutation such as kataegis [8] can violate this assumption. However, such mutations typically have distinguishable properties [8, 9] and so can be filtered.

#### 4.3 Indels

Only SNVs are used to define mutation time. Indels are not included, even though they also accumulate over the course of tumor development. This is because the greater degree of alteration caused by indels compared to SNVs causes a systematic alignment bias. Fewer indel-containing reads will correctly align to the reference genome compared to SNVs, causing indel multiplicities to be systematically underestimated. Although computational methods have been developed to correct this alignment bias [10], the simplest approach is to define mutation time using SNVs only.

#### 4.4 The number of SNVs required to time a gain

A sufficient number of SNVs is required to time copy number gains with reasonable precision. This can be problematic for certain cancer types with low mutation rates [5]. More generally, it means that whole genome sequencing data is required for timing, as the number of SNVs measured by whole exome sequencing or targeted panels is typically too low to time all but the largest copy number gains.

#### 4.5 Infinite sites assumption

A recent study tested the validity of the infinite sites assumption across the Pan-Cancer Analysis of Whole Genomes (PCAWG) cohort [11]. Although sequential mutations to the same base pair could be identified in a number of tumors, they remain overall very rare in the context of the total SNV burden. The limited violations will cause a slight bias towards measuring later gains, due to the increase in SNVs with multiplicity greater than one.

#### 4.6 Subclonal copy number changes

Subclonal copy number events will lead to SNVs with fractional multiplicity states. Most SNV gain timing methods assume entirely clonal tumor copy number profiles and therefore only measure integer multiplicity states. This can lead to systematic biases in the measurement of gain timing. For example, a subclonal loss on a gained allele would lead to an underestimation in the number of SNVs with multiplicity >1 and therefore the timing measurement to be biased earlier.

It is possible to infer limited subclonal copy number from whole genome sequencing data [12]. The gain timing tool *MutationTimeR* used this information to correct the effect of subclonal copy number changes on gain timing measurements [6]. However, this required the ancestral relationships between subclonal copy number states to be assumed. Nevertheless, an assumption of clonality has been found to be reasonable for most tumor copy number events [7] and so any timing bias resulting from subclonal changes will generally be small. In this work, we use clonal copy number profiles to time copy number gains.

### 5 Problems with timing more complex gains

The equations used to relate SNV multiplicity proportions to timing are often under-determined. Consider a segment that undergoes three copy number gains to arrive at a 3+2 state (Fig. 6).

As before, we can write the equations for the number of mutations with each clonal multiplicity:

$$\begin{aligned} N_1 &= R(T_2 + 3T_3 + 5T_4) \\ N_2 &= R(T_1 + 2T_2 + T_3) \\ N_3 &= RT_1 \end{aligned} \tag{7}$$

Now there are five unknowns and four equations. This means this set of

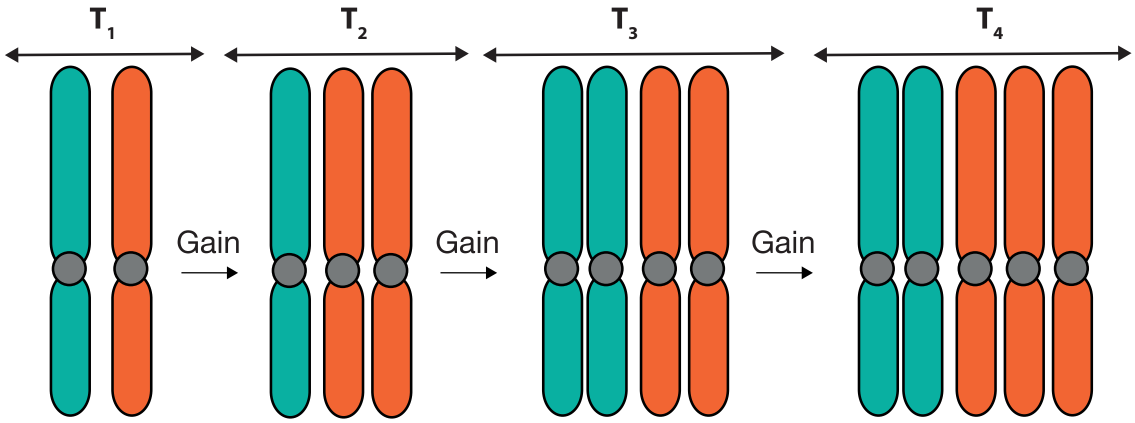

Figure 6: Schematic of a set of copy number gains leading to a 3+2 state.

equations is under-determined. However the timing of the first gain can be fully solved [6].

$$T_1 = \frac{5m_3}{m_1 + 2m_2 + 3m_3} \tag{8}$$

If we assume that the first and second gain occur at the same time so $T_2 = 0$, as could occur during a WGD (Fig. 7a) then the equations become fully determined. This is the assumed route history and equation used by the gain timing tool *MutationTimeR* to time the gains in a 3+2 state [6].

$$\begin{aligned} T_1 &= \text{WGD Timing} = \frac{5m_3}{m_1 + 2m_2 + 3m_3} \\ T_3 &= \frac{5(m_2 - m_3)}{m_1 + 2m_2 + 3m_3} \end{aligned} \tag{9}$$

This route posits that a single independent gain occurs after the WGD. However, it is reasonably plausible, with an additional loss event, that the gain could instead occur before the WGD (Fig. 7b) which leads to a separate set of equations for the timing of the independent gain and the WGD.

The routes displayed in Figure 7 are two potential histories that lead to a 3+2 copy number state. The route given in Figure 7a is the most parsimonious,

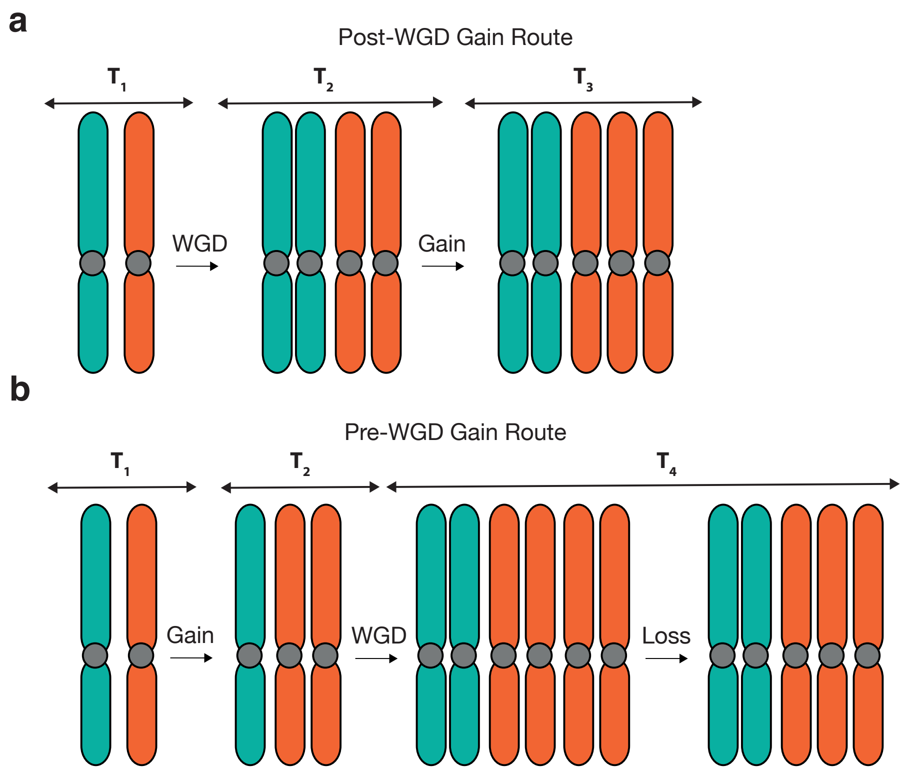

Figure 7: Schematic of two routes in a WGD tumor that lead to a 3+2 state. **a,** The most parsimonious route involving a single gain post-WGD **b,** A less parsimonious route with a pre-WGD gain and post-WGD loss.

with the fewest events, and so is the one that is often assumed to occur [6, 13]. However, the route in Figure 7b is also plausible and only requires a single loss, a common event in WGD tumors. This is not exhaustive, there are other routes that can lead to a 3+2 copy number state. In the next section, we develop a representation of all event histories that can lead to gained copy number states.

### 6 Representing routes as binary trees

A copy number gain will give rise to two alleles with a shared set of SNVs. This inheritance pattern can be represented in a binary tree structure. Greenman *et al.* [4] was the first to use tree structures to represent the gain history of a given segment. In this study, the structural variant and copy number information were used to jointly estimate a sequence of events that give rise to a given copy number profile. The time between each event was defined as a separate period. A tree was then constructed with a set of nodes for each time period corresponding to each allele. The directed edges between the nodes represented the inheritance patterns between the time periods. A pair of trees was used to represent the history of both parental alleles.

We developed a conceptually similar tree representation that allowed the possible route histories for a given copy number segment to be enumerated. In this representation, all leaf nodes represent the observed alleles and each ancestral node a copy number gain (Fig. 8). As with Greenman *et al.*, each copy number segment is represented by two trees, corresponding to both parental alleles. Gains that arise from a WGD are represented as a separate node class as they result in simultaneous gains across the tumor and therefore can be distinguished from independent copy number gains.

As this tree representation uses nodes to represent copy number events rather than time periods, each tree structure represents a distinct route history. As it generically represents copy number gain inheritance patterns, all possible route histories can be described using this tree structure. Therefore, this representation can be used to systematically enumerate the possible routes that lead to a given copy number state, without relying on structural variant data as in Greenman *et al.* [4].

We generate all possible binary trees using a simple recursive approach. We ensure that each tree structure is unique by only generating trees where the number of nodes on the right side of a bifurcation is greater than or equal to those on the left side. After generating all possible tree structures, we then enumerate all possible valid combinations of WGD nodes for the tree structure. Restricting to cases with no more than one WGD, only trees with at most a single WGD node in all paths from the initial to the leaf nodes are considered valid. Figure 9 gives the tree representation of all gain routes leading to a 4+2 copy number state under these constraints. While unlikely,

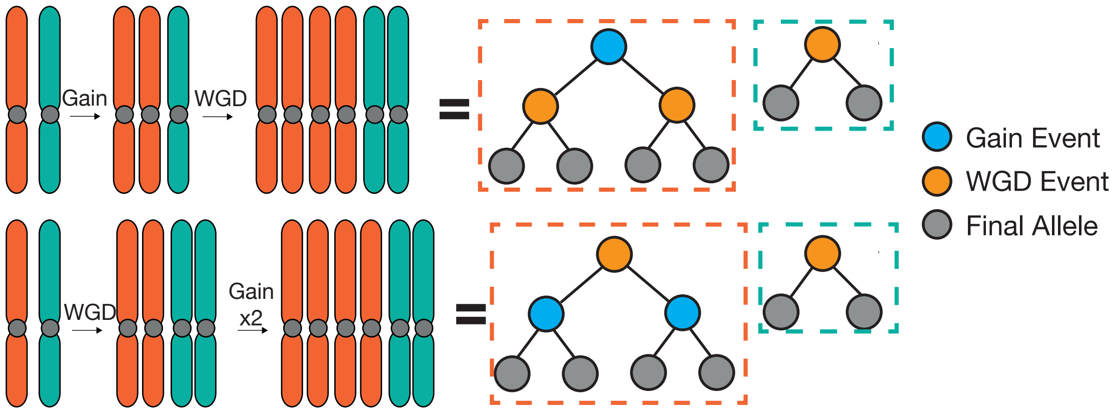

Figure 8: Examples of two routes to a 4+2 copy number state and their corresponding tree representation.

it is possible to have tree representations in WGD tumors that have no WGD nodes. This can occur when all additional alleles gained through the WGD are lost.

We used the tree representations to enumerate all possible routes across a range of copy number states in WGD tumors. The number of routes rises exponentially with increasing copy number. As an example, there are 8 possible routes for a 3+2 copy number state and 17,112 possible routes for a 10+4 copy number state.

Certain tree representations imply the occurrence of a copy number loss. In genome-duplicated tumors, trees with leaf nodes that do not inherit from a WGD node must have had a loss event. (Fig. 10). This is because a WGD causes all alleles to be duplicated. Therefore, any allele that does not share an inheritance with at least one other allele through a WGD must have lost the corresponding allele(s) that were gained during the WGD (Fig. 10).

By enumerating all possible tree structures, it is possible to identify all possible routes leading to a given copy number state. By possible routes, we refer to all routes that can potentially leave a trace in terms of shared SNVs between alleles. Gains that are subsequently lost cannot be timed. For example, the timing of the gain in a 1+1 state that arises through a gain and

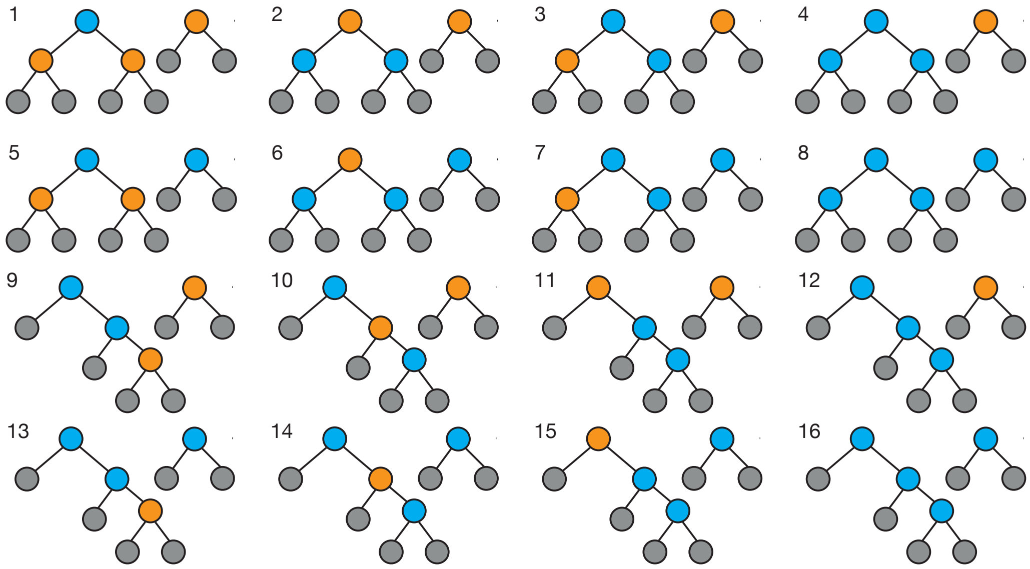

Figure 9: Tree representation of all possible gain routes detectable using SNV multiplicities that lead to a 4+2 copy number state, assuming no more than a single WGD. Losses are omitted from this representation for clarity.

then a subsequent loss cannot be measured through this technique (Fig. 11). The only exception to this are the losses of gained alleles that arise through a WGD, as these can be inferred from the presence of a WGD in the tumor as a whole.

### 7 Using the representations to time complex gains

The tree structures provide a useful basis to time the copy number gains in the routes they represent. They can be straightforwardly applied to infer the relationship between SNV multiplicity and timing similar to equations 1, 3 and 7. As an example, consider the route given in Figure 12. The equations for the number of SNVs with each multiplicity state can be written as follows:

$$\begin{aligned} N_1 &= R(T_2 + T_4 + T_5 + T_7 + T_8) \\ N_2 &= R(T_3 + T_6) \\ N_3 &= RT_1 \end{aligned} \tag{10}$$

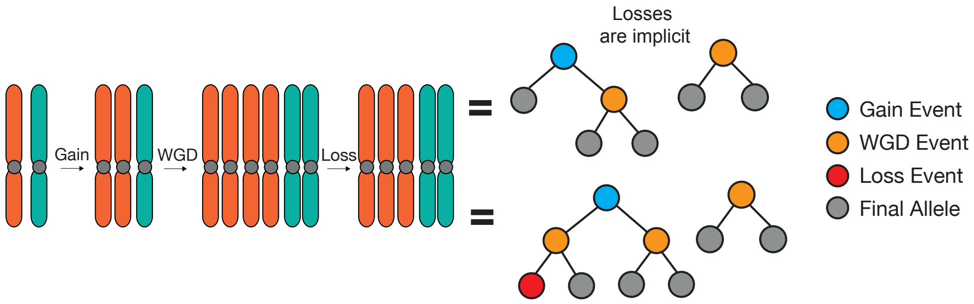

Figure 10: Schematic of a set of copy number events leading to a 3+2 copy number state and the associated tree representation. Although a copy number loss is necessarily implied by the event history, it is omitted for clarity.

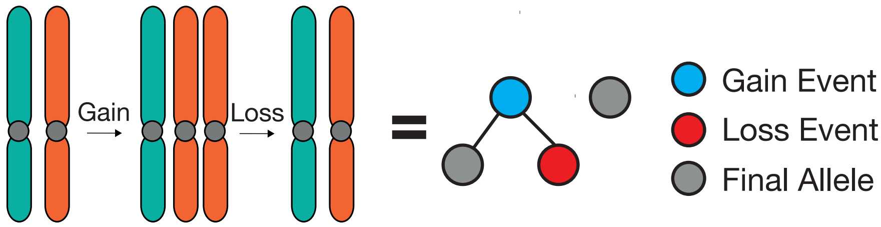

Figure 11: Schematic of a route history and its corresponding tree representation where a 1+1 state gains an allele which is then lost. The gain in this route history cannot be timed, as no record of the gain remains in the SNV data.

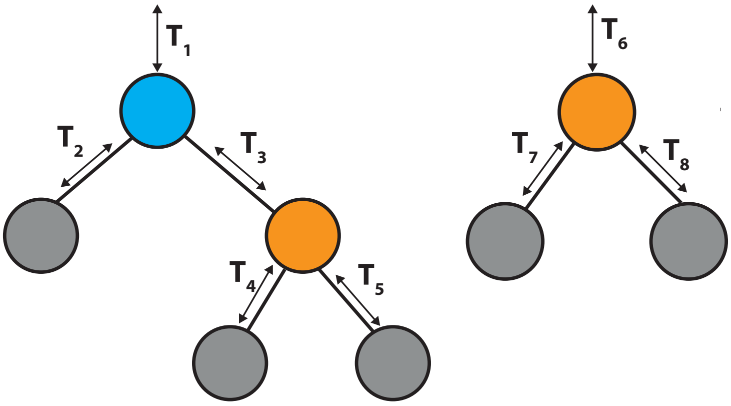

Figure 12: A tree diagram representing a route leading to a 3+2 copy number state, annotated with the time periods between copy number events.

There are two separate constraints that apply to these multiplicity equations. Due to the definition of mutation time, the sum of the time periods of all paths from each root to the leaves must equal one. Additionally, the WGD is simultaneous, and therefore there is an additional constraint that $T_1 + T_3 = T_6 = T_{WGD}$. Using these constraints and some relabelling of time periods, we can arrive at the same equations as in equation 7.

This can be applied to any copy number tree. The relationship between gain timing and SNV multiplicity and timing as well as the two timing constraints can be represented in two separate binary matrices $\mathbf{A_M}$ and $\mathbf{A_C}$.

$$R\mathbf{A}_M\mathbf{T} = \mathbf{N} \tag{11}$$

$$\mathbf{A}_C\mathbf{T} = \mathbf{C} \tag{12}$$

Here, $\mathbf{N}$, $\mathbf{C}$, and $\mathbf{T}$ are the vectors encoding the SNV multiplicities, gain timing constraints and time periods, respectively. For example, the two matrices for the previous route (Fig. 12) are given by:

$$R\begin{pmatrix} 0 & 1 & 0 & 1 & 1 & 0 & 1 & 1 \\ 0 & 0 & 1 & 0 & 0 & 1 & 0 & 0 \\ 1 & 0 & 0 & 0 & 0 & 0 & 0 & 0 \end{pmatrix}\begin{pmatrix} T_1 \\ T_2 \\ T_3 \\ T_4 \\ T_5 \\ T_6 \\ T_7 \\ T_8 \end{pmatrix} = \begin{pmatrix} N_1 \\ N_2 \\ N_3 \end{pmatrix} \tag{13}$$

$$
\begin{pmatrix}
1 & 0 & 1 & 1 & 0 & 0 & 0 & 0 \\
1 & 0 & 1 & 0 & 1 & 0 & 0 & 0 \\
1 & 1 & 0 & 0 & 0 & 0 & 0 & 0 \\
0 & 0 & 0 & 0 & 0 & 1 & 1 & 0 \\
0 & 0 & 0 & 0 & 0 & 1 & 0 & 1 \\
1 & 0 & 1 & 0 & 0 & 0 & 0 & 0 \\
0 & 0 & 0 & 0 & 0 & 1 & 0 & 0
\end{pmatrix}
\begin{pmatrix}
T_1 \\ T_2 \\ T_3 \\ T_4 \\ T_5 \\ T_6 \\ T_7 \\ T_8
\end{pmatrix}
=
\begin{pmatrix}
1 \\ 1 \\ 1 \\ 1 \\ 1 \\ T_{WGD} \\ T_{WGD}
\end{pmatrix}
\tag{14}
$$

It is not necessary to invert these general matrix equations directly to find a relationship between multiplicity proportions and gain timing for the route. Indeed, if the equation is underdetermined, a direct solution can be somewhat complex to interpret and use. Instead, we can avoid this by using equations 11 and 12 directly.

It is straightforward to use computational methods to find a single arbitrary valid solution $\mathbf{T_S}$ to equation 12. Then, by sampling vectors $\mathbf{T_N}$ from the null space of $\mathbf{A}_C$ such that $\mathbf{A}_C\mathbf{T_N} = \mathbf{0}$ we can sample from the full set of solutions to equation 12 as any $\mathbf{T_S} + \mathbf{T_N}$ will be a solution. We can then use equation 11 to find the multiplicity proportions that correspond to each sampled timing vector.

### 8 Multiplicity spaces

So far, we have outlined a general method to identify all timeable event histories that can give rise to a given complex copy number state, accounting for a potential WGD timing constraint. We then showed how these representations can be used to sample possible gain timing values and associated multiplicity proportions for each route.

We now consider the space of possible multiplicity proportions for different routes across a number of copy number states. By definition, all possible clonal multiplicity proportions for a segment with a major copy number of $n_A$

will exist in the standard simplex of dimension $n_A - 1$. In this section, we show that within the simplex for a copy number state, the space of possible multiplicity proportions is often distinct for different routes. Therefore, this permits different route histories to be distinguished using SNV information.

#### 8.1 Multiplicities spanned by a 3+0 copy number state

As the simplest example of a complex gained route, consider a 3+0 copy number state. In a WGD tumor, there are three possible route histories (Fig. 13). We arbitrarily label these routes A, B and C. In route A, the independent gain occurs before the WGD, whereas it occurs after the WGD in route B. Finally, route C has two independent gains, meaning that all extra alleles gained through the WGD have been lost.

All three routes have the same overall tree structure. This structure has a fully determined relationship between SNV multiplicity and gain timing. Therefore, both the first and second gain timing can be uniquely determined from SNV multiplicity. Nevertheless, the WGD constraint restricts the possible multiplicity space. This allows us to make inferences on parsimony in copy number solution evolution. Route B is the most parsimonious solution for this state in a WGD tumor. This is because routes A and C imply additional losses as they contain final alleles that do not inherit from a WGD node.

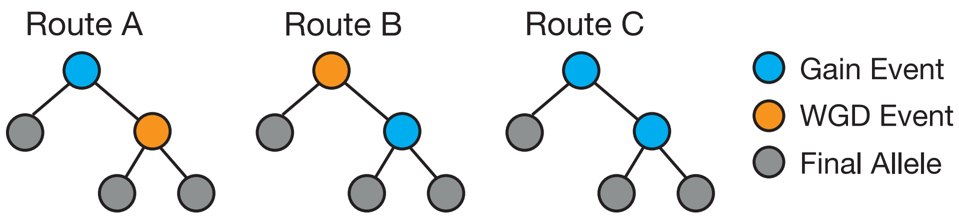

Figure 13: Tree representations of the three possible routes that can lead to a 3+0 state in a WGD tumor.

For a given WGD timing, routes A and B have a single degree of freedom. They therefore have a one-dimensional range of possible multiplicity states.

In route A, the independent gain primarily dictates the relative proportion of multiplicity two and three SNVs, whereas in route B it is the relative proportion of multiplicities one and two. We sample the possible multiplicity states using the Markov Chain Monte Carlo (MCMC)-based hit and run algorithm (Section 10.3, Fig. 14).

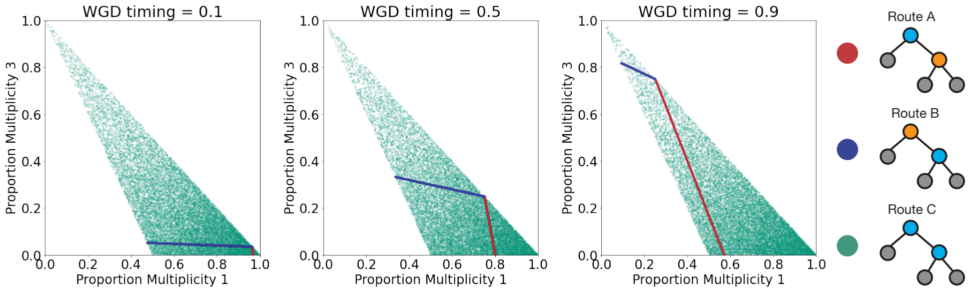

Figure 14: Samples of possible multiplicity states spanned by the three routes that can lead to a 3+0 state in a WGD tumor. Shown for three example WGD timings.

The set of possible multiplicity proportions for routes A and B change according to the WGD timing. Later genome-duplications increase the potential range of gain timing for route A and thus widen the span of possible multiplicity states. Similarly, the range of possible multiplicity states for route B is reduced with later WGD timing.

Route C has two independent gains and thus has two degrees of freedom in the multiplicity states. Because neither gain arises from the WGD, the space of possible states is the same regardless of WGD timing. As the timing of either independent gain in route C could be equal to a given WGD timing, the possible multiplicity states for routes A and B are both subsets of route C.

Nevertheless, given a multiplicity state that is consistent with route A or B, it remains more likely that it arose from these routes rather than route C, given that route C would require a loss of the WGD gained alleles as well as an independent gain at the same time as the WGD. This can be encoded in the increased density of multiplicity samples from routes A and B, if all routes are given an equal number of samples.

#### 8.2 Multiplicities spanned by a 4+0 copy number state

Although the timing of the WGD constraint will vary, there is only one overall possible tree structure for major copy number three gains. This means that when combined across all possible WGD timings, all routes for these major copy number states span the same entire set of multiplicity states.

This is no longer the case for higher major copy number states, as there are multiple possible tree structures. Consider a 4+0 copy number state. In this case, there are two separate tree structures, regardless of the WGD timing constraint (Fig. 15). The two structures diverge based on the third and final gain. If it occurs on one of the alleles created in the second gain, then the route is called unbalanced, otherwise it is called balanced.

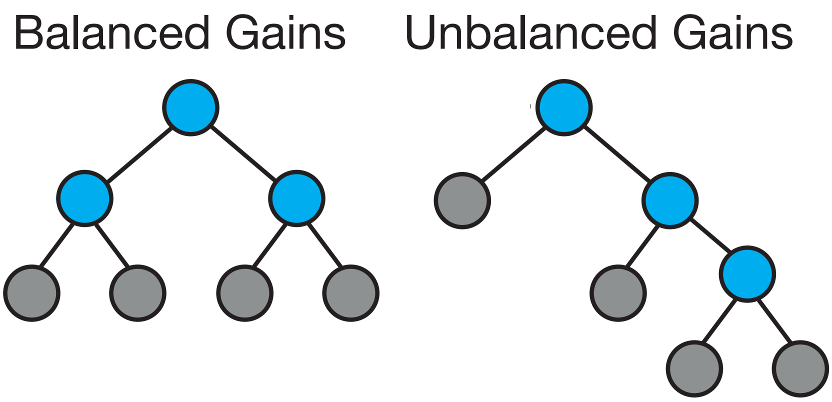

Figure 15: The two possible tree structures for a 4+0 state.

The relationship between multiplicity and gain timing is fully determined for the unbalanced gain structure and under-determined for the balanced gain structure. Without rigorous justification, this can be seen by considering the number of descendants of each gain node. The three gains in the unbalanced case each give rise to a different number of alleles and thus each relates to a different multiplicity state. For the balanced structure, the second and third gains both lead to multiplicity two SNVs and therefore, have under-determined timing. This can be resolved if the two secondary gains are assumed to occur simultaneously.

Therefore, there is ambiguity when timing major copy number four gains, both from the two possible tree structures but also from the under-determined timing of the secondary gains in the balanced gains routes. As discussed earlier, this meant that the timing of major copy number four gains was not fully considered in the three methods used by Gerstung *et al.* [6].

However, the two routes span different possible multiplicity spaces. We therefore realized that the routes could be distinguished for a large number of possible gain timings. In the most obvious case, any segment with any number of SNVs with multiplicity three must arise through the unbalanced gains route (Fig. 16). In this case, to visualize the boundaries between the different multiplicity proportions, we systematically enumerate all possible gain timings up to a small precision, rather than sampling using the hit and run approach.

But even when the unbalanced route is restricted to cases where the third gain immediately follows the second, such that there are no multiplicity three SNVs, there are still differences among the two possible multiplicity spaces (Fig. 17). In these circumstances, the unbalanced gain route only spans a subset of the full probability simplex across multiplicities one, two and four. In contrast, the balanced gains route spans the entire possible set of multiplicity one, two and four proportions. Therefore, for a large region of this space, SNV multiplicity proportions can be unambiguously assigned to the balanced gain route.

This sampling approach also allows for the ambiguity in the timing of the secondary gains in the balanced gain route to be quantified, although not resolved. By collecting samples in a sufficiently local region around a given multiplicity state, the set of possible gain timings that correspond to the multiplicity state can be sampled. This collects any useful timing information about the secondary gains from the multiplicity proportions, even though its precise timing is under-determined. This approach is valid for any under-determined route history.

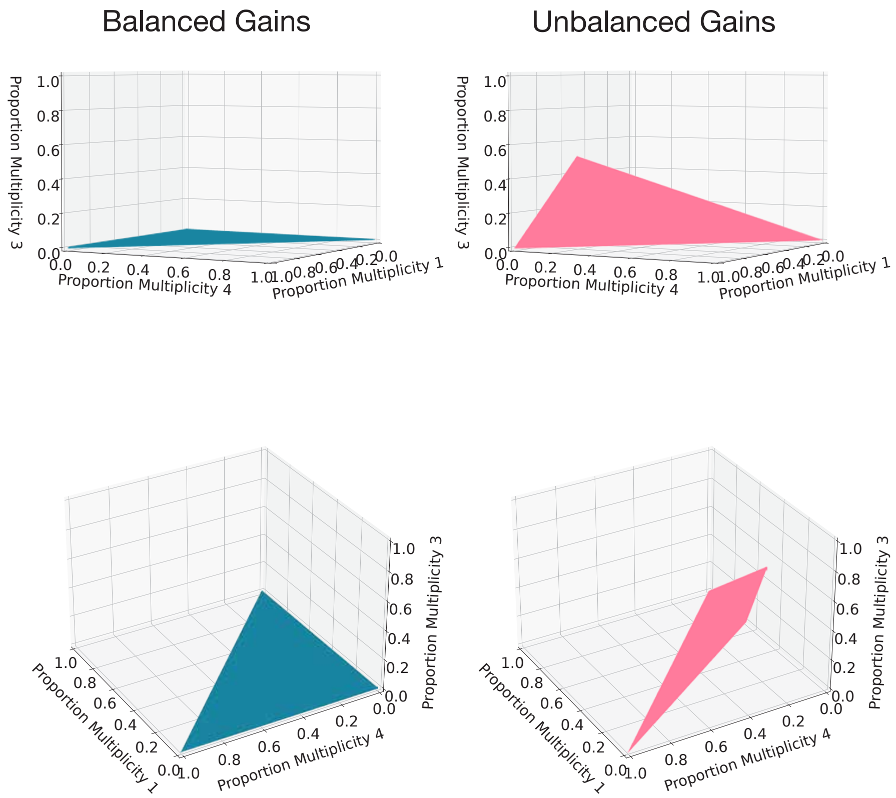

Figure 16: The set of possible multiplicity states corresponding to balanced and unbalanced gains for a 4+0 copy number state.

#### 8.3 Multiplicity spanned by more complex states

For more complex copy number states, the range of possible gain timings and multiplicities span simplexes of higher dimensions and are impossible to directly visualize. As discussed earlier, the number of possible routes also rises exponentially with increasing copy number.

We display an example of this complexity in Fig. 18. Here, we show the first two principal components of the 5 dimensional space spanned by samples from the possible multiplicity proportions for all 34 routes that lead to a 5+2 copy number state, given an arbitrary WGD timing of 0.4. 90.3% of the total variance can be attributed to these first two components. There is significant overlap between the states, but the constraints for each route mean that

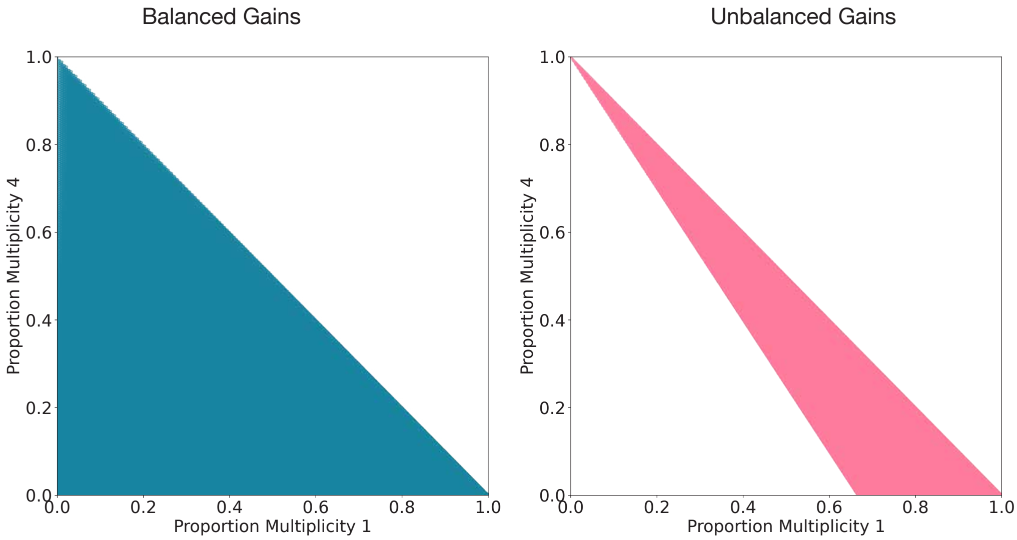

Figure 17: The set of possible multiplicity states corresponding to balanced and unbalanced gains for a 4+0 copy number state. The balanced case is restricted to gain timings that lead to no multiplicity three SNVs, to match the space spanned by the balanced case.

different routes have different areas and have density over multiplicity space when sampled with uniform distribution over gain timing.

We then sought to more formally assess whether the different multiplicity states spanned for each route could be distinguished across copy number states. We considered the routes leading to the 20 most common copy number states. We sampled gain timings and associated multiplicity proportions for each route using the hit and run approach (Section 10.3). By calculating the fraction of nearest neighbors for a given multiplicity proportion that arose from the same route (Section 11), we could measure how well separated the multiplicity spaces were for each complex copy number state.

We found that an average of 57.8% of nearest neighbors across all routes matched the correct route (Fig. 19). As expected, this varied significantly between copy number states. The number of nearest neighbors with the same route decreases as the number of possible routes increases, from 99.0% for the three routes in the 3+0 state to 21.2% for the 136 routes for

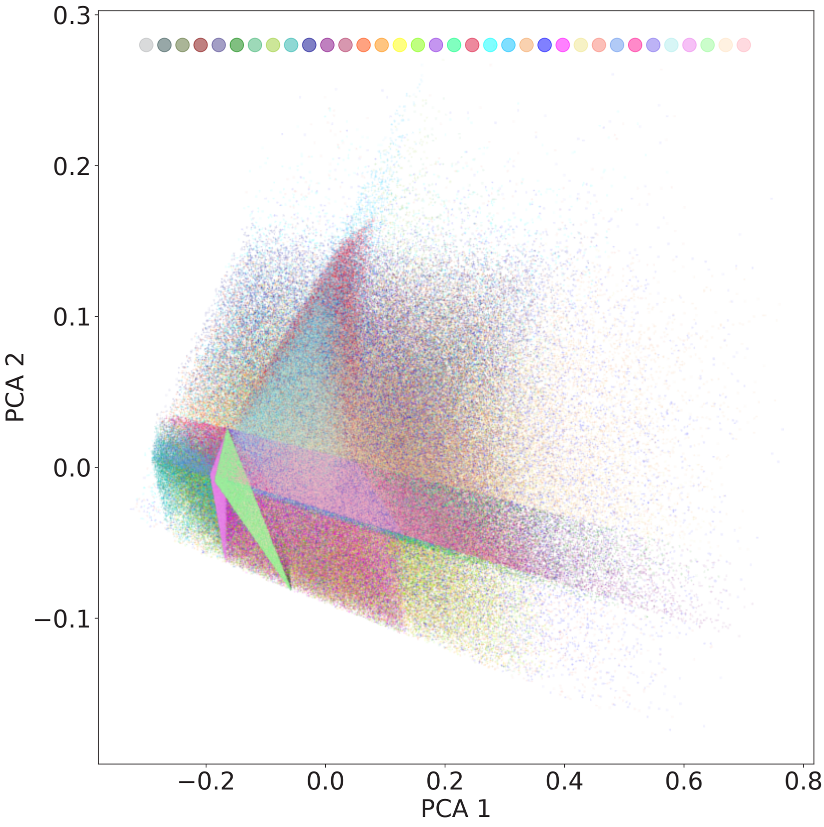

Figure 18: 20,000 samples from the set of possible multiplicity states for all 34 possible routes leading a 5+2 copy number state given a WGD timing of 0.4.

5+4 copy number gains.

Notably, this relationship is not monotonic with regard to the number of routes for a copy number state. For example, the nearest neighbor proportion was on average 26.6% across the 226 routes in the 7+2 state compared to 21.2% for the 136 routes leading to 5+4. This can be explained by the minor copy number.

While a higher major copy number increases the number of routes, it also increases the dimensions of the multiplicity space. In contrast, a higher minor copy number still increases the number of routes without expanding the dimension of the multiplicity space. For all states, the proportion of nearest neighbors is significantly higher than would be expected under the random distribution of multiplicities proportions for each route across the full simplex.

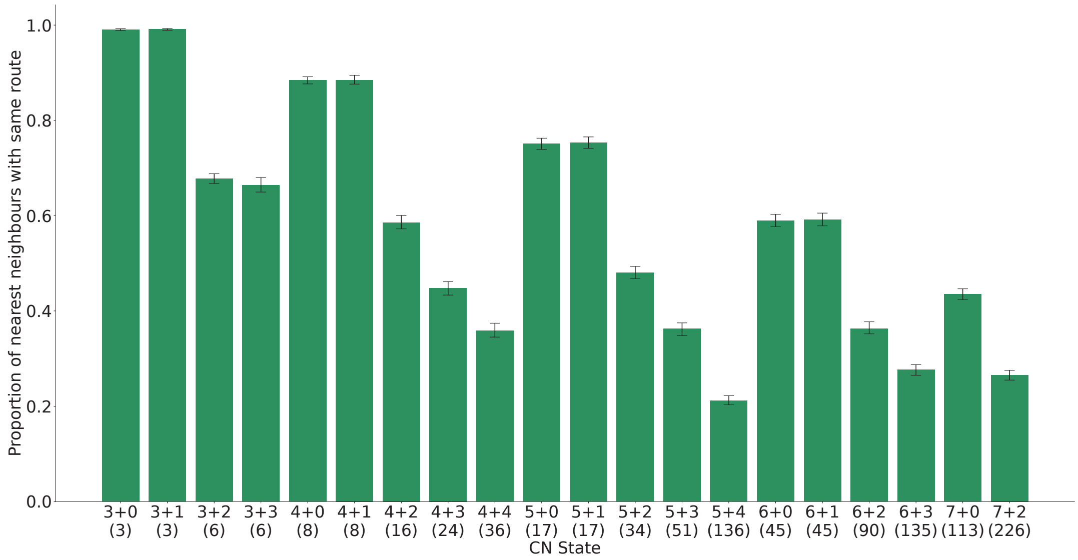

Figure 19: The proportion of nearest neighbors that arise from the same route for a number of complex copy number states. The proportions across a range of WGD timings. The number of possible routes for each state is given in brackets.

It should be emphasized that the separation between the multiplicity states for different routes is partly a result of relative density. Any independent gain can match the timing of a given WGD. Therefore, routes with all independent gains will always be able to match routes with the same tree structure and WGD constraints.

However, as each independent gain represents a new degree of freedom, only a small proportion of samples will have independent gains that match the WGD timing. This can be thought of as a weak parsimony constraint. If a given multiplicity proportion can be formed through a route with WGD constraints, the local density of multiplicity proportions from such a route will be much higher than from the corresponding route with all independent gains.

### 9 GRITIC

These results suggest that many complex routes and their conditional gain timing can be distinguished through multiplicity proportions. This forms the basis of GRITIC, the approach we developed to time complex gains. The main details of the GRITIC method are described in the following sections and are briefly summarized here.

For each segment, we apply the hit and run sampling procedure to obtain the set of possible gain timings and associated multiplicity proportion for each route. The likelihood of each sampled multiplicity proportion is evaluated given the SNV read counts using a binomial mixture model. The likelihoods are used to calculate the conditional gain timing distributions for each route. The marginal likelihood of each route is then computed by averaging the likelihood over all sampled multiplicity proportions.

GRITIC uses the simultaneous occurrence of a WGD as a constraint in WGD tumors to improve fitting. The WGD timing distribution is calculated by considering major copy number two segments, as we assume the majority of gains with this major copy number arise through a WGD. Each segment with a major copy number of two is timed independently. The point that corresponds to the maximum overlap of their timing credible intervals is then computed. The timing of the WGD is estimated by jointly timing a single gain across major copy number two segments with timing credible intervals that overlap with the point of maximum overlap.

After the WGD is timed, the gains leading to more complex segments are timed using the procedure above. The WGD distribution is used as a constraint when sampling possible gain timings for a given route. In the case of non-WGD tumors, all routes are sampled without a WGD.

### 10 GRITIC method

#### 10.1 Overview

The principles outlined in the previous section are used by GRITIC to time the copy number gains. A simple high-level outline of the GRITIC procedure for a genome-duplicated sample is:

1. Each segment with a major copy number of two is timed independently.
2. The point in mutation time that overlaps with the greatest fraction of gain timing of major copy number two segments is calculated.
3. The timing of the WGD is estimated by jointly timing the gain across all major copy number two segments with a timing credible interval that overlapped with the point of maximum overlap.

Then, for each timeable complex segment, we do the following.

1. The tree representations for each possible route are enumerated.
2. An MCMC procedure is carried out for each route to sample the possible set of gain timings given the WGD timing distribution.
3. The likelihood of each sampled gain timing is evaluated from its corresponding multiplicity proportions and SNV read count data.
4. The probability of each route is calculated from the average likelihood of its gain timing samples, divided by the sum of the average likelihoods for every possible route.
5. This probability is then stored along with the distributions of gain timing conditioned on each route and the SNV multiplicity proportions.

The above procedure is also followed for all gained segments with sufficient SNVs in non-WGD tumors, though only routes without WGD constraints are considered. More details about each of the sections of GRITIC are given in the subsections below.

#### 10.2 Generating trees

##### 10.2.1 Generating tree structures

The binary tree structures are generated with a simple recursive approach. To ensure that each distinguishable gain structure corresponds to a single tree, only trees with at least half the leaves on the right side of a bifurcation are generated.

##### 10.2.2 Generating WGD nodes

We use a brute-force approach to generate all possible combinations of WGD nodes for each given tree structure. All possible combinations of all lengths of the tree's internal nodes are checked to see if it is a valid set of WGD nodes. A set of WGD nodes is valid if no WGD node inherits from another.

Then, for each valid set of WGD nodes for each tree structure, we compute the combined hash of the tree structure and WGD node status using the Weisfeiler Lehman graph hash method [14] as implemented in `networkx` [15]. As each tree with the same hash has an equivalent structure, we only select one tree with each hash. In theory, these structures could be much more efficiently generated with a recursive algorithm instead of brute force, but we find that this approach works sufficiently quickly for the tree structures that we consider.

#### 10.3 Sampling valid timing states

We use an MCMC approach to sample uniformly from the set of timing vectors $\mathbf{T}$ that are solutions to equation 12. These can then be converted to multiplicities using equation 11. We first find a single valid solution to equation 12, using a non-negative least squares solver to enforce that $\mathbf{T} \geq \mathbf{0}$.

From this initial solution, we compute a chain of valid solutions using the hit and run algorithm [16]. At each step, a random direction is chosen from the null space of $A_C$. The next position in the chain is found by moving a random distance in this direction with a distance uniformly sampled between the limits that maintain $\mathbf{T} \geq \mathbf{0}$.

This procedure was simplified in the case where the null space of $A_C$ had rank one, corresponding to routes with a single independent gain. For computational efficiency, the states were sampled with uniform spacing along the direction of the null space, with the limits given as before. The set of sampled states was then shuffled.

In GRITIC, the chain is reset and a new WGD timing constraint is obtained after 500 samples are taken. We found that the initial solution is often in a region of the space that is difficult for the sampler to move away from. We therefore allow a very large number of initial steps, until the sampler moves an initial distance of 0.001 in timing space from its initial position. We permit up to 50,000 steps for the initial move away from the start position, before then applying a further 25 burn in steps.

#### 10.4 Subclonal multiplicity sampling

The sampled clonal multiplicities must be weighted to account for subclonal mutations in the segment. For each clonal multiplicity sample, we randomly sample the relative fraction of clonal SNVs and those in each subclonal cluster. The sampled clonal multiplicity proportions obtained from the gain timing are then weighted by the clonal SNV proportion and combined with the proportion of multiplicities in each subclonal cluster. This gives a full multiplicity proportion where the likelihood can be evaluated using the SNV read counts.

To aid convergence, we use the number of SNVs across the tumor contained in the clonal peak and each of the subclones as a prior for sampling the proportion of subclonal SNVs for each segment. This is a sample-wide estimate, determined by a subclonal reconstruction algorithm and given as input to GRITIC. As the subclonal proportions for a given segment can differ substantially from the overall sample-wide subclonal proportion, we keep this prior relatively weak. The clonal and subclonal proportions for each segment are sampled from a Dirichlet distribution $Dir(\alpha)$ where $\alpha$ is defined as:

$$\alpha_i = \frac{S_i}{\sum_{j=1}^{N} S_j} + 1 \tag{15}$$

Here $S$ is the vector of the number of SNVs determined by the subclonal deconvolution algorithm to be part of the $N$ clonal and subclonal clusters.

Each additional subclonal cluster increases the dimension of the sampled space and thus the number of samples required to adequately cover it. However, the proportion of SNVs in each subclone is not relevant for the gain timing, we are only interested in the proportion of clonal SNVs. Therefore, to aid convergence, we limit each tumor to two subclones.

For any tumor with more than two subclones, the subclone with the largest cancer cell fraction (CCF) is maintained and the smaller subclones are merged by taking the average CCF of the subclones weighted by the number of SNVs they contain. This improves inference speed on tumors with a large number of subclones without impacting the accuracy of estimating the proportion of clonal SNVs.

#### 10.5 Ensuring sufficient density of samples

Increasing the number of sampled timing states improves the precision of the timing measurements and route probabilities. To ensure the most efficient use of computational resources, we implemented a process that sampled timing states up to a sufficient density. To account for the density of the full sampling process, we computed the density in a space formed by concatenating the timing and subclone proportion vectors.

The density of sampled states was approximated using a nearest neighbors approach. As a proxy for density, we computed the proportion of a random subset of states that had at least one neighboring sample within a Euclidean distance of 0.05. We defined the space to have reached a sufficient sampling density when this density measure reached 90%.

This density computation is periodically computed during the sampling process. Once the threshold is met, the sampling is stopped. Due to the curse of dimensionality, this density threshold can take a very long time to be reached for high dimension space. Therefore, we use a secondary backup threshold that stops sampling once 500,000 samples have been reached.

In practice, this means that some copy number states finish with a low density of sampled timing vectors. For example, the average final density across all 7+0 routes is 0.316 but is as low as 0.0006 for some routes. Nevertheless, it is clear from the tests using simulated data that the current densities permit the gain timing in complex segments to be measured with sufficient precision. With further work, a more sophisticated sampling procedure could be implemented that further optimizes the trade off between run time and precision.

Currently, GRITIC samples uniformly across all possible timing states to compute the marginal likelihood for each route. The gain timings conditioned on each route are obtained from resampling these states according to their multiplicity likelihood. An accurate timing distribution could be obtained with substantially fewer samples if the posterior gain timing for each route was sampled directly, perhaps using a Metropolis-Hastings approach. However, computing the marginal likelihood of each route directly from the posterior gain timings is significantly more involved [17, 18] and we find our current approach works well in practice.

#### 10.6 Evaluating multiplicity likelihoods

The likelihood of a given proportion of SNVs with each possible multiplicity $m_i$ for a given copy number segment can be inferred from the number of mutated and reference reads for the SNVs in the segment.

Specifically, we consider the variant allele frequency (VAF) for each SNV.

This is the proportion of alternative reads for the SNV regions. The expected VAF for a mutation with multiplicity $M$ in an autosomal chromosome can be written as:

$$VAF_M = \frac{\rho f M}{n^T \rho + 2(1-\rho)} \tag{16}$$

Here $\rho$ is defined as the proportion of tumor cells in the biopsy, known as the purity. $f$ is the proportion of tumor cells that contain the SNV, the CCF. The total tumor copy number for the segment is given by $n^T$. The $2(1-\rho)$ term in the denominator arises as non-cancerous cells have two copies of each autosomal region. To avoid issues with a varying baseline number of chromosomes, only autosomes are analysed in the current version of GRITIC. The equation uses the constant mutation multiplicity assumption [19], where it is assumed that each SNV has the same multiplicity in all of the cells that contain the SNV.

We further assume that all subclonal SNVs have a multiplicity of one. Then, for SNVs in a subclonal cluster with CCF $s_i$, their expected VAF is given by:

$$VAF_M = \frac{\rho s_i}{n^T \rho + 2(1-\rho)} \tag{17}$$

To simplify, as GRITIC only uses clonal SNVs for timing purposes, we include the CCF in our definition of multiplicity. Each subclonal cluster is therefore considered as a distinct multiplicity state. With the infinite sites assumption, an SNV can have a maximum multiplicity of the major copy number $n^A$ of the segment.

For a sample with $N$ subclonal clusters $s_1, s_2, ..., s_N$ and a genomic region with major copy number $n^A$ the set $Q$ of possible multiplicities can be written $Q = \{s_1, s_2, .., s_n, 1, ..., n^A\}$.

We use a binomial distribution to model the number of alternative reads obtained for each SNV. The probability that a given mutation has multiplicity $M$ is then proportional to:

$$P(M|n^T, n^A, R^M, R^T) \propto Bin(R^T, R^M, VAF_M) \tag{18}$$

Here $R^T$ is the total number of reads at the mutated position, $R^M$ the number of reads that contain the SNV and $Bin$ the binomial probability mass function. The likelihood that a segment has a combination of SNV multiplicity proportion $\mathbf{m} = [m_{s_1}, m_{s_2}, ..., m_{s_n}, m_1, ..., m_{n^A}]$ is given by a binomial mixture model multiplying over all SNVs in the segment $j$. This is computed in log space for numerical stability.

$$P(\mathbf{m}) \propto \prod_j \sum_{i \in Q} m_i P(i|n^T, n^A, R_j^M, R_j^T) \tag{19}$$

The quantity required for timing is the proportion of clonal SNVs with multiplicity $i$, $m_i^c$. This can be found by dividing $m_i$ by the sum of all the multiplicity proportions corresponding to clonal SNVs. Throughout the text, unless otherwise stated, we use $m_i$ to refer to $m_i^c$ for notational convenience.

#### 10.7 Power correction for multiplicity likelihood

The higher the multiplicity of a given SNV, the greater the fraction of mutated reads and thus the higher the likelihood of the SNV being detected. This can lead to an overestimate of the proportion of SNVs with high multiplicities. This bias will be particularly apparent in tumors with low purity or depth of sequencing. We therefore applied a power to detect correction to each multiplicity state.

We approximate the conditions where a mutation would be detected by assuming that an SNV will only be detected if and only if three mutant reads are detected. For each clonal and subclonal multiplicity state $M$, we estimate the proportion of SNVs, $P_M$ that would be missed by this threshold.

$$P_M = \frac{1}{N} \sum_i P(X_i \leq 2 | n_i^T, M) = \frac{1}{N} \sum_i Bin_{CDF}(2, n_i^t, R_i^T, VAF_M) \quad (20)$$

Here $X_i$ is the number of mutated reads, $n_i^T$ the total copy number and $R_i^T$ the sequencing depth for the $i^{th}$ out of $N$ SNVs in the segment, respectively. $Bin_{CDF}$ represents the binomial cumulative distribution function. Then, for each multiplicity proportion $\mathbf{m}$ corresponding to a set of gain timings, we calculate the corrected proportion $\mathbf{m}'$ that would be expected to be observed given the power to detect.

$$m_i' = \frac{P_i m_i}{\sum_j P_j m_j} \quad (21)$$

We calculate the likelihood of $\mathbf{m}'$ using the equations in the previous section to give a power-corrected estimate of the likelihood of $\mathbf{m}$.

#### 10.8 Counting the number of events

We calculate the total number of events that correspond to each route such that a high number of events can be penalized to provide a lower bound on non-parsimony. Calculating the number of gain events is trivial and only dependent on the tree structure. It is simply the sum of all independent gain nodes in a given tumor plus an additional event corresponding to the WGD if the tumor is classified as such.

Identifying the number of losses for a route is more involved. As discussed earlier, losses are implicit in the tree representation and depend on the relative timing of gains and the WGD as well as the overall structure of the tree. A loss event must have occurred between all connections of independent gain nodes with a gain timing that intersects the WGD timing (Fig. 20). All final nodes, those that correspond to the observed alleles, have a timing of one. A minor copy number of zero or one additionally implies a pre- or post-WGD loss respectively (Fig. 20).

To estimate the number of events for a given route, we take the average the number of events implied in one hundred independent samples from the posterior gain timing distribution.

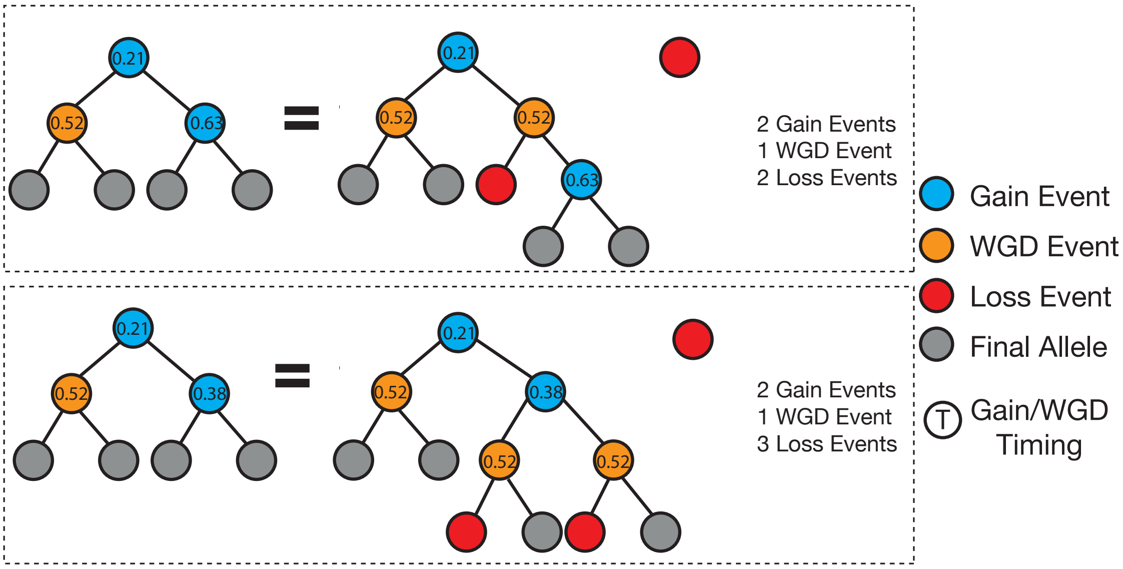

Figure 20: Example of the different number of loss events implied by different gain timings for a particular route leading to a 4+0 state given a WGD timing.

#### 11 Nearest neighbor evaluation methods

The following describes the methods for the nearest neighbor evaluation processed detailed in these supplementary methods. For each route that can lead to the 20 most common complex copy number states, we ran a chain that sampled 20,000 valid timing vectors using the hit and run approach. To assess convergence, we partitioned this data into two halves, the first and last 9,000 samples. Therefore, 18,000 samples were included for each copy number state. This was computed separately for 11 different WGD timing values between 0.01 and 0.99. As the WGD timing was fixed rather than sampled from a distribution, the chain was run continually for 20,000 steps instead of being periodically reset, as in GRITIC.

Then as before, each sampled timing state was converted into multiplicity states using equation 11. For each copy number state and WGD timing, we took 1,000 multiplicity states at random and collected their 10 nearest neighbors from the remaining multiplicity states. We assessed the proportion of these nearest neighbors that were from the same copy number route.

We validated that the convergence of the chain did not confound the estimate of nearest neighbors. For each multiplicity state under consideration, we computed the proportion of nearest neighbors with the same route that originated from the same half of the chain as the multiplicity state. Aggregated across all timing states, this proportion was very close to 0.5 for all copy number states (Fig. 21). This suggests that the copy number states had been adequately sampled and the nearest neighbors with the correct route weren't just multiplicity proportions that were sampled in close succession.

It is worth noting that routes with a single degree of freedom have a different sampling procedure that will bias this estimate. As discussed earlier, the single degree of freedom is sampled with uniform spacing between the valid limits before being shuffled. The shuffling of the obtained states would mean a nearest neighbor proportion of 0.5 would be expected. However, 1,010 out of the possible 1,019 routes across all of the evaluated copy number states have more than one degree of freedom.

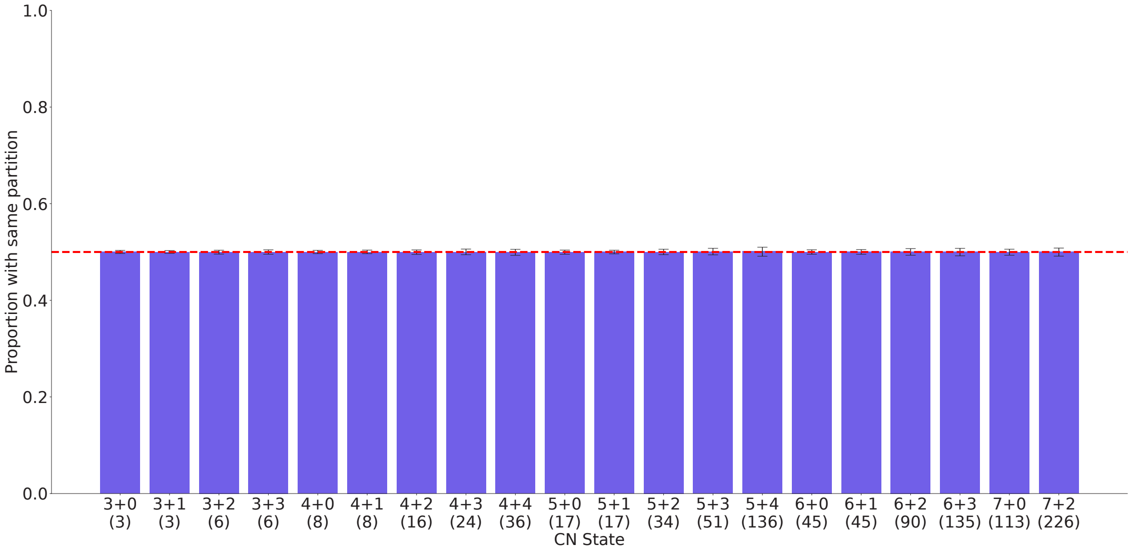

Figure 21: The proportion of nearest neighbors with the same route that arise from the same half of the multiplicity sampled chain as the multiplicity state under consideration. The dashed line indicates the 50% proportion that would be expected under convergence. The number of possible routes for each copy number state is given in brackets.
